# Organ geometry channels cell fate in the Arabidopsis ovule primordium

**DOI:** 10.1101/2020.07.30.226670

**Authors:** Elvira Hernandez-Lagana, Gabriella Mosca, Ethel Mendocilla Sato, Nuno Pires, Anja Frey, Alejandro Giraldo-Fonseca, Ueli Grossniklaus, Olivier Hamant, Christophe Godin, Arezki Boudaoud, Daniel Grimanelli, Daphné Autran, Célia Baroux

## Abstract

In multicellular organisms, sexual reproduction requires the separation of the germline from the soma. In flowering plants, the first cells of the germline, so-called spore mother cells (SMCs), differentiate as the reproductive organs form. Here, we explored how organ growth influences and contributes to SMC differentiation. We generated a collection of 92 annotated 3D images capturing ovule primordium ontogeny at cellular resolution in Arabidopsis. We identified a spatio-temporal pattern of cell divisions that acts in a domain-specific manner as the primordium forms, which is coupled with the emergence of a single SMC. Using tissue growth models, we uncovered plausible morphogenetic principles involving a spatially confined growth signal, differential mechanical properties, and cell growth anisotropy. Our analysis also reveals that SMC characteristics first arise in more than one cell but SMC fate becomes progressively restricted to a single cell during organ growth. Altered primordium geometry coincided with a delay in this fate restriction process in *katanin* mutants. Altogether, our study suggests that tissue geometry canalizes and modulates reproductive cell fate in the Arabidopsis ovule primordium.

## Introduction

A hallmark of sexual reproduction in multicellular organisms is the separation of the germline from the soma. In animals, primordial germ cells (PGCs) are set-aside during embryogenesis from a mass of pluripotent cells. The number of germ cells depends on the balance between proliferation (self-renewal) and differentiation, a process controlled by both intrinsic factors signals from and the surrounding somatic tissues. In flowering plants, the first cells representing the germline, the spore mother cells (SMCs), differentiate only late in development. SMCs arise multiple times, in each flower during the formation of the reproductive organs. In Arabidopsis, the female SMC differentiates in the nucellus of the ovule primordium, a digit-shaped organ that emerges from the placental tissue of the gynoecium. The SMC is recognizable as a single, large, and elongated subepidermal cell, in a central position, within the nucellus, displaying a prominent nucleus and nucleolus (Bajon, 1999; Bowman, 1993; Schmidt et al., 2015; Schneitz, 1995).

Although SMC singleness may appear to be robust, more than one SMC candidate *per* primordium is occasionally seen, yet at different frequencies depending on the *Arabidopsis* accession (9,6% in *Columbia* (*Col-0*), 27% in *Monterrosso* (*Mr-0*), (Rodriguez-Leal et al., 2015). Mutants in *Arabidopsis*, maize, and rice SMC singleness is compromised have unveiled the role of regulatory pathways involving intercellular signaling, small RNAs, as well as DNA and histone methylation (Garcia-Aguilar et al., 2010; Mendes et al., 2020; Nonomura et al., 2003; Olmedo-Monfil et al., 2010; Schmidt et al., 2011; Sheridan et al., 1996; Sheridan et al., 1999; Su et al., 2020; Su et al., 2017; Zhao et al., 2008). As the SMC forms, cell-cycle regulation contributes to the stabilization of its fate in a cell-autonomous manner through cyclin-dependent kinase (CDK) inhibitors and RETINOBLASTOMA-RELATED1 (RBR1) (Cao et al., 2018; Zhao et al., 2017). SMC singleness thus appears to result from a two-step control: first, by restricting differentiation to one cell and second, by preventing self-renewal before meiosis (reviewed in (Lora et al., 2019; Pinto et al., 2019).

However, the precise mechanisms underlying the plasticity in the number of SMC candidates and SMC specification are still poorly understood. In principle, SMC singleness may be controlled by successive molecular cues. However, even in that scenario, such cues must be positional, at least to some extent, and thus involve a spatial component. Over the last decade, many different molecular cues defining spatial patterns in the ovule primordium were identified (Pinto et al., 2019; Su et al., 2020), however their coordination is unknown. Since SMCs emerge at the primordium apex concomitant with its elongation, we hypothesize that geometric constraints during ovule morphogenesis influence SMC singleness and differentiation. Such an hypothesis could explain variation in the number of SMC candidates, ultimately culminating in a single SMC entering meiosis. Answering the questions of whether SMC formation follows a stereotypical or plastic developmental process and whether it is intrinsically linked to or independent of ovule primordium formation would unravel fundamental principles connecting cell fate establishment and organ growth.

Such an analysis requires a high-resolution description of ovule geometry during development. Our current knowledge of ovule primordium growth in *Arabidopsis* is based on two-dimensional (2D) micrographs from tissue sections or clearings. It is described in discrete developmental stages capturing classes of primordia by their global shape and SMC appearance until meiosis and by the presence of integument layers and ovule curvature later-on (Grossniklaus et al., 1998). In addition, a 3D analysis of average cell volumes during primordium growth was recently provided (Lora et al., 2017). Yet, we lack a view of the patterning processes regulating early ovule primordium formation and how the dynamics of cell proliferation contributes to the cellular organization during primordium growth. We thus described and quantified the growth of the ovule primordium at cellular resolution in 3D. We combined 3D imaging, quantitative analysis of cell and tissue characteristics, reporter gene analyses, 2D mechanical growth simulations. In addition, using the *katanin* mutant that affects anisotropic cell growth and cell division patterns (Luptovciak, Komis, et al., 2017; Ovecka et al., 2020), we show that altered ovule morphology lead to ectopic SMC candidates. We also uncovered that differentiation of SMC candidates initiate earlier than previously thought, and provide evidence for a gradual process of cell fate restriction, channeling the specification of a single SMC prior to meiosis.

## Results

### Building a reference image dataset capturing ovule primordium development at cellular resolution

To generate a reference image dataset describing ovule primordium development in 3D and with cellular resolution, we imaged primordia at consecutive stages in intact carpels by confocal microscopy. Carpels were cleared and stained for cell boundaries using a modified PS-PI staining (Truernit et al., 2008) and mounted using a procedure preserving their 3D integrity (Mendocilla-Sato, 2017) (Figure 1A). We selected high signal-to-noise ratio images and segmented them based on cell boundary signals using Imaris (Bitplane, Switzerland) as described previously (Mendocilla-Sato, 2017) (Figure 1B). We manually curated 92 ovules representing seven consecutive developmental stages (7-21 ovules *per* stage, Figure 1B, Table 1, Source Data 1), classified them according to an extended nomenclature (explained in Materials and Methods). The temporal resolution of our analysis led us to subdivide early stages (stage 0-I to stage 0-III) covering primordium emergence prior to the straight digit-shape of the organ set as stage 1-I, where the SMC becomes distinguishable by its apparent larger size in longitudinal views (Grossniklaus et al., 1998) (Figure 1B).

**Table 1.** Classification criteria of Arabidopsis ovule primordia. The table summarizes general characteristics of ovule primordia per stage: cell “layers” above the placenta scored as the number of L1 cells in a cell file drawn from the basis to the top, range thereof, total cell number, ovule shape including height, width, aspect ratio. See also Source Data 1.

|  | 0-I | 0-II | 0-III |
| --- | --- | --- | --- |
| Cells above placenta* | 2.5 (±3.5) (n=21) | 3.6 (±0.5) (n=17) | 5.9 (±1.0) (n=11) |
| range (min-max) | 2 - 3 | 3 - 4 | 5 - 8 |
| total # cells | 28 (±5.7) (n=21) | 38 (±8.0) (n=17) | 80 (±8.4) (n=11) |
| width (μm) (W) | 23 (±3.5) | 26.3 (±2.4) | 28.0 (±3.1) |
| height (μm) (H) | 5.3 (±1.2) | 11.9 (±2.1) | 22.5 (±3.6) |
| H:W ratio | 0.2 (±0.04) (n=13) | 0.45 (±0.06) (n=14) | 0.8 (±0.12) (n=8) |

|  | 1-I | 1-II |
| --- | --- | --- |
| cells above placenta* | 7.3 (±1.3) (n=15) | 9.7 (±1.0) (n=11) |
| range (min-max) | 6 - 10 | 8 - 11 |
| total # cells | 105 (±10.6) (n=15) | 128 (±18.0) (n=11) |
| width (μm) (W) | 27 (±2.0) | 27.8 (±3.5) |
| height (μm) (H) | 30 (±2.8) | 39 (±5.9) |
| H:W ratio | 1.1 (±0.10) (n=6) | 1.4 (±0.22) (n=8) |

**Table 1. Classification criteria of Arabidopsis ovule primordia.**
|  | 2-I | 2-II |
| --- | --- | --- |
| cells above placenta* | 10 (±1.1) (n=10) | 12.6 (±1.0) (n=7) |
| range (min-max) | 9 - 12 | 11 - 14 |
| total # cells | 151 (±18.9) (n=10) | 165 (±19.8) (n=7) |

\* scoring the length of L1 cell file above placenta

**Figure 1.**
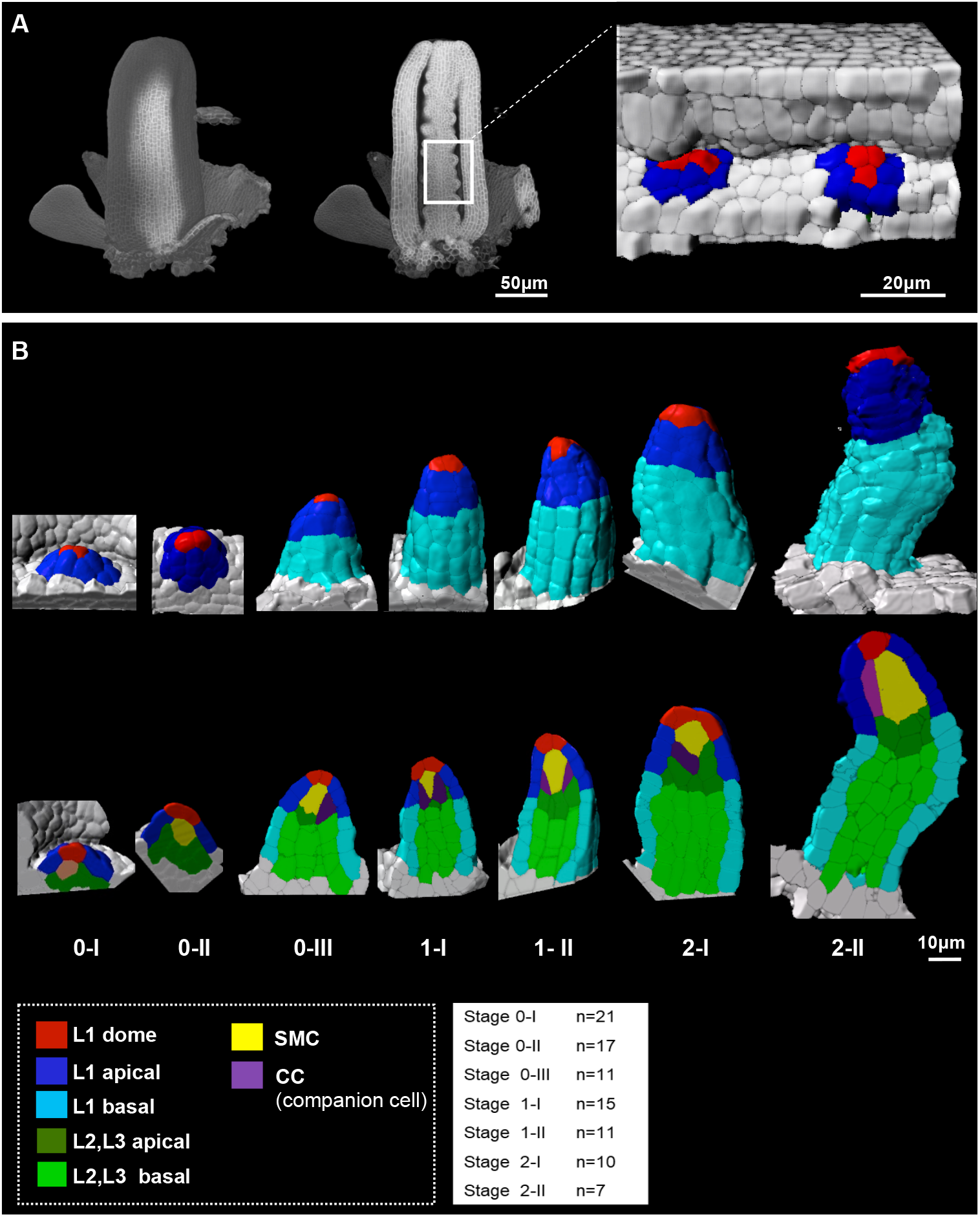
Reference set of 3D segmented images capturing Arabidopsis ovule primordium growth at cellular-resolution. (A) 3D reconstruction of a whole gynoecium stained with PS-PI (cell wall dye) and visualized by CSLM. The cross section allows showing the nascent ovule primordia attached to the placenta. (B) Ovule primordium developmental stages (0-I to 2-II) and organ viewpoints (domains) defined for the 3D quantitative analysis. All segmented data can be analyzed by an interactive interface named OvuleViz. See also Figure supplement 1A-C. n= number of ovules analyzed. Segmented images for all developmental stages are provided in Source Data 1. See also Figure supplement 1 D-E, Materials and Methods.

To evaluate the distinct contribution of domain-, layer- and cell-specific growth dynamics, we labeled cells depending on their belonging to different regions of the ovule primordium: apical vs basal domains, L1/L2/L3 layers. In addition we associated each cell with a cell type: “L1 apical”, “L1 basal”, “L2,L3 apical”, “L2,L3 basal”, “SMC”, “L1 dome” (for the upmost apical L1 cells in contact with the SMC), “CC” (for companion cells, elongated L2 cells adjacent to the SMC) (Figure 1B, Materials and Methods).

To generate a quantitative description of ovule primordia with respect to cell number, size and shape according to cell labels, layers, domains, ovule stage and genotype, we developed an interactive, R-based interface named OvuleViz. The interface imports cell descriptors exported from segmented image files and enables multiple plots from a user-based selection of (sub)datasets (Figure supplement 1A-C, Materials and Methods). This work generated a reference collection of annotated, 3D images capturing ovule primordium development at cellular resolution from emergence until the onset of meiosis. The collection of 92 segmented images comprising a total of 7763 annotated cells and five morphological cell descriptors (volume, area, sphericity, prolate and oblate ellipticity) provides a unique resource for morphodynamic analyses of ovule primordium growth.

To identify correlations between growth patterns and differentiation, we first performed a principle component analysis (PCA) based on the aforementioned cell descriptors, *per* cell type and stage, considered together or separately (Figure supplement 1D-E). In this global analysis, the SMC appears morphologically distinct at late stages (2-I and 2-II) but not at early stages. This prompted us to investigate in detail the contribution of different layers, domains, and cell types to ovule primordium growth and in relation to SMC differentiation.

### Ovule primordium morphogenesis involves domain-specific cell proliferation and anisotropic cell shape patterns

The ovule primordium emerges from the placenta as a small dome-shaped protrusion and grows into a digit-shaped primordium with nearly cylindrical symmetry (stage 1-I) before enlarging at the base (Figure 1B). Using our segmented images, we first quantified global changes in cell number, cell volume, and ovule primordium shape. Our analysis revealed two distinct phases of morphological events. Phase I (stages 0-I to 0-III) is characterized by a 4.5-fold increase in total cell number together with a moderate increase in mean cell volume (10%, P=0.03). By contrast, Phase II (stages 1-I to 2-II) is characterized by a moderate increase in cell number (50%) and the global mean cell volume is relatively constant (Figure 2A, Figure supplement 2A). To quantify the resulting changes in organ shape, we extrapolated a continuous surface mesh of the ovule outline and used it to compute its height and width at the base (Figure 2B-C, Figure supplement 2B, Supplemental File 1). Anisotropic organ growth during Phase I was confirmed by a steady increase in height while primordium width increased moderately (Figure 2C). This contrast in events between Phase I and II is illustrated by the fold-changes (FCs) in cell number and aspect ratio (Figure 2D) which range between 1.5 and 2.0 in Phase I whereas drop to 1.4 and 1.2 in Phase II, respectively. These observations confirmed that Phase I captures a distinct growth dynamic compared to Phase II.

**Figure 2.**
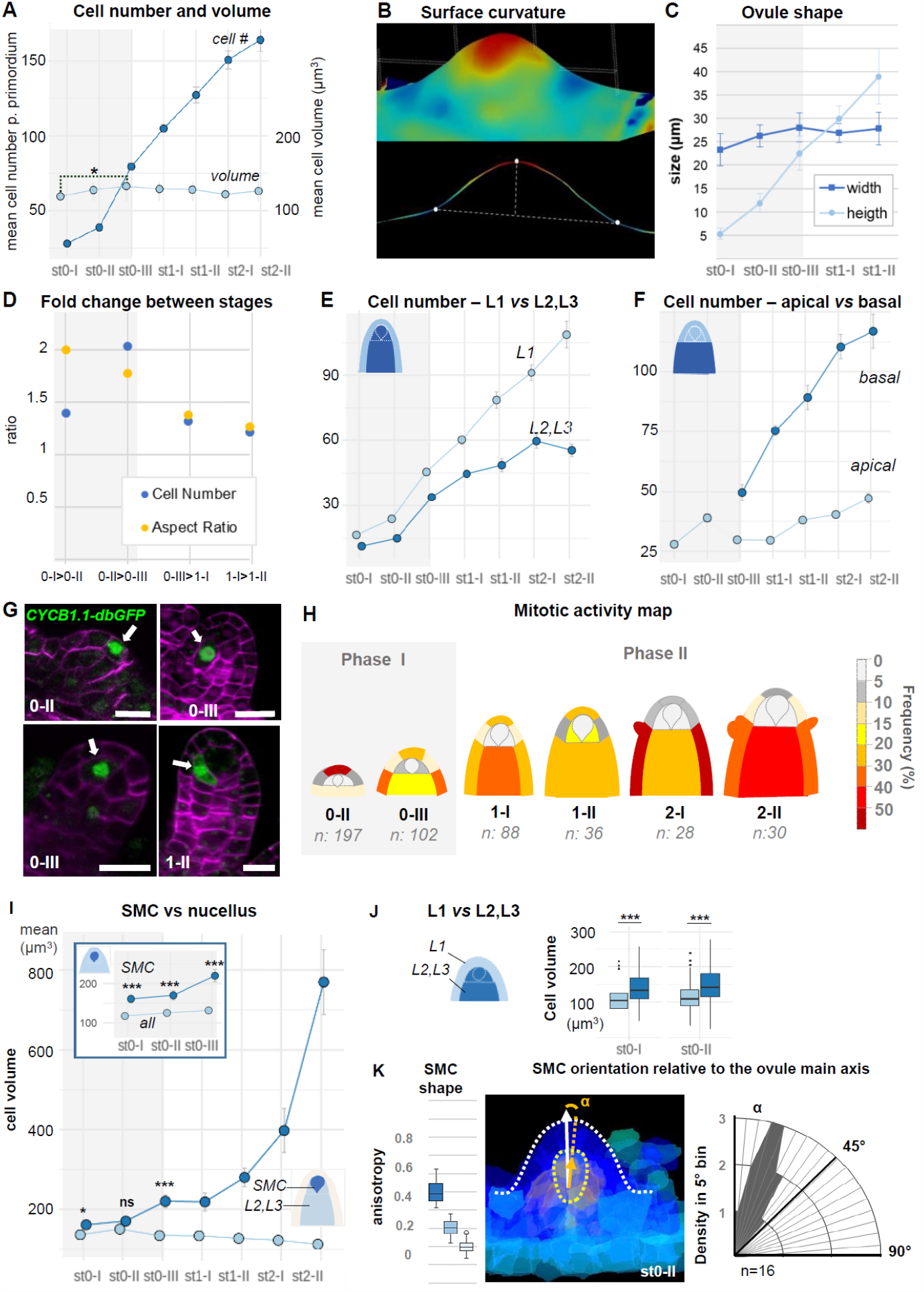
Ovule primordium morphogenesis involves domain-specific cell division and anisotropic cell growth. (A) Mean cell number *per* ovule increases mainly during stages 0-I to 0-III (Phase I), whereas cell volume *per* ovule remains constant on average across primordium development (stages 0-I to 2-II). (B) Representative image of a continuous surface of an ovule primordium mesh and its projected median plane. Dashed lines indicate the minimal and maximal curvature points used to measure organ height and width. Color scale: minimal curvature mm-1 (see also Figure supplement 2B). (C) Anisotropic organ growth during Phase I and until stage 1-II. Mean width and height were quantified *per* stage. (D) Phase I captures a distinct growth dynamic compared to Phase II. Fold-change of cell number and aspect ratio between stages are plotted. (E) Mean cell number is increased at the L1 *versus* L2,L3 layers across developmental stages. (F) Mean cell number is increased in the basal *versus* apical domain across developmental stages. (G) Representative images of ovule primordia expressing the M-phase reporter *promCYCB1*.*1::CYCB1*.*1db-GFP* (CYCB1.1db-GFP). White arrows indicate dividing cells. Magenta signal: Renaissance SR2200 cell-wall label. Scale bar: 10μm. (H) Domain-specific map of mitotic activity during ovule primordium development, scored using the CYCB1.1db-GFP reporter. The frequency of mitoses was calculated *per* ovule domain at each developmental stage and color-coded as indicated in the bar (right). n: total number of scored ovules. (I) Mean SMC candidate volume (dark blue) is significantly increased as compared to L2,L3 cells (pale blue) from stage 0-III onward, on even at earlier stages as compared to all other cells (inset). (J) Mean cell volume is increased in L2,L3 cells as compared to L1 cells, at the two early developmental stages (0-I, 0-II). (K) The SMC consistently displays anisotropic shape with main axis of elongation aligned with ovule growth axis. The SMC anisotropy index (boxplot, left; stage 0-II, n=16 ovules) was calculated from the Maximum (dark blue), Minimum (light blue) and Medium (medium blue) covariance matrix eigenvalues, computed from 3D segmented cells (see Figure supplement 2G). Image (middle): illustration of the SMC main anisotropy axis (orange arrow) related by an angle ‘alpha’ to the main axis of the ovule primordium (white arrow), stage 0-II. Radar plot (right): ‘alpha’ angle measured on z projections for n=16 ovule primordia at stage 0-II. See also Materials and Methods. Error bars: Standard errors to the mean. Differences between cell types or primordium domains were assessed using a two-tailed Man Whitney U test in (A) and (I); a two-tailed Wilcoxon signed rank test in (J). *p≤0.05, **p≤0.01, ***p≤0.001. See also Figure supplement 2, Source Data 2, Materials and Methods.

Next, to capture possible specific patterns of growth, we analyzed cell number, cell size, and cell shape using different viewpoints: one comparing the L1 and L2-L3 layers and one contrasting the apical *vs*. basal domains. Counting cell number *per* viewpoint clearly showed a dominant contribution of the epidermis (L1) relative to the subepidermal layers and of the basal relative to the apical domain (Figure 2E, 2F). To verify these findings with a cellular marker, we analyzed the M-phase-specific *promCYCB1*.*1::CYCB1*.*1-db-GFP* reporter (abbreviated CYCB1.1db-GFP) (Ubeda-Tomas et al., 2009). We scored the number of GFP-expressing cells among 481 ovules and plotted relative mitotic frequencies *per* cell layer and domain for each ovule stage to generate a cell-based mitotic activity map (Figure 2G-H, Source Data 2, Materials and Methods). In this approach, subepidermal (L2) cells beneath the dome were distinguished from underlying L3 cells to gain resolution on the L2 apical domain where the SMC differentiate. Consistent with our previous observation, in Phase I, a high proliferation activity was scored in L1 cells at the primordium apex (scoring 64% of mitotic events). By contrast, the L2 apical domain remains relatively quiescent (contributing only 3% of the mitotic events). During Phase II, the majority (60%) of mitotic events is found in the basal domain, consistent with the progressive population of the basal domain. It is noteworthy that during this phase, few mitotic events are detected in L2 apical cells, with the exception of SMC neighbor cells that show frequent divisions at stage 1-II. Thus, genetic reporter analysis confirmed a biphasic, temporal pattern of cell division with changing regional contributions, suggesting the L1 dome and the basal domain as consecutive sites of proliferation, contributing to morphological changes in Phase I and II, respectively.

Average cell size analysis, by contrast, did not reveal significant changes during primordium development with the notable exception of the SMC (Figure 2A, Figure supplement 2C). The distinct size of the SMC candidate is already detected at stage 0-III when compared to other L2,L3 cells (Figure 2I) or even earlier (stage 0-I), when compared to all other cells (Figure 2I inset). Size differentiation of the SMC is not uniform among ovules, demonstrating plasticity in the process (Figure supplement 2D). In addition, cells from the L2 and L3 layers are larger than L1 cells already at stage 0-I (Figure 2J, Figure supplement 2E) possibly due to a longer growth phase, consistent with the low division frequency observed previously.

We then investigated cell shape changes during primordium elongation, using ellipticity and sphericity indices computed following segmentation. The analysis did not reveal significant differences between domains or layers (Figure supplement 2F). This could indicate either a highly variable cell shape or local, cell-specific differences. We thus more specifically analyzed the subepidermal domain where the SMC differentiates. Companion cells showed an increasing ellipticity (and decreasing sphericity), starting at stage 1-I and culminating at stage 2-I (Figure supplement 2F). By contrast, the SMC only showed a moderate decrease in sphericity at late stages and no distinctive ellipticity at early stages (Figure supplement 2F) when compared to other cells. To get more information on SMC shape, we compared the maximum, medium and minimum anisotropy index. For this, we developed an extension for MorphoMechanX (Barbier de Reuille et al., 2015), (www.morphomechanx.org) to (i) perform a semi-automatic labeling of cell layers and cell types from a cellularized mesh obtained from segmentation data and (ii) compute in each 3D cell the principal axes of shape anisotropy and the corresponding indexes (Figure supplement 2G, Supplemental File 1, movie in Supplemental File 2). The averaged maximum anisotropy shape index of the SMCs was consistently above the medium (and hence above the minimum) anisotropy index (Figure 2K, Figure supplement 2H). We measured the degree of alignment of the SMC major axis with the main growth axis of the ovule during early stages (stage 0-II) and found a mean angle of 22° (+/- 11°, n=16) (Figure 2K). This confirmed that the SMC has a consistent anisotropic shape from early stage onwards, with a distinguishable major axis aligned with primordium axis.

Taken together, these results suggest that anisotropic primordium growth is linked to a biphasic, domain-specific cell proliferation, alternating between the L1 dome at phase I and the basal domain at phase II, combined with localized, anisotropic expansion in the L2 apical domain. In this process, SMC characteristic distinct size, anisotropic shape and orientation aligned with the growth axis of the primordium emerge already in Phase I, while entering a pronounced growth and elongation at Phase II concomitantly to organ shape elongation. While primordium elongation is not explained by anisotropic cell growth alone but also by cell proliferation (Figure supplement 2F), the observation that cells are elliptic suggests a potential role for anisotropic cell growth. We explore this property in the next section through an *in-silico* approach.

### 2D mechanical simulations of ovule primordium development relate its coordinated growth to SMC shape emergence

Organ shape is determined by the rate and direction of cell growth, in turn affected by signaling and the mechanical state of the tissue and its geometry, with possible feedback interactions between them (Bassel et al., 2014; Echevin et al., 2019). Mechanical constraints arise from the growth process in form of tensile and compressive forces that, in turn, influence cell and tissue growth depending on their elastic/visco-plastic properties (Echevin et al., 2019). To determine the role of ovule primordium growth on SMC differentiation, which involves early differential cell growth, we sought to understand the contributions of local growth rate and anisotropy, and their relation to signaling and mechanical constraints (Coen et al., 2004; Kennaway et al., 2011).

We first aim at determining morphogenetic principles of primordium growth at the tissue level by developing a 2D Finite Element Method (FEM)-based mechanical model, which considered two uncoupled but complementary modes of growth (Bassel et al., 2014): (i) signal-based growth in which an abstract growth factor captures the cumulative effects of biochemical signals, without considering its explicit mode of action (i.e., cell wall loosening, increase in turgor pressure) (Boudon et al., 2015; Coen et al., 2004); and (ii) passive strain-based growth in which relaxation accommodates excess of strain (and therefore also excess of stress) that accumulated in the tissue, by relaxing its reference configuration according to the amount of strain(Boudon et al., 2015; Bozorg et al., 2016). The mechanical equilibrium is computed to ensure compatibility within the tissue that is locally growing at different rates and orientations. This introduces residual internal compressions and tensions to which the action of turgor pressure has to be added up (Boudon et al., 2015; Rodriguez et al., 1994). A polarization field is used to set the direction of anisotropic growth for both growth mechanisms (Coen et al., 2004). The simulation consists of iterations where the mechanical equilibrium is computed before each growth step is specified by signal- and strain-based growth (Supplemental File 1). This strategy extends previous tissue growth models (Bassel et al., 2014; Boudon et al., 2015; Bozorg et al., 2016; Kuchen et al., 2012; Mosca, 2018).

We designed a starting template consisting of an L1 layer distinct from the underlying L2, L3 tissue based on different growth and material properties (Table 2, Figure supplement 3, Supplemental File 1). We first set the model components to produce a realistic elongated, digit-shaped primordium with a narrow dome and an L1 layer of stable thickness during development, fitting experimental observation (Figure supplement 2C) (Reference Model, FEM-Model 1) (Figure 3A, Figure supplement 3A). This model combines the following hypotheses: an initial, narrow domain of anisotropic, signal-based growth with a high concentration in inner layers, a broad domain competent for passive strain-based relaxation and material anisotropy.

**Table 2.** Hypotheses used to generate the mass-spring (MS) and continuous Finite Element Method (FEM) based simulations. Several growth and mechanical hypotheses were listed at start of modelling. To evaluate their effect on primordium growth each hypothesis was excluded (-) in at least one scenario. The FEM and mass-spring (MS) models presented in Figure 3 and S3 are numbered according to the scenarios (1-8) in the table. Since it is not possible for MS to simulate material anisotropy, Model 1 was only tested with FEM. The hypothesis of Growth anisotropy is always active for the L1 layer in the models reported in the table. Empty dots for “Material anisotropy” were considered only for the FEM models. See also Figure supplement 3 and Supplemental File 1 for modelling principles, results and detailed computational methods.

| Modeling hypotheses |  | Models |  |  |  |  |  |  |  |
| --- | --- | --- | --- | --- | --- | --- | --- | --- | --- |
|  |  | 1 | 2 | 3 | 4 | 5 | 6 | 7 | 8 |
| Growth anisotropy |  | ● | ● | - | ● | ● | ● | ● | ● |
| Material anisotropy |  | ● | - | ○ | ○ | ○ | ○ | ○ | ○ |
| Passive Strain-based relaxation |  | ● | ● | ● | ● | ● | ● | ● | - |
| Signal-based growth |  |  |  |  |  |  |  |  |  |
| Distribution | L1 only | - | - | - | - | - | ● | - | - |
|  | Inner tissue only (pi-shape) | - | - | - | - | - | - | ● | - |
|  | L1 + inner tissue (pit-shape) | ● | ● | ● | - | ● | - | - | ● |
|  | L1 + inner tissue (broad distribution) | - | - | - | ● | - | - | - | - |
| Fixed high concentration | L1 only | - | - | - | - | ● | ● | - | - |
|  | Inner tissue only | - | - | - | - | - | - | ● | - |
|  | L1 + inner tissue | ● | ● | ● | ● | - | - | - | ● |

**Figure 3.**
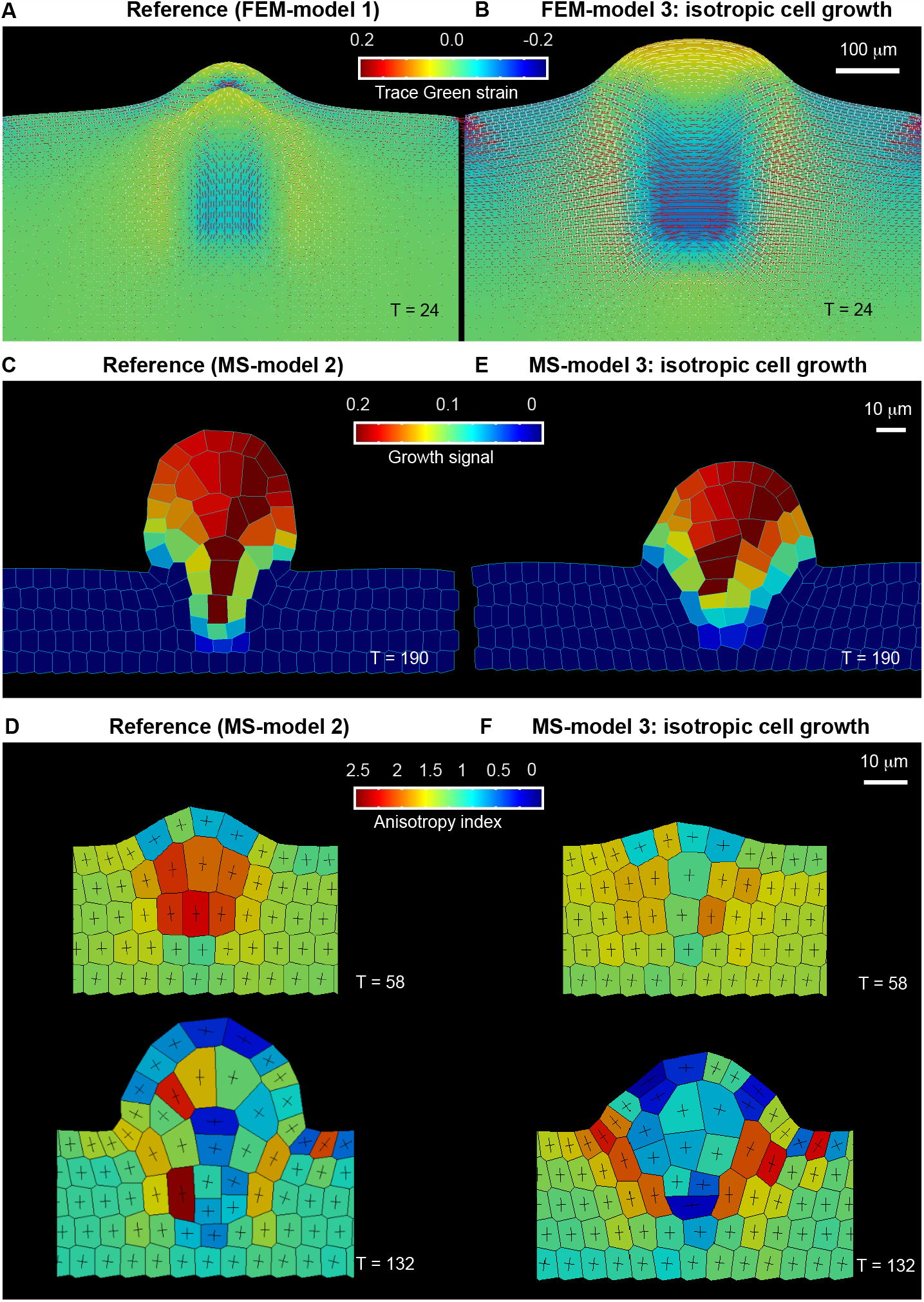
Mechanical and cell-based 2D simulation models of ovule primordium development predict that ovule shape depends on cell growth anisotropy. (A-B) Finite Element Model (FEM), tissue-based simulations of ovule primordium growth. (A) Growth stability is reached at T=24 in the reference model FEM model 1. (B) Simulations omitting the anisotropic growth parameter (FEM-model 3) show abnormal ovule dome shape at the same simulation time. The level (Trace green strain) and principal directions (fine lines, white: expansive, red: compressive) of accumulated strain is shown. (C-F) Mass Spring (MS) cell-based simulations of ovule primordium growth showing the growth signal distribution and anisotropy index at the indicated simulation times (T). (C) Reference model (MS-model 2) showing a realistic primordium shape with straight flanks, sharp curvature at the apex and narrow base at T=190. (D) The reference model shows the emergence of a large, anisotropic cell with trapezoidal shape in L2 at T=58, confirmed at T=132. (E) Simulation with isotropic cell growth in inner layers (MS-model 3) produces a primordium with enlarged apex and basis and flatter dome at the same simulation time as in the reference model. (F) L2 cells in MS-model 3 show reduced anisotropy as compared to the Reference model. See also Figure supplement 3, Table 2, Supplemental File1 for modelling hypotheses and methods.

Then, using the versatility of the modeling framework to vary initial conditions, we tested the influence of the spatial distribution of the specified growth signal on primordium growth. As first variation, we let the growth signal diffuse broadly in the domain while maintaining the selected initial cells with prescribed high intensity growth signal (FEM-Model 4). The emerging primordium is appreciably broader than the Reference Model (Figure supplement 3B). We then explored what is the contribution of growth signal in the L1 compared to in inner tissue to the primordium growth. In model FEM-Model 5, prescribed high growth signal is present only in the L1 and it is free to diffuse in the inner layers. This model produced a sharp primordium, narrower and taller than the Reference Model, but does not preserve L1 thickness (Figure supplement 3B). The L2 apical domain is narrower as compared to the Reference Model. When the growth signal is absent in the inner tissue (FEM-Model 6), signal growth in the L1 alone is not sufficient to enable primordium growth (Figure supplement 3B).

To answer the complementary question whether growth signal is required in the L1 to grow a primordium when this is present in the inner layers, this hypothesis (Table 2) was removed in FEM-Model 7 (Figure supplement 3B). This scenario indeed enables to grow a digit shape structure (as long as strain-based growth is permitted), yet the dome appears shallower than the Reference Model. To conclude, both models where high growth signal is selectively present only in L1 or in inner tissue layers can produce a primordium. Yet, absence of growth signal in the inner layers result in drastic shape alterations. This favors a scenario where inner tissue-driven growth has a fundamental role.

Next, through modulation of passive strain-based growth, we determined that a broad tissue domain uniformly competent for strain accommodation is necessary to resolve the high accumulation of stress, which limits primordium elongation (FEM-Model 8, Figure supplement 3C). We also explored the contribution of material anisotropy (FEM-Model 2 and FEM-Model 2a, Figure supplement 3C): even in the case of isotropic material, it is possible to grow a digit-shape protrusion, yet with a wider dome, and an increased L1 thickness, in contrast with experimental observations. To restore the L1 thickness, it is sufficient to prescribe material anisotropy in the L1 exclusively.

Finally, we asked whether growth anisotropy must be specified in the model or if organ geometry and mechanical constraints are sufficient to specify primordium shape. Clearly, removing growth anisotropy component abolished the digit shape of the primordium and produced a hemi-spherical protrusion (FEM-Model 3, Figure 3B).

In summary, we identified parsimonious growth principles shaping the ovule primordium and suggesting different contributions of the epidermis and inner layers: an active tissue growth, mostly inner-driven and requiring a narrow, pit-shape domain of growth-signal, complemented by passive tissue growth with a necessary response of the L1 to accommodate accumulated strain. Furthermore, material anisotropy in the L1 is predicted to play a role in constraining L1 thickness as observed experimentally.

Next, we aimed to analyze how cell division and cell shape, particularly that of the SMC, are connected to tissue growth patterns at cellular scale. For this, we developed a complementary model with cellular resolution. This vertex-based model features cells as 2D polygons whose walls are mass spring (MS) segments (Amar, 2011; Smith, 2009). This model integrates mechanics (Hooke’s law for MSs) with signal- and strain-based growth as before (Supplemental File 1), and cells are able to divide after they reach a threshold area, the newly inserted walls following the rule “shortest wall through centroid” (Supplemental File 1). We simulated growth starting with a virtual grid of cells representing a flat placenta and focused on early stages of ovule primordium formation until stage 1-I. We tested the same combination of growth hypotheses as for the continuous model (Table 2), except for material anisotropy that cannot be easily embedded in a vertex-based model. As we observed that L1 cells do not divide periclinally, nor change in volume on average (see Figure supplement 2C), we deduced that their growth occurs mainly along the surface of the primordium. For this reason, we enforced this feature in our model, preventing growth of anticlinal walls in the L1 (both for signal-based growth and passive strain-based growth, Supplemental file 1). This constraint limits the ways an organ can grow in a digit-shaped fashion, as the L1 does not progressively populate the inner tissues by division. Furthermore, our models allowed for larger cells in subepidermal layers compared to the epidermis, in agreement with our observation (Figure supplement 2E).

We observed that MS-Model 2 best captures the characteristic shape and aspect ratio of ovules (Figure 3C-D, Figure supplement 3D). This Reference Model confirmed the growth hypotheses from FEM-based simulations regarding the distribution and orientation of signal-based growth (Figure supplement 3B) and passive strain-based growth (Figure supplement 3C). Altering growth hypotheses similarly to that in the FEM models produce similar results (Figure Supplement 3B,C), yet with more dramatic shape changes in the case of lack of strain-based relaxation (MS-Model 8 and MS-Model 8a) and growth signal concentration in inner tissue (MS-Model 5, Supplemental File 3). In the Reference Model (MS-Model 2), an L2 apical cell with a trapezoidal shape, elongated along the main direction of ovule growth, emerged consistently during simulation (note that these cells still divide in the models as no special rule has been assigned to them) (Figure 3D). These are similar to SMC candidates at stage 0-II in real primordia, in a 2D longitudinal, median section through the ovule (Figure 2K). The elongated-trapezoidal shape of such cell is not a prescribed feature of the model, but rather emerges from the combination of assigned anisotropic cell growth and geometrical constraints imposed by the surrounding growing tissues.

Next, we wanted to explore the role of cell growth anisotropy in ovule primordium shape and SMC emergence. The MS-Model 3 (Figure 3E-F) corresponds to a virtual mutant where cell growth was set to isotropic in inner layers (the L1 was maintained with anisotropic growth to preserve its thickness). This led to an ovule primordium with a wider and flatter dome, comparable to the FEM-Model 3. Despite the absence of specified growth direction in inner tissue, due to geometrical constraints, the primordium still grows mostly vertically. We wanted to assess whether such geometrical constraints would enable the formation of an elongated, trapezoidal SMC candidate also in the case of prescribed isotropic growth. As Figure 3E-F shows, the SMC candidate does not display a stereotypical shape and is not even elongated. Yet, mild cell anisotropy can be reached if the SMC candidate is allowed to grow twice more than in the reference model (Figure Supplement 3E, MS-Model 3a).

Altogether, the different simulations suggest that SMC anisotropy and characteristic shape can also be an emerging property of ovule primordium growth connected to geometrical constraints, even if, in the absence of specified anisotropic growth. The complementary model altering cell growth anisotropy specifically in the L1 further suggested that directional cell growth in this layer is necessary to accommodate inner-driven growth and permit primordium elongation (MS-Model 3b, Figure supplement 3F).

In all of the above simulations, the candidate SMC eventually divided as the models missed a causative rule to differentially regulate cell division. When we prevented the SMC candidate to divide, its enlargement overrode primordium shape control in our simulations creating an enlarged dome (Figure supplement 3G). This suggests that a mechanism may limit SMC growth in real primordia.

Altogether, 2D mechanical-based simulations in both continuous tissue, or cell-based approaches confirmed a role for localized cell growth compatible with experimental observations. The simulations also pointed to the importance of a differential role for the epidermis (accommodation) and the inner tissue (major growth component) as well as anisotropic cell growth as necessary for ovule primordium shape. Further, the simulations suggested that SMC formation can be an emerging property of primordium geometry.

### *katanin* mutants show a distinct ovule primordium geometry

To experimentally test the prediction of the isotropic growth models we analyzed ovule primordium growth and SMC fate establishment in *katanin* mutants with well described and understood geometric defects: in absence of the microtubule-severing protein KATANIN, the self-organization of cortical microtubules in parallel arrays is hindered, thereby decreasing the cellulose-dependent mechanical anisotropy of the wall and resulting in more isotropic growth (Bichet et al., 2001; Burk et al., 2002). Here, we specifically analyzed the shape and cellular organization of ovule primordia in *katanin* mutants using the *botero* (*bot1-7*) (*Ws* background), *lue1*, and *mad5* alleles (*Col* background) (Bichet et al., 2001; Bouquin et al., 2003; Brodersen et al., 2008), collectively referred to *katanin -* noted *kat -* mutants hereafter. We generated and analyzed a new dataset of 59 annotated 3D digital *kat* and corresponding wild-type ovule primordia at stages 0-III, 1-I, and 1-II (Source Data 4A). *kat* mutant primordia clearly showed an increased size and a more isotropic shape (Figure 4A), being 1.5 times bigger in volume than wild-type primordia (*P*=0.007, stage 0-III, Figure 4B) with a smaller aspect ratio (*P*<0.01 stages 1-I, 1-II, Figure 4C). Because the width-to-height ratio does not inform on the shape at the flanks, we derived an equation to estimate the “plumpiness” of the primordia: primordia with rounder flanks will have a higher bounding box occupancy, that is, the volume fraction of a fitting, 3D parallelepiped (bounding box) effectively occupied by the primordium (Figure 4D, left), than straight-digit shape ovules of the same aspect ratio. Mutant primordia clearly deviate from wild-type primordia in their relationship between aspect ratio and bounding box occupancy relationship at stage 1-I (Figure 4D, Figure supplement 4A), when the primordium normally starts elongating along the major growth axis. These measurements confirmed a marked attenuation of anisotropic growth in *kat* ovule primordia as was observed in roots, shoot organs, or seeds (Bichet et al., 2001; Hervieux et al., 2016; Luptovciak, Samakovli, et al., 2017; Ren et al., 2017; Uyttewaal et al., 2012; Wightman et al., 2013; Zhang et al., 2013). Yet, at a global level, the mean cell number, cell volume, and sphericity are not significantly different between *kat* and wild-type primordia (Figure supplement 4B-C). Thus, these global approaches did not resolve the origin of primordia mis-shaping.

**Figure 4.**
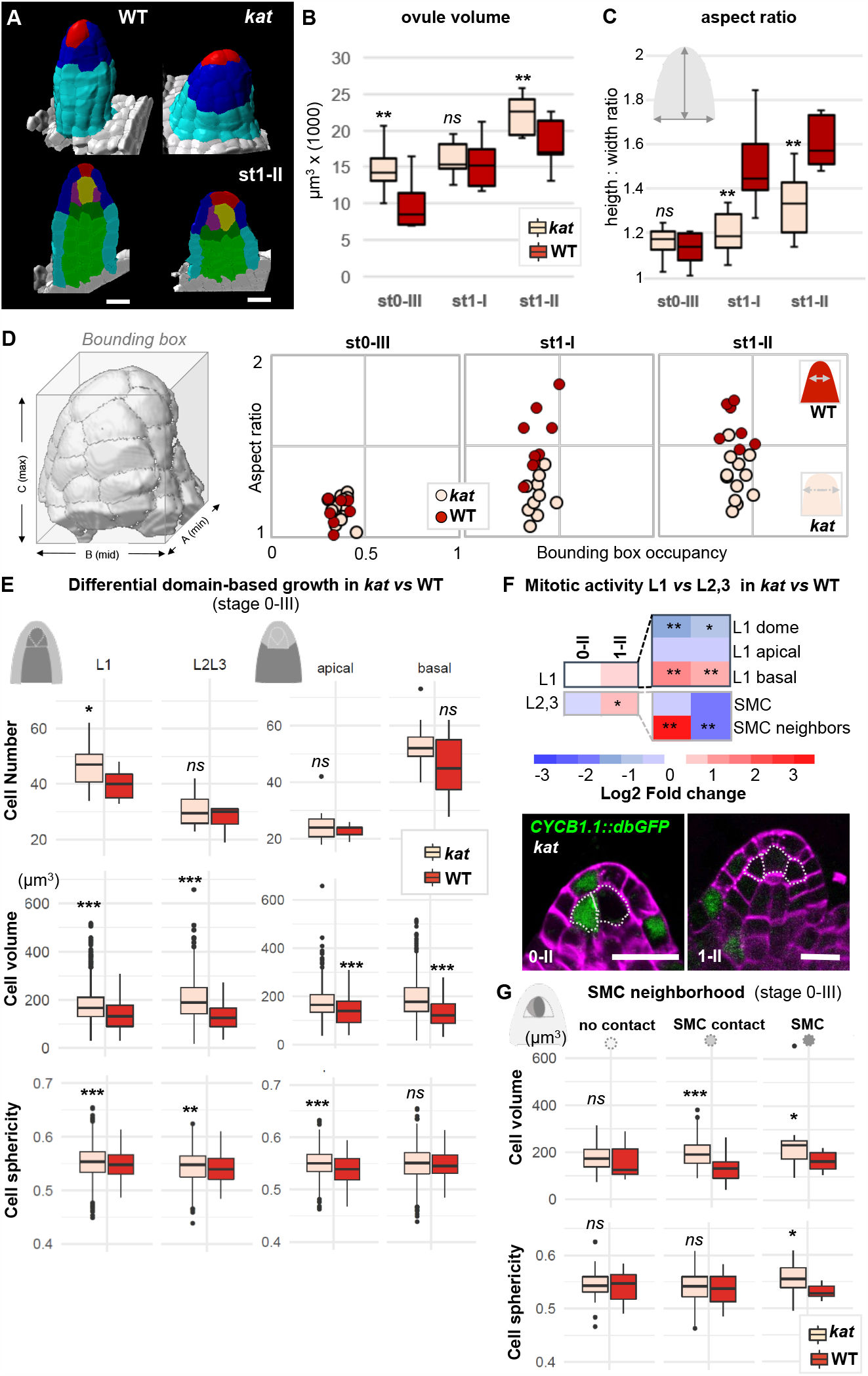
*katanin* mutants show a distinct ovule primordium geometry. Comparison between wild-type (WT, Ws-4 accession) and *katanin* (*kat, bot1-7* allele) ovule primordia. (A) 3D segmented images at stage 1-II. External organ view (top) and longitudinal sections (bottom) are shown. Scale bar 10μm. See Source Data 2A for full datasets. (B-D) Morphological difference between WT and *kat* primordia measured by their volume (B), the aspect ratio (C), and aspect ratio to bounding box occupancy relationship (D). D, Left: scheme representing the bounding box capturing the primordium’s 3D surface. See also Figure supplement 4A. (E) Quantification of cell number, cell volume, and sphericity in comparisons of L1 *versus* L2,L3 and apical *versus* basal domains at stage 0-III. See also Figure supplement 4C. (F) Mitotic activity domains are altered in *kat* primordia. Top: Heatmap of Log2 fold change of mitotic activity in the *kat* mutant (*lue1* allele) *versus* WT (Col-0) *per* domain, at two developmental stages. The frequency of mitoses was measured as in Figure 2. Full maps of mitotic activity in different mutant alleles are shown in Figure supplement 4E. Bottom: representative images of *kat* primordia (*lue1* allele) expressing CYCB1.1db-GFP. Dashed lines mark L2 apical cells. Magenta signal: Renaissance SR2200 cell wall label. Scale bar 10μm. (G) Mean cell volume and sphericity are increased in *katanin* L2 apical cells in contact with the SMC. SMC, cells in contact with the SMC, and cells not in direct contact with the SMC, are compared at stage 0-III. Color code in all plots: Dark red: WT; Salmon: *kat* mutant. Error bars: standard error of the mean. Differences between WT and *kat* mutants in (B), (C), (E), and (G) were assessed using a Mann Whitney U test; a two-tailed Fischer’s exact test was used in (F). P values: *p≤0.05, **p≤0.01, ***p≤0.001. See also Figure supplement 4 and Source Data 4.

To refine the analysis, we contrasted different layers as done for wild-type primordia and found local alterations that provide an explanation for the formation of broader and more isotropic primordia in *kat* mutants. Indeed, at stage 0-III *kat* ovules display a broader epidermis domain composed of more and larger cells than in wild-type (*P*=0.03 and *P*<0.001, respectively) (Figure 4E). Mitosis frequency analysis indicated a shift in cell division from the apex towards the basis (2.4 times less mitoses in the L1 dome domain and 2.7 times more in the L1 basal domain, compared to wild-type) (Figure 4F, Figure supplement 4E), consistent with the increased cell number observed in L1. Yet increased divisions is not a general characteristic of *kat* primordia since overall, the apical and basal domains are not massively overpopulated. By contrast, *kat* cells are generally larger and slightly more spherical in all domains (Figure 4E, Figure supplement 4C-D). When looking specifically at the L2 apical domain, where the SMC differentiates, we noticed an increased relative frequency of CYCB1.1db-GFP expression specifically in *kat* SMC neighbors at stage 0-II, whereas it decreases at stage 1-II (Figure 4F, Figure supplement 4E). This is in stark contrast with SMC neighbors in wild-type primordia displaying first a relative mitotic quiescence at stage 0-II, then enhanced mitotic activity at stage 1-II. Yet, in the SMC, no mitotic activity increased was observed in *kat* as compared to wild-type primordia (Figure 4F, Figure supplement 4E).

In conclusion, the absence of *KATANIN*-mediated cell growth anisotropy is associated with spatio-temporal shifts in cell divisions leading to an altered primordium geometry including a flatter dome and enlarged basis. Interestingly, this coincided with the occurrence of additional large cells in the L2 apical domain of *kat* primordia (SMC direct neighbor cells, Figure 4G, Materials and Methods) similar in size to the SMC. This raises the question of the identity of these ectopic, enlarged cells.

### Altered ovule primordium geometry in *katanin* mutants induces ectopic SMC fate

Following the observation of misshaped *kat* primordia and coinciding with larger L2 apical cells, we asked whether this had an impact on cell identity and SMC establishment. Like in wild-type, a clear SMC is identifiable in *kat* primordia yet it appears slightly bigger and more spherical with some variability over stages (Figure 5A, Figure supplement 5A). Interestingly, and consistent with the increased mitotic frequency in SMC neighbors observed at earlier stages, at stage 1-II, we scored additional SMC neighbors in *kat* primordia, that were on average larger (14%, *P*=0.006) and slightly more isotropic in shape (∼11% less ellipsoid *P*=0.03, ∼2% more spherical, *P*<0.01) than in the WT (Figure 5B). At stage 1-II, 34% *kat* primordia (n=109) showed more than one enlarged, subepidermal cell; decreasing to 14.4% at stage 2-II (n=104), in stark contrast with wild-type primordia showing a majority (86 to 96%) of primordia with an unambiguous, single SMC (Figure 5C). The *kat* phenotype is thus reminiscent of mutants affecting SMC singleness (Pinto et al., 2019) and we hypothesized that these enlarged cells could be ectopic SMC candidates.

**Figure 5.**
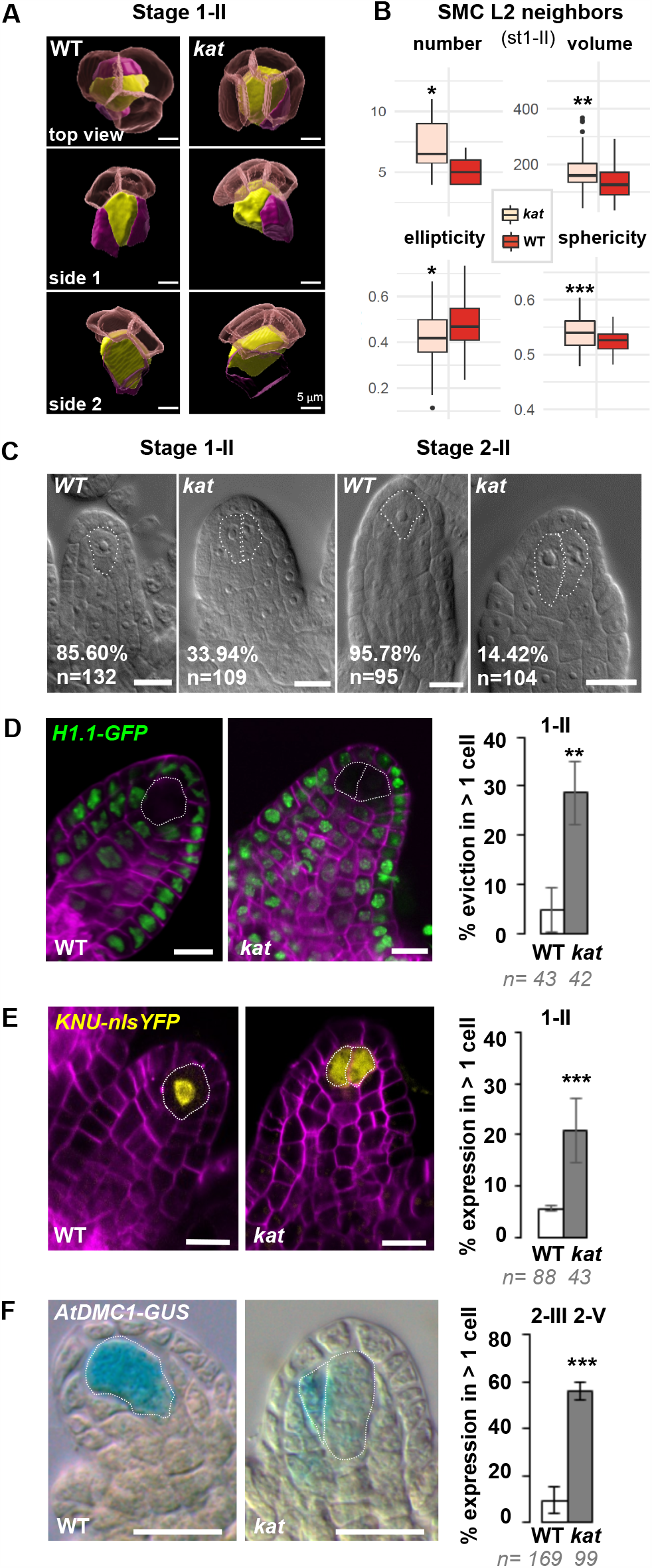
Altered ovule primordium geometry in *katanin* mutants is associated with multiple SMCs. (A) SMCs lose their typical pear shape in *katanin* (*kat*) mutants. 3D images of the apical-most cells according to top and side views as indicated, showing the SMC (yellow), SMC neighbors (purple), and the L1 dome (transparent red). (B) Differential properties of L2,L3 apical domain cells (SMC and SMC neighbors) in terms of cell number, mean cell volume, ellipticity, and sphericity at stage 1-II. See also Figure supplement 5A and Source Data 4B. (C) Representative images of cleared wild-type (WT) and *kat* ovule primordia. The % indicate the frequency of ovules showing one SMC for WT primordia, or multiple SMC candidates (dashed lines) for *kat* primordia. (D-E) Representative images and quantification of SMC fate markers in WT and *kat* primordia: eviction of the *H1*.*1::H1*.*1-GFP* marker (green, D) and ectopic expression of *KNU::nls-YFP* (yellow, E) in more than one cell *per* primordia, are increased in *kat* primordia. See also Figure supplement 5B. (F) The meiotic marker *AtDMC1::GUS* is ectopically expressed in *kat* ovules. Mutant alleles: *bot1-7* (A-B), *mad5* (C-E), *lue1* (F). Additional *kat* alleles, stages and detailed quantifications are presented Figure supplement 5 and Source Data 5. Magenta signal in (B) and (C): Renaissance SR2200 cell wall label. Scale bars for (A): 5 μm; for (C), (D), (E), (F): 10μm. n: number of ovules scored. Error bar: standard error of the mean. Differences between WT and *kat* genotype*s* were assessed using a Mann Whitney U test in (B), and a two-tailed Fischer’s exact test in (C), (D), (E), and (F). P values: *p≤0.05, **p≤0.01, ***p≤0.001. See also Figure supplement 5 and Source Data 5.

To verify this hypothesis, we introgressed several markers of SMC identity in *kat* mutant alleles. The first marker is a GFP-tagged linker histone variant (H1.1-GFP) that marks the somatic-to-reproductive fate transition by its eviction in the SMC at stage 1-I (She et al., 2013). H1.1-GFP eviction occurred in more than one cell in 28.6% of *kat* primordia (n=43, Figure 5D). The second marker reports expression of the *KNUCKLES* transcription factor (KNU-YFP) in the SMC (Tucker et al., 2012). Detectable as early as stage 1-I in the wild type, it was ectopically expressed in 21% of *kat* primordia at stage 1-II (n=43) (Figure 5E, Figure supplement 5B). Third, to test whether the ectopic SMC candidates have a meiotic competence, we analyzed the AtDMC1-GUS reporter (Agashe et al., 2002; Klimyuk et al., 1997), and indeed scored ectopic expression in 57% of *kat* primordia (n=56) (Figure 5F). We also analyzed the *pWOX2-CenH3-GFP* marker labeling centromeres starting at the functional megaspore stage (De Storme et al., 2016) and found 20.7% *kat* primordia (n=29) with two labeled cells, as compared to 3.8% (n=26) in the wild type (Figure supplement 5C).

Taken together, these results show that ectopic, abnormally enlarged SMC neighbors in *kat* primordia show at least some characteristics of SMC identity, and ectopic spores are also observed, indicating that reproductive fate is altered in *kat* ovule primordia.

### SMC singleness is progressively resolved during primordium growth

Based on our analysis of *kat* primordia, it appeared that the frequency of ectopic SMC candidates was high at early stages but decreased over time (Figure 5C). This is reminiscent of the phenotypic plasticity in SMC differentiation observed, to a lesser degree, in different *Arabidopsis* accessions (Rodriguez-Leal et al., 2015). To characterize this plasticity during development we analyzed 1276 wild-type primordia from stage 0-II to 2-II and scored the number of primordia with one or two enlarged, centrally positioned, subepidermal cells (class A and B, respectively, Figure 6A-C). In the wild type, the majority of ovules showed a single candidate SMC (class A) at stage 0-II but 27% of primordia (n=289) had two SMC candidates (class B), this frequency decreasing to 3% at stage 2-II (n=103). This finding is consistent with ∼ 5% primordia in wild-type at stage 1-II showing H1.1-GFP eviction (n=43) and KNU-YFP expression (n=88) in more than one cell, respectively (Figure 5D-E). Thus, instead of being immediate, SMC singleness can arise from a progressive restriction of fate among several SMC candidates during development. In wild-type primordia, SMC singleness is largely resolved at stage 1-I (Figure 5E).

**Figure 6.**
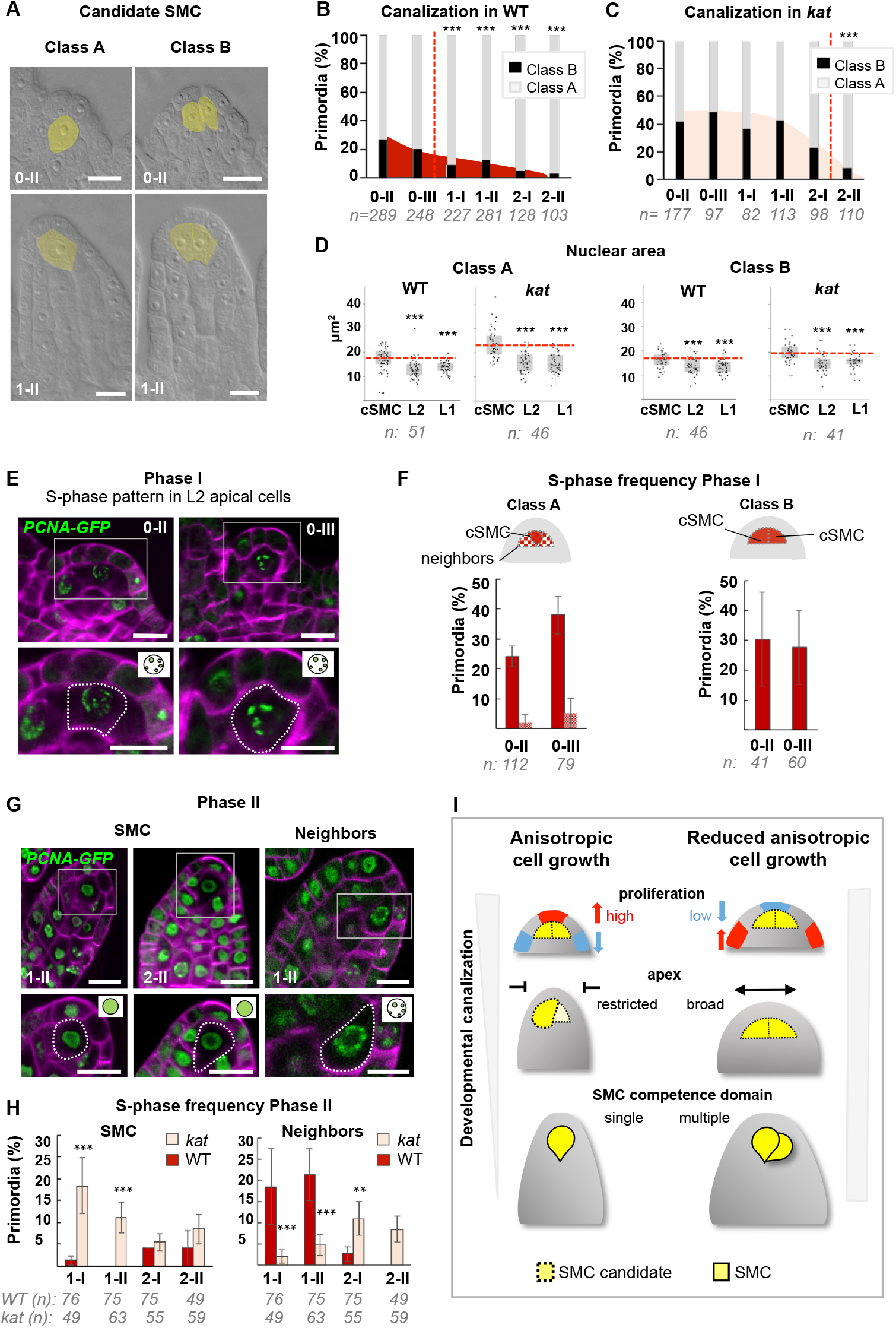
SMC singleness is progressively resolved during primordium growth. (A-B) In wild-type (WT) plants, ovule primordia harbour one (class A) or occasionally two (class B) SMC candidates with the frequency of class B gradually decreasing during development, reminiscent of developmental canalization. Typical images obtained by tissue clearing with SMC candidates highlighted in yellow (A) and plots showing the percentages of classes A and B (B). The frequency of class B ovules is significantly reduced from stage 1-I, suggesting that canalization (represented by the red dashed line) occur before that stage. The plot coloration is a visual aid only. (C) Developmental canalization is delayed in *katanin* mutants (*kat, mad5* allele). The proportion of class B ovules is significantly reduced from stage 2-II only. Quantifications for two additional mutants carrying *kat* alleles are presented Figure supplement 6A. The plot coloration is a visual aid only. (D) The candidate SMCs initially identified as enlarged cells consistently show an enlarged nucleus in both class A and class B primordia. Nuclear area is compared between the candidate SMCs (cSMC) and surrounding L2 and L1 cells (see Figure supplement 6D and Materials and Methods for further details). Box plots include jittered data points to visualize data variability. Red lines represent the median of cSMC nuclear area for comparison with other cell types. Equivalent quantifications for stage 0-III are presented in Figure supplement 6E. (E-F) During Phase I (stages 0-II, 0-III), S-phase is detected in candidate SMCs at higher frequency than in neighbors cells in class A ovules, and in both SMC candidates in class B ovules. Representative images of ovule primordia showing the speckled S-phase pattern of the *pPCNA1::PCNA1:sGFP* reporter (green) (E). Magenta signal: Renaissance SR2200 cell wall label. Scale bar 10μm. Quantification of speckled S-phase pattern in the SMC candidate and L2 neighbors (class A ovules) or in both SMC candidates (class B) (F). (G-H) During Phase II (stages 1-I to 2-II), in class A wild-type ovule primordium, SMC exits S-phase, and neighbors cells undergo S-phase; while *katanin* primordia show the opposite pattern. Representative images of class A ovule primordia primarily showing the nucleoplasmic pattern of *pPCNA1::PCNA1:sGFP* in SMCs at stages 1-II (left panel) and 2-II (middle panel); and of the speckled S-phase pattern in neighbors cells (right panel). Quantification of speckled S-phase pattern in SMC candidates and neighbors in Phase II wild-type and *katanin* primordia (I). Representative images and quantifications of Phase I *katanin* primordia and of class B primodia are presented in Figure supplement 6F-H. See also Figure 2H. (I) Model for the role of *KATANIN* in primordium growth and SMC differentiation (graphical abstract). In WT primordia, SMC differentiation follows a developmental canalization process influenced by cell growth anisotropy that shapes the primordium apex. In *kat* mutants, reduced anisotropy modifies the cell proliferation pattern, enlarges the apex and the L2,L3 apical domain, leading to multiple SMC candidates (delayed canalization). We propose that ovule primordium shape, controlled by anisotropic cell growth, determines SMC singleness. *Images*: scale bars : 10um. *Graphs*: n= total number of ovules. Error bar: standard error of the mean (F, I). Differences between cell types, domains, or genotypes as indicated in the graphs were assessed using *Wilcoxon signed rank test in* (*D*), *and* a two-tailed Fischer’s exact test in B, C, F, and I. P values: *p≤0.05, **p≤0.01, ***p≤0.001. Quantifications for additional alleles and detailed quantifications are provided in Figure supplement 6 and Source Data 6.

Next, we quantified class A and B primordia in *kat* mutants by scoring 2587 ovules of three different mutant alleles (Figure 6C, Figure supplement 6A). Clearly, plasticity is more strongly expressed with 42% (n=202) class B primordia at stage 0-II in *kat* compared to 26% (n=289) in the wild-type. In addition, the resolution process is delayed in *kat* mutant primordia as up to 33% class B primordia persist at stage 1-II (n=113) compared to 12% in the wild type (n=281). Consistently, the two SMC markers used previously, clearly identify ectopic SMC candidates in *kat* primordia at stage 1-I (Figure supplement 6B-C), and more significantly at stage 1-II (28.5% ectopic eviction of H1.1-GFP and 21% ectopic expression of KNU-YFP, Figure 5D-E). In addition, at stage 2-II, when 17% of cleared *kat* primordia (n=110) showed ectopic SMCs, H1.1-GFP eviction and KNU-YFP expression was only found in 8% (n=25) and 6.7% (n=74) of the primordia, respectively (Figure supplement 5B-C). Therefore, molecular events associated with SMC fate are partially uncoupled from cell growth during the resolution of SMC singleness in *kat*.

Another outcome of this study is the early emergence of SMC candidates, based on cell size mostly and confirming our former 3D cell-based analysis that showed increased cell volume of the L2,L3 apical cells already at stages 0-I/0-II (Figure 2J). To corroborate this finding using a different criterion, we measured the nuclear area, which is a distinctive feature of SMCs (Rodriguez-Leal et al., 2015; She et al., 2013). We compared the nuclear area of SMC candidate to that of surrounding L1 and L2 cells, in both class A and B primordia, at stages 0-II and 0-III. Wild-type class A and B primordia showed an enlarged nucleus in the candidate SMCs from stage 0-II onwards, and this correlation was also true in *kat* primordia (Figure 6D, Figure supplement 6D-E).

It remains however difficult to resolve the precise timing of SMC establishment at these early stages due to the lack of molecular markers. Yet, we rationalized that we may distinguish the SMC candidates from their neighbors by their cell cycle pattern, where cells entering meiosis may engage in a specific S-phase compared to regularly cycling mitotic cells. To this aim, we used a GFP-tagged PCNA variant marking the replication machinery, *pPCNA1::PCNA1:sGFP* (PCNA-GFP) (Yokoyama et al., 2016). When engaged at active replication forks, PCNA-GFP shows nuclear speckles characteristic of S-phase (Strzalka et al., 2011). During G1/G2, it remains in the nucleoplasm; and is undetectable in M-phase. We specifically quantified the distribution patterns of PCNA-GFP in cells from the L2 apical domain in wild type and *kat* primordia, separately for class A and B ovules (Figure 6E-H). In Phase I wild-type primordia, PCNA-GFP was always detectable, in both classes, indicating that L2 apical cells are rarely in M-phase, consistent with the seldom detection of mitoses using CYCB1.1db-GFP. We observed an S-phase pattern consistently in one (Class A) or more (Class B), centrally positioned L2 apical cells, presumably corresponding to SMC candidates (Figure 6E). This pattern was captured in a large proportion of primordia at stage 0-II and 0-III: ∼24% (n=112) and ∼40% (n=79) respectively for class A; 30% (n=41) and 28% (n=60) for Class B, respectively (Figure 6F). Such high frequencies could be generated either by a slow S-phase in SMCs candidate only (the persistence of the marker increasing the probability to score it repeatedly in our sample size), or by a regular (short) mitotic S-phase in a high number of SMC candidates. The low detection frequency of the mitotic marker CYCB1.1db-GFP in SMC candidates allow rejecting the latter possibility. Thus, the likeliest interpretation is that candidate SMCs enter a slow S-phase from early stages onwards, probably meiotic, although this cannot be assessed with this marker. In *kat* primordia at Phase I, Class A ovules showed a wild-type pattern in the SMC at stage 0-II, but a reduction at stage 0-III, suggesting alterations in S-phase entry; and class B ovules, by contrast, displayed high frequency of S-phase pattern, which indicate either longer S-phase or multiple dividing cells (Figure supplement 6G).

In Phase II, SMC candidates showed absence of S-phase pattern in a large majority of both Class A and B primordia (96,6% on average, over all Phase II) (Figure 6G-H, Figure supplement 6G-H) in agreement with the presence of newly replicated DNA at stage 1-I (She et al., 2013). By contrast with Phase I, however, S-phase is now detected in SMC neighbors (∼21% at stage 1-II) (Figure 6G-H) consistent with the divisions observed in these cells (Figure 2D). Strikingly in Phase II, *kat* primordia display a higher frequency of S-phase patterns in SMC candidates (Figure 6H, Figure supplement 6F,H), suggesting that S-phase duration or entry timing is delayed compared to wild type. SMC neighbors, by contrast, show a lower frequency of S-phase patterns (Class A), consistent with reduced division in these cells (Figure 4F).

Collectively, our data indicate a cellular heterogeneity in terms of size, nuclear size and S-phase patterns, of the L2 apical domain as compared to L1, which leads to the emergence of one or several SMC candidates as early as stage 0-II. The gradual decrease in the number of primordia with ambiguous SMC candidates demonstrates a developmentally-regulated resolution of SMC fate to a single cell. This process is associated with a specific cell-cycle progression, cellular elongation, and robust expression of SMC fate markers. In *kat* primordia displaying geometry alterations, SMC singleness is largely compromised: plasticity in SMC emergence is increased and fate resolution to a single SMC is delayed.

## Discussion

Organogenesis involves coordinated cell division and cell expansion, complex growth processes orchestrated by biochemical and mechanical cues (Echevin et al., 2019). How cell differentiation is coordinated in space and time during organ growth and whether these processes are interrelated are central aspects for the elucidation of patterning principles (Whitewoods et al., 2017). The female germline is initiated with SMC differentiation in the ovule primordium. The SMC emerges as a large, elongated subepidermal cell that is centrally located at the apex of the primordium, a digit-shaped organ emerging from the placenta. To study how SMC fate relates to ovule organogenesis, we generated a reference collection of images capturing ovule primordium development at cellular resolution in 3D and determined cell division frequencies in the different domains. We observed a biphasic pattern of cell divisions alternating in the epidermis and inner layers, as well as the apical and basal domains, in Phase I (stages 0-I to 0-III) and Phase II (stages 1-I to 2-II).

However, this approach did not allow us to resolve the driving morphogenetic factors. For this reason, we developed continuous and cell-based 2D simulations of primordium growth. The different simulations revealed key growth principles shaping the ovule primordium and uncovered differential roles for the epidermis and inner layers. Notably, an inner tissue-driven growth model, where the L1 also contributes expansion of the primordium, best described ovule primordium growth. This is reminiscent of the possible model describing leaf primordium emergence (Peaucelle et al., 2011). In addition, best-fit models produced by both cell-based and FEM simulations predicted a growth-promoting signal in a confined domain along a vertical stripe at primordium emergence. Candidates growth signals are phytohormones, peptides, and small RNAs known to affect ovule primordium growth (Kawamoto et al., 2020; Pinto et al., 2019; Su et al., 2020). The domain of auxin response restricted in the L1 dome and of cytokinin signaling localized in a region basal to the SMC in Phase II primordia (Bencivenga et al., 2012) also suggest a confined growth signal. Whether the signaling domains are established already in Phase I and plays a causative role in primordium patterning remains to be determined. The epidermis, by contrast, is predicted to play a key role in accommodating the constraints generated by inner-tissue growth. In this layer, passive strain-based growth and anisotropic material properties possibly resolve mechanical conflicts arising between tissue layers that grow at different rates (Hervieux et al., 2017). In line with this hypothesis, we observed frequent divisions in L1 apical cells *in vivo* that support the expansion of the epidermis while inner-tissues develop.

Interestingly, while our models did initially not contain an *a priori* rule to produce the typical, elongated shape of the SMC, it emerged consistently as a trapezoidal-shaped L2 cell in the cell-based reference model. This shape likely emerges from the combination of assigned anisotropic cell growth and geometrical constraints imposed by the surrounding growing tissues. Explicitly blocking SMC division during the simulation not only enabled its expansion as expected, but also pushed surrounding cells and strongly deformed ovule morphology. Thus, ovule growth homeostasis *in vivo* likely requires a mechanism to accommodate the differential growth of the SMC. A gradual reduction of SMC turgor pressure is a plausible scenario that would limit SMC size and prevent overriding the constraints provided by surrounding cells, similarly to what was suggested for the shoot apical meristem (Long et al., 2020). In turn, a gradient of pressure in field of cells could provide positional information through the directional movement of water and other molecules thereby linking organ growth homeostasis, cell growth, and cell fate (Beauzamy et al., 2014; Long et al., 2020). Such a mechanism could also participate in determining the domain of the growth-signal predicted by our simulations.

Another prediction of our growth models is the key role of anisotropic cell growth in controlling the geometry of the ovule primordium. Primordia of the *katanin* (*kat*) mutant, deficient in the microtubule severing enzyme KATANIN (Luptovciak, Komis, et al., 2017) resemble virtual mutant primordia generated by models where cell growth is isotropic in the inner layers. *kat* primordia have a flatter dome and large basis associated with global alterations in cell proliferation pattern, cell size and cell shape. Interestingly, *kat* primordia develop ectopic SMC candidates as early as stage 0-II. The most parsimonious hypothesis is that the altered geometry in *kat* primordia expands the domain of cells competent to form SMCs. Yet, we cannot exclude a direct effect of the *kat* mutation on L2 apical cells, disconnected from organ geometry, which would induce *de novo* SMC fate. However, this scenario is unlikely because we would expect increased cell growth and slower mitoses (Luptovciak, Komis, et al., 2017), resulting in a reduced division frequency of L2 apical cells, which is not the case. Instead, we measured increased cell divisions in L2 SMC neighbors at stage 0-II. Also, the delayed divisions of SMC neighbors in *kat* at stage 1-II cannot explain the formation of ectopic SMC candidates at stage 0-II. Therefore, the most parsimonious explanation is that the emergence of several SMC candidates in *kat* primordia is an effect of ovule primordium geometry. In this scenario, isotropic cell growth and altered mechanical constraints in the tissue acting from primordia emergence onwards, lead to divisions and patterning alterations expanding the domain of cells competent for SMC fate. This working model paves the way to explore the role of the epidermis geometry in controlling regulators of cell cycle and SMC fate in L2 apical cells. This is reminiscent to the shoot apical meristem epidermis acting on dome shape and stem cells regulators in underlying layers (Gruel et al., 2016; Savaldi-Goldstein et al., 2007). The predicted role of the primordium epidermis to accommodate - and perhaps feedback on – underlying growth constraints is particularly interesting considering the role of mechanical cues on gene regulation (Fal et al., 2017; Landrein et al., 2015). The epidermis of the ovule primordium is also a known source of signaling cues (Kawamoto et al., 2020; Pinto et al., 2019; Su et al., 2020). In addition, phytohormones act themselves on KATANIN-mediated oriented cell growth and cell division (Luptovciak, Komis, et al., 2017). In this context, it is possible to speculate that misshaping of *kat* primordia may also arise from a disrupted feedback altering the distribution pattern of the hypothesized growth signals.

Furthermore, our analyses unveiled new characteristics of the SMC establishment process. We found that SMC candidates emerge from within a mitotically quiescent L2 apical domain, consistent with the finding that the H3.1 histone variant HTR13 is evicted, marking cell cycle exit (Hernandez-Lagana et al., 2020). In addition, SMC candidates have a markedly long S-phase compared to surrounding cells. These observations are reminiscent of the animal germline where mitotic quiescence is a prerequisite to meiosis (Kimble, 2011; Reik et al., 2015). Also, early SMC candidates already display a typically large cell and nucleus size and an elongated shape aligned with the primordium growth axis. Collectively, we found that SMC characteristics are established much earlier than previously thought, i.e. soon after primordium emergence (stages 0-II /0-III). Moreover, these characteristics frequently arise in more than one SMC candidate at Phase I, resolving into a single SMC at the onset of Phase II. This clearly documents a developmental sequence of plasticity at SMC fate emergence and progressive resolution of SMC fate in a single cell. This is reminiscent of developmental canalization, which refers to the capacity of an organism to follow a given developmental trajectory in spite of disturbances (Hallgrimsson et al., 2019; Scharloo, 1991; Waddington, 1942). Cell fate canalization is well studied in animal systems where it is modulated by intercellular signal-based feedback mechanisms (Heitzler et al., 1991), epigenetic regulation (Pujadas et al., 2012) and organ geometry (Huang et al., 2020; Huang, 1992; Pujadas et al., 2012; Royer et al., 2020). In plants, canalization is better known in the context of organogenesis during phyllotaxis and developmental robustness (Godin et al., 2020; Lempe et al., 2013). Our study expands the examples of a canalization process at the level of a cellular domain, in the Arabidopsis ovule primordium. While L2 apical cells initially share the competence to form SMC candidates, leading to plasticity at SMC emergence, the progressive restriction of cell fate possibilities in the primordium apex ultimately leads to only one SMC committed to meiosis. Our results are in line with a formerly proposed canalization process operating during SMC establishment (Grossniklaus et al., 1998; Rodriguez-Leal et al., 2015). Despite the fact that several mutations (Pinto et al., 2019; Mendes et al., 2020), including *kat* (this study), alter SMC singleness, canalization remains a robust process securing the formation of a single embryo sac for most of these genetic perturbations in Arabidopsis. How this developmental mechanism buffers phenotypic inter-individual variations and whether it is evolutionary constrained remains to be determined.

In this study, we quantified plasticity among ovule primordia and progressive fate resolution during primordium growth in wild-type reference accessions. We characterized specific cellular events associated with these processes, notably differential cell growth and cell division. SMC fate emergence is characterized by mitotic quiescence and cellular growth in one or more L2 apical cells. SMC singleness resolution is associated with re-entry in a somatic cell cycle (this study), and re-incorporation of a replicative histone H3.1 (Hernandez-Lagana et al., 2020) of initial candidates neighboring the SMC. How known epigenetic and signaling factors interplay to secure SMC singleness remains to be determined. Similar to mouse and Drosophila, where tissue mechanics and organ geometry were shown to contribute to cell fate canalization (Chan et al., 2017; Huang et al., 2020; Royer et al., 2020), we propose that ovule primordium geometry contributes to channel SMC fate in the apex and to the resolution of a single SMC. In this conceptual framework, *kat* increases plasticity and delays the resolution process towards SMC singleness (working model Figure 6I).

Altogether our work proposes a conceptual framework linking organ geometry, cell shape and cell fate acquisition in the ovule primordium, potentially of broader relevance in plant patterning. In addition, the image resource published in this study is complementary to others capturing ovule development at later stages (Lora et al., 2017; Vijayan et al., 2020). It also populates a growing number of 3D-segmented images of plant tissues and organs (Wolny et al., 2020), which collectively build the fundaments to developmental atlas integrating morphogenesis with gene expression (Hartmann et al., 2020).

## Materials and Methods

### Key Resources Table

| REAGENT or RESOURCE | SOURCE | IDENTIFIER |
| --- | --- | --- |
| **Chemicals** |  |  |
| SCRI Renaissance Stain 2200 | Renaissance Chemicals | http://www.renchem.co.uk/index.php/specialty-chemicals-division/item/48-selected-fluorescent-dyes-and-brighteners-for-microscopists |
| Propidium Iodide (mPS-PI staining solution) | Sigma- Aldrich | Catalog # P4170 |
| Sodium metabisulphite (mPS-PI staining solution) | Sigma- Aldrich | Catalog # S9000 / PubChem: 329824616 |
| **Deposited Data** |  |  |
| Images dataset 3D segmentation ovule primordium wild-type (*Col-0*, *Ws-4*) and *katanin* mutant (*bot1-7* allele) | This paper, *submission to IDR repository underway* | https://idr.openmicroscopy.org/ |
| **Experimental Models: Organisms/Strains** |  |  |
| *Arabidopsis thaliana*, ecotype *Col-0* | NASC | NASC ID: N22625 |
| *Arabidopsis thaliana*, ecotype *Ws-4* | NASC | NASC ID: N5390 |
| *Arabidopsis thaliana*, *bot1-7* mutant | H.Höfte, (Bichet et al., 2001) |  |
| *Arabidopsis thaliana*, *mad5* mutant | O.Voinnet (Brodersen et al., 2008) |  |
| *Arabidopsis thaliana*, *lue1* mutant | (Bouquin et al., 2003); NASC | NASC ID: N57954 |
| *Arabidopsis thaliana*, *promCYCB1.1::CYCB1.1db-GFP* | M.Bennett (Ubeda-Tomas et al., 2009) |  |
| *Arabidopsis thaliana*, *promH1.1::H1.1-GFP* | (She et al., 2013) |  |
| *Arabidopsis thaliana*, *promKNU::nls-YFP* | M.Tucker (Tucker et al., 2012) |  |
| *Arabidopsis thaliana*, *promAtDMC1::AtDMC1-GUS* | I.Siddiqi (Klimyuk et al., 1997) |  |
| *Arabidopsis thaliana*, *promWOX2::CenH3-GFP* | N.DeStorme (De Storme et al., 2016) |  |
| *Arabidopsis thaliana*, *promPCNA1::PCNA1-GFP* | S. Matsunaga (Yokoyama et al., 2016) |  |
| **Software and Algorithms** |  |  |
| IMARIS Cell | Bitplane, AG | https://imaris.oxinst.com/ |
| ExportImarisCells, plugin for IMARIS | This paper | https://github.com/barouxlab/ExportImarisCells |
| MorphoGraphX | Freely available | http://www.mpipz.mpg.de/MorphoGraphX |
| MorphoMechanX | Freely available | https://www.mpipz.mpg.de/MorphoGraphX/MorphoMechanX/ |
| MassSpringModels_ovuleGrowth2D, plugin for MorphoMechanX | This paper | https://github.com/GabriellaMosca/MassSpring_2DovuleGrowthModel |
| FEM_2DOvuleGrowthModel, plugin for MorphoMechanX | This paper | https://github.com/GabriellaMosca/FEM_2DOvuleGrowthModel |
| 3DAutoLabeling-ShapeQuant, plugin for MorphoMechanX | This paper | https://github.com/GabriellaMosca/3DAutoLabeling-ShapeQuant |
| OvuleViz, R-based data plotting interface | This paper | https://github.com/barouxlab/OvuleViz |
| R / RStudio | Freely available | https://www.r-project.org/<br>http://www.rstudio.com/ |
| OMERO | Freely available | https://www.openmicroscopy.org/omero/ |
| ImageJ / FIJI | Freely available | https://imagej.net/Fiji |

### Plant growth and plant material

*Arabidopsis thaliana* plants were grown under long-day conditions (16 hours light) at 20-23 C in a plant growth room. Columbia (Col-0) and Wassileskija (Ws-4) accessions were used as wild-type controls depending on the mutant background used in the experiment. Three *katanin* alleles were used: *bot1-7* (Bichet et al., 2001) in Ws-4 accession, *lue1* (Bouquin et al., 2003) and *mad5* (Brodersen et al., 2008) both in the Columbia (Col-0) accession. Homozygous mutant individuals for all the *katanin* alleles were identified on the basis of their recessive vegetative phenotype. The following published markers were used: pCYCB1.1:db-GFP (Ubeda-Tomas et al., 2009), pKNU:nls:YFP (Tucker et al., 2012), pH1.1:H1.1:GFP (She et al., 2013), AtPCNA1:sGFP (Yokoyama et al., 2016), pAtDMC1:GUS (Agashe et al., 2002; Klimyuk et al., 1997), pWOX2:CenH3:GFP (De Storme et al., 2016), and crossed to *katanin* mutants, and to Ws-4 ecotype for *bot1-7* allele comparisons.

### 3D imaging and image processing (segmentation and labeling)

Entire carpels were stained using the pseudo-Schiff propidium iodide (PS-PI) cell wall staining procedure providing excellent optical transparency for 3D imaging in depth in whole-mount. We described previously the manipulation, staining, mounting of the flower carpels and imaging procedures (Mendocilla-Sato, 2017). Cell-boundary based image segmentation was done using ImarisCell (Bitplane) as described in details previously (Mendocilla-Sato, 2017). Each ovule was manually labelled in Imaris using customized Cell Labels for the different cell types and domains colored as shown in Figure 1. We defined the labels as follows:

SMC (Spore Mother Cell): most apical central enlarged L2 cell. At stage 0-I, as enlargement is not always detected visually, the most apical L2 cell was then selected as candidate SMC (cSMC).

- L1: epidermal cells
- L1 dome: most apical cells in contact with SMC
- L2/L3: cells below the epidermis. L2 and L3 were not distinguished originally.
- Apical domain: group of cells at the apex of the primordium and encompassing the SMC and direct neighbor cells.
- Basal domain: group of cells at the basis of the primordium below the apical domain and until, but not including cells of the placental surface. At stages 0-I and 0-II only an apical domain is defined. A basal domain appears only starting stage 0-III.
- CC (Companion Cell): L2 cells in apical domain in contact with the SMC with an elongated shape (as
- judged in ovule longitudinal median section using the “clipping plane” IMARIS tool).
- SMC contact: cells in contact with the SMC.

Semi-automated segmentation requiring user input and manual labelling can be error prone. To reduce the error rate, the 92 images were segmented and labeled by one author, but verified and curated by two others. All Imaris files used in the study will be available at the IDR repository (www.idr.org).

### Ovule stage classification

Ovule development is described according to a well-accepted nomenclature (Schneitz, 1995). The first stage, initially defined as stage 1-I, indistinctly grouped primordia from emergence until digit shape. To enable describing early morphogenetic processes, however, we propose to (i) restrict stage 1-I to the final digit shape stage and (ii) subdivide preceding stages as stages 0-I, 0-II and 0-III, as shown in Figure 1B. These developmental stages are classified according to the approximate number of cell layers protruding above the placenta and overall shape of the ovule as described in Table 1 and Source Data 1.

### Quantification of cell number, size and shape and interactive plotting using OvuleViz

Several cell descriptors were retrieved using the ImarisCell’ Statistics function and exported as .csv files: cell area, cell volume, cell sphericity, cell ellipticity (oblate and prolate). We developed an interactive R-based data plotting interface, OvuleViz, reading the Imaris-derived data within the exported files ordered by genotype then stages (Figure supplement 1). OvuleViz is freely available at https://github.com/barouxlab/OvuleViz, and is based on a *shiny* interface for R. OvuleViz allows plotting selectively one or several of the cell descriptors for chosen stages and genotypes, along different visualization (scatter plots, box plots, histograms). In addition, OvuleViz retrieves the cell number from the number of objects in the .csv file.

### 3D quantification of ovule volume and shape using IMARIS

Ovule volume and shape were quantified on 3D segmentations using IMARIS software, to compare *katanin* and wild-type genotypes. For each ovule, all segmented labeled cells were duplicated and fused as a single cell object. The *ImarisCell’ Statistics* function was used to retrieve “cell volume” and “bounding box OO” (object oriented 3D bounding rectangle, exporting the minimum (A), mid (B) and maximum (C) lengths of the object bounding rectangle). Width to Height ratio was calculated by dividing the maximum (C) length by the mean of mid (B) + minimum (A) lengths. Bounding box occupancy was calculated as the ratio of ovule volume by the bounding box volume.

### Modeling and image analysis with MorphoMechanX

The details of tissue growth models and MorphoMechanX-based processing are described in Supplemental File 1.

### 2D cytological analysis of cleared ovule primordia and quantifications

For cytological examination of cleared ovule primordia, flower buds from wild-type and mutant plants were harvested and fixed in formalin-acetic acid-alcohol solution (40% formaldehyde, glacial acetic acid, 50% ethanol; in a 5:5:90 volume ratio) for at least 24h at room temperature. After fixation, samples were washed two times with 100% ethanol and stored in 70% ethanol. Gynoecia of 0.2–0.6 mm in length were removed from the flowers with fine needles (1 mm insulin syringes), cleared in Herr’s solution (phenol : chloral hydrate: 85% lactic acid : xylene: clove oil in 1:1:1:0,5:1 proportions), and observed by differential interference contrast microscopy using a Zeiss Axioimager Z2 microscope and 40X or 60X oil immersion lenses. Picture were acquisition was done with a sCMOS camera (Hamamatsu ORCA Flash V2). Nuclei area measurements were carried out with ImageJ software, using the manual contour tool “Oval”.

### Fluorescence microscopy and quantifications

Imaging of ovule primordia stained in whole-mount for cell boundary was done as described (Mendocilla-Sato, 2017) using a laser scanning confocal microscope Leica LCS SP8 equipped with a 63X glycerol immersion objective and HyD detectors.

Imaging of the GFP and YFP markers was performed using a laser scanning confocal microscope Leica LCS SP8 equipped using a 63X oil immersion objective and HyD detectors. Samples were mounted in 5% glycerol with the cell wall dye Renaissance 2200 (SR2200) diluted 1/2000. The following wavelengths were used for fluorescence excitation and detection: Renaissance: excitation 405nm, and detection 415–476 nm; GFP: excitation 488m and detection 493-550 nm; YFP: excitation 514nm and detection 590-620nm. Channels contrast and intensity were adjusted using ImageJ or OMERO.

The mitotic activity in both wild-type and *katanin* ovules was quantified by scoring the cells expressing *promCYCB1::dbCYCB1-GFP* (M phase reporter) on 3D stacks, in each ovule domain at each developmental stage. Only ovules showing at least one cell expressing *promCYCB1::dbCYCB1-GFP* were included in the analysis. At a given stage, the frequency of mitoses per domain is the ratio between the number of GFP positive cells within a domain in all observed ovules, to the total number of GFP positive cells found in all domains in that population of ovules.

To quantify ectopic expression of H1.1-GFP, KNU-YFP and pWOX2:CenH3-GFP markers, the percentage of ovules presenting expression (or eviction in the case of H1.1-GFP) of the marker in more than one cell was scored on 3D stacks. To quantify PCNA-GFP patterns, using 3D stacks, cells of the L2 apical domain were classified according to the “speckled” or “nucleoplasmic” patterns, or absence of the marker.

### Histochemical detection of *uidA* reporter gene product (GUS staining)

Gynoecia of 0.4–0.6 mm in length were removed from the flowers with fine needles and placed in staining solution, using high stringency conditions for ferro- and ferricyanide concentrations to limit GUS product diffusion (0.1% Triton X-100, 10 mM EDTA, 5 mM ferrocyanide, 5 mM ferricyanide and 20mg/ml 5 -bromo-4-chloro-3-indolyl-beta-d-glucuronic acid cyclohexyl-ammonium salt (X-gluc, Biosynth AG, Staad, CH) in 50 mM phosphate buffer), for 96h at 37°C. After staining, the samples were mounted in clearing solution (50% glycerol, 20% lactic acid diluted in water) and observed by differential interference contrast microscopy using a Zeiss Axioimager Z2 microscope. Picture acquisition was done with a color sCMOS camera (Axiocam 503 color Zeiss).

### Statistical analysis

To identify the main cellular descriptors – cell area, cell volume, sphericity, ellipticity oblate, ellipticity prolate – explaining variance of each cell types, at each ovule primordium growth stages and between genotypes, we used principal component analysis (PCA) on a cells’ descriptors matrix. PCA was executed using the R software version 3.6.3. In all different subsets of data, PCA was performed by singular value composition of the centred and scaled data matrix. Data entries with missing values were removed before analysis. All Cell data as exported by Imaris were uploaded but only cells with a ‘Cell Label’ as described in the method section ‘3D imaging’ were plotted. For visualization of PCAs, only the first two principal components were represented in both score and loading plots.

To determine if the means between two datasets were significantly different we used two-tailed t-test when the data were normally distributed (n> 30). For all datasets with n <30, we assume that normality was not possible to assess properly (as small samples most often pass normality tests), thus we used non parametric tests. Wilcoxon Signed-Rank two-tailed test was used for paired quantifications, and Mann Whitney U two-tailed test was used for unpaired quantifications. Tests were performed in Excel or in R (wilcox.test function). To compare ovule proportions, we used two-tailed Fischer’s Exact test, using https://www.langsrud.com/fisher.html, available online. Variability was assessed using the Standard Error of the mean (SE), indicated in the graphs and/or the supplemental data when applicable. For some datasets, boxplots were used to improve visualization of data distribution. The number of samples and biological replicates for each experiment are indicated in the figure and/or figure legends and/or supplemental data.

## Author contributions

CB, DA, GM, EHL, DG, OH, AB, CG: Conceptualization; EHL, GM, CB, DA, OH, AB, CG, DG: Methodology; GM, NP, AG: Softwares; EHL, GM, EMS, DA, CB: Validation; EHL, GM, AG, DA, CB: Formal analysis; EHL, EMS, DA, CB: Investigation; EMS, AF, OH: Ressources; EHL, GM, NP, DA, CB: Data curation; EHL, GM, EMS, DG, DA, CB: Writing – Original Draft; EHL, GM, OH, AB, CG, UG, DG, DA, CB: Writing – Reviewing and Editing; EHL, GM, DA, CB: Visualization; CB, DA, DG, OH, AB, UG: Supervision; CB, DA, DG: Project administration; CB, DA, DG, OH, AB, CG, UG: Funding Acquisition. All authors reviewed and revised the manuscript.

## Data availability

All materials, scripts and datasets generated and analyzed in the current study are available at repositories (see Materials and Methods) or from the corresponding authors upon reasonable request.

## Acknowledgments

We thank S. Strauss (Max Planck Institute for Plant Breeding Research, Cologne, Germany) and B. Lane (John Innes Centre, Norwich, U.K.) for providing code and assessment for mesh generation and simulations; S. Guyer (Bitplane, Switzerland) and M. Lartaud (CIRAD, France) for their help with IMARIS analyses; O. Leblanc (IRD Montpellier, France) for his help with data visualization in R; the Montpellier imaging facility MRI, member of the national infrastructure France-BioImaging supported by the French National Research Agency (ANR-10-INBS-04) and the Center for Microscopy and Image Analysis of the University of Zurich (ZMB) for microscopy imaging infrastructures; M. Ueda (Nagoya University, Japan), I. Siddiqi (Center for Cellular and Molecular Biology, India), N. de Storme (Katholieke Universiteit Leuven, Belgium), S. Matsunaga (Tokyo Science University, Japan), M. Tucker (University of Adelaide, Australia), and the NASC stock center for seed germplasms.

This work was funded by the University of Zürich, the IRD, a PRCI grant from the Agence Nationale de la Recherche/Swiss National Science Foundation (#ANR-16-CE93-0002) to CB and DA; grants from the Swiss National Science Foundation (# 310030B_160336 to UG, # IZCOZ0_182949 to CB), from the Commission for Technology and Innovation (CTI grant #16997) to CB, from the Baugarten Stiftung Zürich to CB, a CONACYT fellowship (# 438277) to EHL, and a fellowship from the Forschungskredit of the University of Zurich (#FK-74502-04-01) to GM.

## Conflict of interest

The authors declare no conflict of interest.

## Notes

### Competing Interest Statement

The authors have declared no competing interest.

### Summary of Updates

Text revised and Supplemental figures updated

