## Supplementary material for "Organ geometry channels cell fate in the Arabidopsis ovule primordium": Figures Supplements

### OvuleViz, an interactive interface enabling dynamic analyses of segmented data

#### A. Plotting 'à la carte'

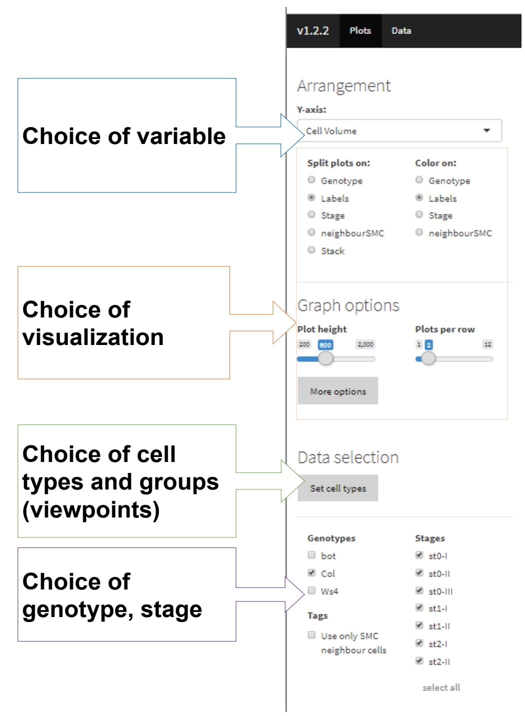


#### B. Plot types

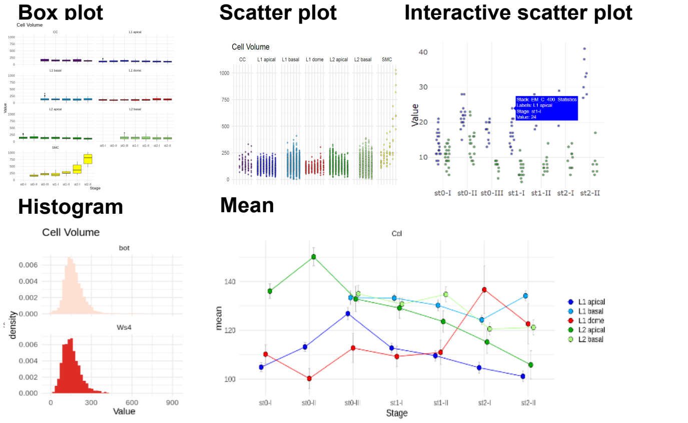


#### C. Export data tables

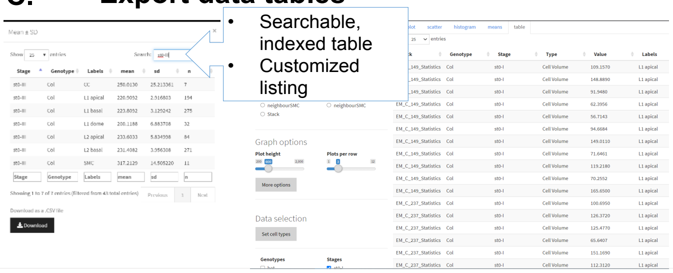


### Principal Component Analyses

#### D. All

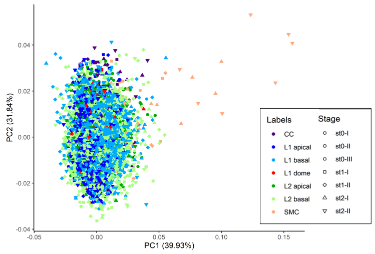


| All | PC1 | PC2 | PC3 | PC4 | PC5 |
|---|---|---|---|---|---|
| E*(oblate)* | -0.15 | -0.698 | 0.16 | -0.682 | -0.006 |
| E*(prolate)* | 0.203 | 0.671 | 0.215 | -0.68 | -0.009 |
| Vol | 0.685 | -0.175 | -0.089 | 0.014 | -0.702 |
| Area | 0.683 | -0.179 | 0.102 | 0.05 | 0.699 |
| Sph | 0.03 | 0.031 | -0.954 | -0.264 | 0.137 |
| Std Dev | 1.413 | 1.261 | 1.034 | 0.574 | 0.116 |
| p Var | 0.399 | 0.318 | 0.214 | 0.066 | 0.003 |
| Cumul. | 0.399 | 0.717 | 0.931 | 0.997 | 1 |

E, Ellipticity; Vol, volume; Sph, sphericity; Std Dev, standard deviation; p Var, proportion variance, Cumul., cumulative proportion

#### E. per stage

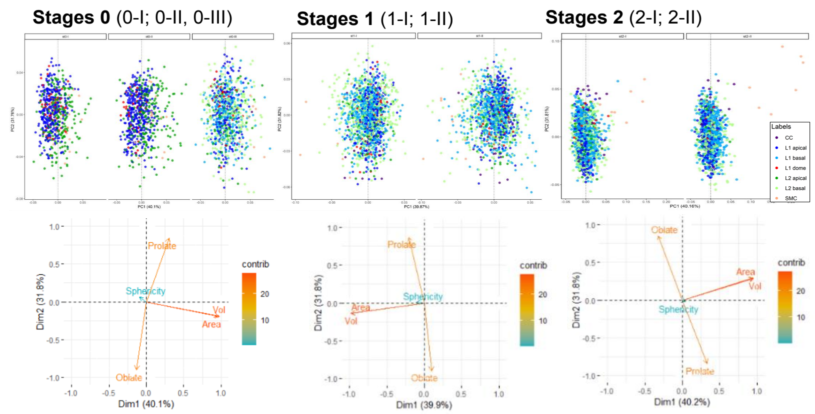


| Per Stage | PC1 | PC2 | PC3 | PC4 | PC5 | PC1 | PC2 | PC3 | PC4 | PC5 | PC1 | PC2 | PC3 | PC4 | PC5 |
|---|---|---|---|---|---|---|---|---|---|---|---|---|---|---|---|
| E*(oblate)* | -0.093 | -0.712 | -0.176 | -0.673 | -0.006 | 0.069 | -0.709 | 0.178 | 0.679 | 0.000 | -0.226 | 0.673 | 0.159 | 0.686 | 0.008 |
| E*(prolate)* | 0.215 | 0.667 | -0.224 | -0.677 | -0.009 | -0.146 | 0.689 | 0.205 | 0.680 | 0.007 | 0.231 | -0.665 | 0.193 | 0.683 | 0.008 |
| Vol | 0.681 | -0.147 | 0.174 | 0.022 | -0.696 | -0.699 | -0.110 | -0.052 | -0.030 | 0.705 | 0.670 | 0.227 | -0.076 | 0.008 | 0.702 |
| Area | 0.691 | -0.154 | -0.034 | 0.07 | 0.702 | -0.693 | -0.106 | 0.130 | -0.073 | -0.697 | 0.667 | 0.231 | 0.107 | -0.023 | -0.700 |
| Sph | -0.067 | 0.047 | 0.942 | -0.288 | 0.151 | -0.075 | 0.007 | -0.952 | 0.266 | -0.132 | 0.030 | -0.015 | -0.959 | 0.247 | -0.131 |
| Std Dev | 1.416 | 1.26 | 1.041 | 0.564 | 0.08 | 1.412 | 1.261 | 1.032 | 0.584 | 0.098 | 1.417 | 1.261 | 1.031 | 0.566 | 0.137 |
| p Var | 0.401 | 0.318 | 0.217 | 0.064 | 0.001 | 0.399 | 0.318 | 0.213 | 0.068 | 0.002 | 0.402 | 0.318 | 0.213 | 0.064 | 0.004 |
| Cumul. | 0.401 | 0.719 | 0.935 | 0.999 | 1 | 0.399 | 0.717 | 0.930 | 0.998 | 1 | 0.402 | 0.720 | 0.932 | 0.996 | 1 |

**Figure Supplement 1_Related to Figure 1. Approaches for morphodynamic analyses of ovule primordium development using a reference set of 3D segmented images.**

**Figure Supplement 1_Related to Figure 1. Approaches for morphodynamic analyses of ovule primordium development using a reference set of 3D segmented images.**

(A) OvuleViz, an interactive interface enabling the analysis of segmented data based on exported cell statistics. The different snapshots display several features of OvuleViz, enabling the generation of plots from chosen cell statistics for chosen ovule primordium stage, genotype, cell labels or groups thereof (viewpoints).

(B) Examples of plot types that can be generated by OvuleViz.

(C) Possibility to export all or a subset of data as summary table or raw dataframe for further analyses external to OvuleViz.

(D) PCA of cell descriptors from the wild-type reference dataset (*Arabidopsis* Col-0) (see Source Data 1), for all cell types and all stages. PCA was performed on a dataset containing Sphericity, Ellipticity oblate, Ellipticity prolate, Area, and Volume. Only the first and the second Principal Components (PCs) are shown. The loading plot and table indicate the variables' contribution to each component.

(E) Same as (D) for all cell types grouped by stage: stage 0 (0-I; 0-II), stage 1 (1-I; 1-II), and stage 2 (2-I; 2-II). Variables' contributions are presented by loading plots for each PC and summary table.

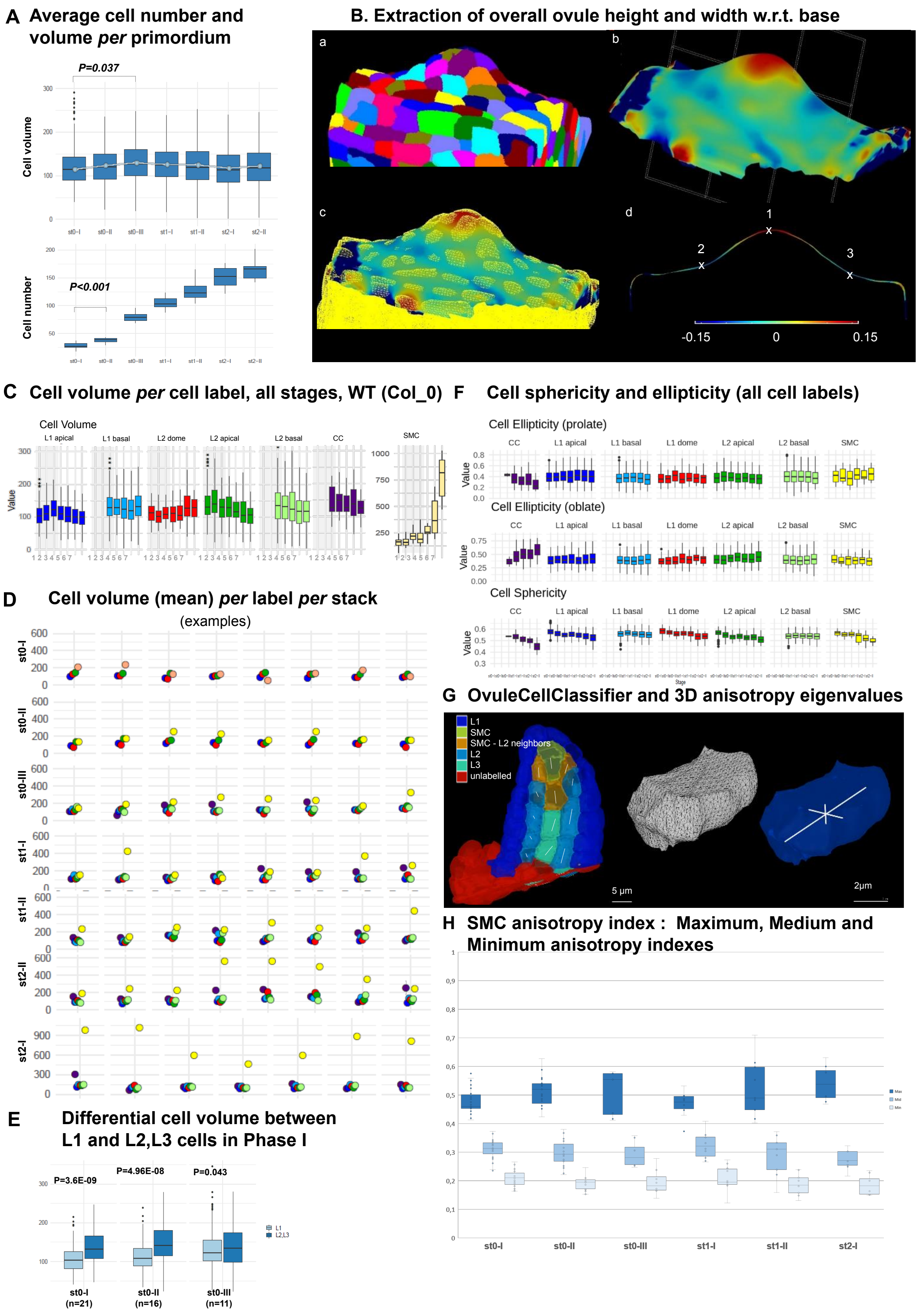


**Figure supplement 2_Related to Figure 2. Ovule primordium morphogenesis involves domain-specific cell division and anisotropic cell growth patterns.**

**Figure supplement 2_Related to Figure 2. Ovule primordium morphogenesis involves domain-specific cell division and anisotropic cell growth patterns.**

(A) Average cell number and cell volume in wild-type (Col) ovule primordium during development. Mann Whitney U test was used to assess the difference of each stage relative to the previous stage. Related to Figure 2A.

(B) Extraction of overall ovule height and width used in Figure 2B,C. (a) segmented stack imported in MorphoGraphX, (b) continuous surface mesh created in MorphoGraphX with the minimal curvature projected, the clipping plane is shown (dashed grid), (c) comparison between cell-mesh and continuous mesh, (d) Longitudinal section of the mesh showing maximal (1) and minimal (2 and 3) curvature points. Heatmap: minimal curvature $mm^{-1}$. Related to Figure 2B-C.

(C) Box plots and line plot (error bar is standard error of the mean) showing the mean cell volume in each ovule domain during all developmental time points. Mann Whitney U test was used to assess the difference between stages (L1 apical). Related to Figure 2I.

(D) Scatter plots comparing the cell volume of SMC (yellow) to the average cell volume of other cells types (same color code as in Figure 1) *per* ovule primordium, per stage. Related to Figure 2I.

(E) Comparison of cell volume between L1 and L2L3 cells in Phase I ovule primordia (stages 0-I to 0-III). Wilcoxon Signed Rank test was used to assess the difference between domains. Related to Figure 2J.

(F) Quantification of cell sphericity and ellipticity (prolate and oblate, see Materials and Methods) at all ovule primordium domains during development.

(G) Computation of covariance matrix eigenvalues for cell anisotropy analysis: Left: semi-automated cell type classification showing labelled cells along a longitudinal cut of an ovule. L2 SMC neighbors (orange) stands for all L2 cells sharing a portion of their wall with the SMC (yellow). Unlabelled cells (red) are not considered in the analysis. For the inner cells, the main anisotropy axis (the eigenvector of the covariance matrix corresponding to the biggest eigenvalue) is shown with a white line. Right: representation of eigenvectors of the covariance matrix (white lines) used for the PCA analysis on cell anisotropy on a random cell. The eigenvectors are scaled by their own eigenvalue normalized over the sum of the three eigenvalues. The longest eigenvector is aligned with the longest cell axis, the shortest with the shortest cell axis and the mid eigenvector with the cell axis orthogonal to the other two (and of intermediate length). Related to Figure 2K. See also movie in Supplemental File 2 and Supplemental File 1.

(H) SMC anisotropy index calculated from Maximal (dark blue), Mid (mid blue) and Minimal (pale blue) eigenvalues, normalized by the sum of all, from stage 0-I to stage 2-I.Related to Figure 2K. P values: *$p \leq 0.05$, **$p \leq 0.01$, ***$p \leq 0.001$. See also Source Data 2.

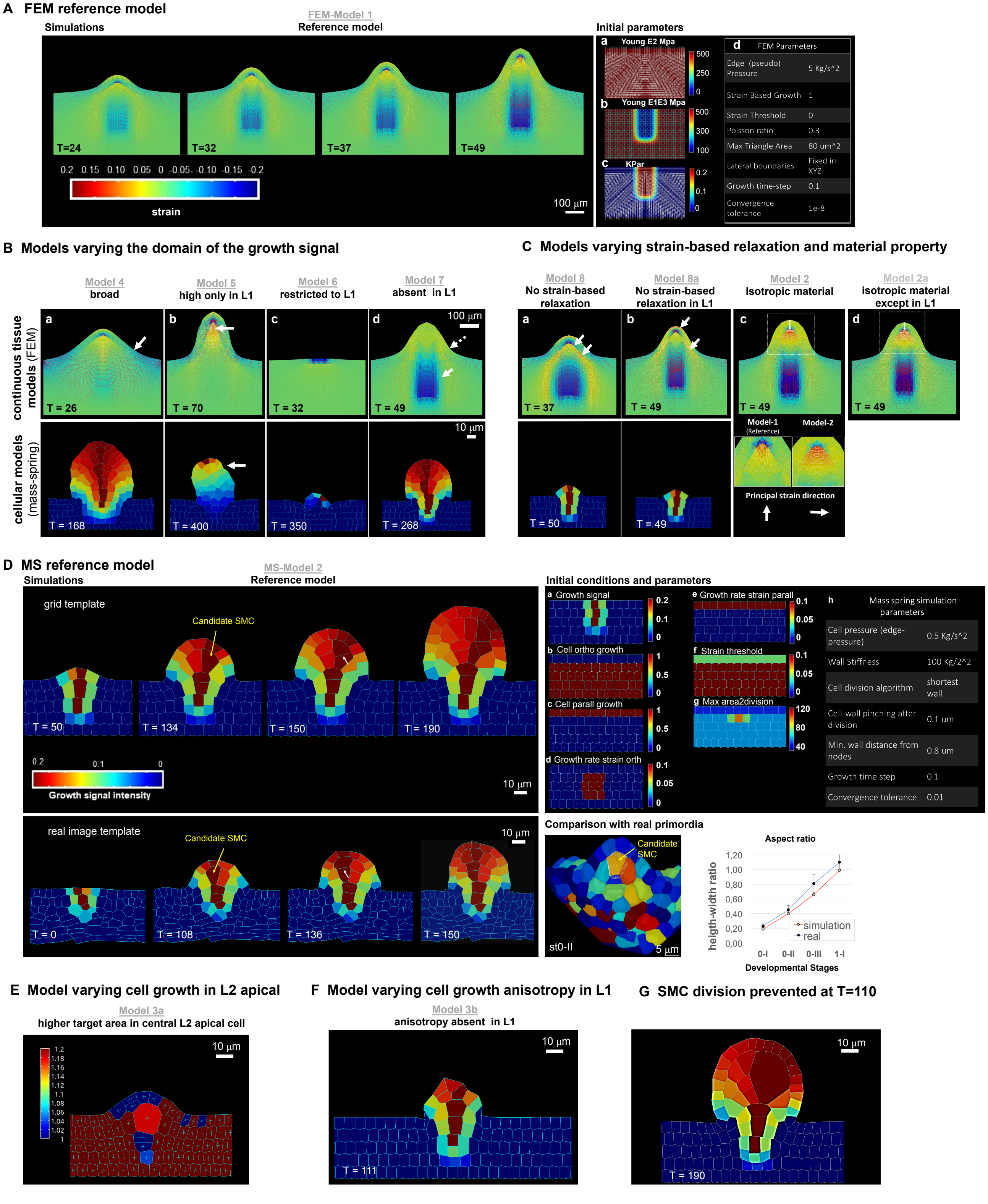


**Supplement Figure 3_ Related to Figure 3. FEM- and MS-based simulations, reference models, and additional variations.**

**Supplement Figure 3_ Related to Figure 3. FEM- and MS-based simulations, reference models, and additional variations.**

(A) *Time series of the FEM reference simulation: FEM-Model 1 and its preparation.* **Left***:* reference simulation progression; the lines represent the in-plane principal strains: white for extensive, red for compressive. The heatmap represents the trace of the Green-Lagrange strain tensor. **Right**: preparation of the reference template (a-c) and initial parameters (d). (a-c) the starting template is a flat tissue representation distinguishing L1 and inner layers on which anisotropic material (a, Young's modulus E2), material stiffness (b, Young's modulus E1E3), and the growth signal (c, Kpar) domain are set. The white lines indicate the direction of fiber reinforcement for the anisotropic material (a), the intensity of the stiffness in the isotropic component of the material (b), the direction of signal-based growth, which is orthogonal to the fiber reinforcement direction (c). In addition, all lateral boundaries of the simulation template (except the top one) were fixed in all degrees of freedom. The table (d) integrates the visual information with global parameters. See additional explanations in the Supplemental File 1.

(B) *Variation of the signal-based growth distribution with respect to the FEM- and MS-based reference models.* For all model variations, the result is shown for the continuous FEM-based simulations (top) and the MS-based simulations (bottom) at the simulation time point indicated (T). (**a**) Model 4: the signal is allowed to diffuse broadly. In both FEM- and MS-based simulations, the primordium is broader and the flanks are not orthogonal to the surface (arrow) as in the reference models. (**b**) Models 5: L1-driven growth hypothesis (1). Here the growth-specifying signal is set at a high and fixed (i.e., non-diffusing) concentration only in the L1. In both simulations, these conditions allow the formation of a protrusion but not a realistic primordium. For both simulations, the growth time went beyond the reference time to reach the same final height as in the reference models and to compensate for the prescribed reduced growth intensity. Specific observations are made for the different simulations: FEM-Model 5 shows high compression forces below the L1 (arrow: compressive areas are marked by the blue color and red lines). One interpretation is that these compressions may arise because the L1 "collapses" towards the center in the absence of inner tissue growth. Note the narrower apical, subepidermal domain than in the reference model. Furthermore, in the periclinal direction the L1 experiences significant compressions (indicated by the red lines). MS-Model 5: the growth is not balanced and generated an unstable apex (arrow, winding in different directions during growth, see movie in Supplemental File 3). (**c**) Models 6: L1-driven growth hypothesis (2). Here the growth signal is restricted solely to the L1. In this case, the difference is striking for both FEM- and MS-based simulations: the models are not able to create a significant buckle and, even if strain-based growth is permitted, the inner tissue is not pulled upwards. The growing part of the L1 ends being compressed by the non-growing counterpart (as shown by the compressive strain marked in blue in the image for the FEM-based simulation), causing the simulation to numerically fail earlier than the reference (FEM) or in any case not forming a primordium (MS). (**d**) Models 7: exclusive inner-tissue growth hypothesis. Here, the growth signal is absent in the L1. The L1 performs exclusively passive strain-based relaxation. In this model, we test whether it is plausible that growth can be entirely initiated by the inner tissues, provided the L1 has a higher ability to relax (compared to the remaining tissue) due to induced strain. In such a scenario, the inner layer, which is prescribed to grow as in the reference model, pushes the L1, causing accumulation of tensile strains in the periclinal direction that get released due to passive strain-based relaxation. Both FEM- and MS-based simulations are able to produce a digit-shaped protrusion, even if, in the FEM-based simulation, it is shorter with less straight flanks (i.e., smoother transition from the placenta to the flanks; dotted arrow). This can be explained by the absence of compressive strains at the base of the ovule (plain arrow) in comparison with the reference model. The MS-based model displays a primordium with increased width at the apex. A possible explanation for this may be the absence of growth constraints generated by the L1. In both cases, the passive strain-based relaxation coefficient of the L1 was higher than in the reference model. Whether this is biologically relevant remains to be addressed experimentally.

(C) *Models varying passive strain-based relaxation and material properties.* Results are shown for the continuous FEM-based simulations (top) and the MS-based simulations (bottom) at the simulation time point indicated (T). (**a**) Models 8: absence of passive strain-based relaxation in all the tissue. In both simulations, this results in a poorly developed primordium (and numerically the simulation fail earlier than the reference time), likely due to the accumulation of high strains (and stresses), particularly visible in the FEM-based model (high accumulation of compressive and tensile strains near each other in the inner tissue as well as in the L1, arrows). (**b**) Model 8a: variation of the previous model with absence of passive strain-based relaxation only in the L1. In both simulations, the primordium is shorter than in the reference case, although less severe in the FEM-based simulation. Specifically, in the L1 there is a high accumulation of tensile stresses on the outer walls in the apical portion and in the inner walls at the base (arrows). (**c**) FEM-Model 2: absence of anisotropic material properties in all tissues. Although the simulation produced a well-shaped primordium, the L1 is slightly thicker (double sided arrow, 1.2-fold compared to reference Model 1) and there is a larger L2,L3 apical domain characterized by a principal strain oriented 90° to that in the reference model (insets). These morphological differences with the reference model can be explained by the fact that passive strain-based relaxation in an isotropic material has an equal effect in the periclinal and the anticlinal direction, producing a thicker L1 and a rounder dome. In (**d**) FEM-based Model 2b: variation of previous Model 2. Here, only the L1 has anisotropic material properties. This restores the L1 thickness control but the abnormally high intensity and modified direction of compressive strains are similar to that in Model 2.

(D) *Mass spring reference model and initial template preparation, comparison with data from real ovules.* **Top left:** MS-Model 2 simulation at four different time points. The heatmap indicates the intensity of the growth-specifying signal. **Bottom left**: reference simulation performed on a template obtained from a real ovule placenta (see also Supplemental File 1). **Top right**: Initial parameters at simulation start. Field values as indicated (**a-g**) used to set up the reference configuration shown on the placenta at simulation time 0 and MS global parameters **(h)**. More details can be found in the Supplemental File 1. **Bottom right**: comparison between simulation and real primordia. *Image*: 3D, segmented ovule primordia stage 0-II from the wild-type reference set (see main text), longitudinal view: the SMC (orange) is outlined in a white triangular mesh (color-code refers to cell volume). Its morphology is to be compared to SMC candidates produced in the reference Model MS-Model 2 (e.g., T=134 for grid-based simulation and T=108 for real-image template-based simulation): the cell is elongated, centrally located in the apical dome and presents a broader upper periclinal wall surface compared to the bottom one. *Plot*: comparison of height-width ratio along simulation progression in MS-Model 2 and developmental progression in real (from segmented images of the wild-type reference set, see main text). For the simulation, two equi-distanced time-points between the $1^{st}$ (T = 0) and the last simulation time step (T = 190) have been used to compute the values for intermediate phases.

(E) *Variation of MS-Model 3- higher target area in central, L2 apical cell (MS-Model 3a).* This model is based on the same hypotheses as in MS-Model 3. The only difference being that the central L2 cell, which in the other models has a target area of 100 $um^2$, has now a bigger target area prior to division (200 $um^2$). The aim was to test whether longer cell growth influences anisotropy. The heatmap code represents the anisotropy index as defined in Supplemental file 1 and is scaled up to 1.2, hence with a narrower range than in Figure 3, to allow to appreciate the mild anisotropy level in the candidate SMC (index 1.18, a fully isotropic cell has index equal to 1, the fact that surrounding cells reach the maximum anisotropy index on this limited scale is not informative here). Furthermore, one can qualitatively appreciate that the cell shape is rather trapezoidal. This shows that if the SMC candidate grows longer than neighbouring cells, tissue geometrical constraints can confer cell anisotropy and a trapezoidal shape even in the context of isotropic specified growth, to reach a certain amount of anisotropy.

(F) *Model lacking cell growth anisotropy specifically in the L1 layer (MS-Model 3b).* Strong alterations of ovule dome growth and shape were observed when isotropic cell growth was assigned exclusively to the L1 layer (complementary to growth anisotropy hypothesis in MS-Model 3). The ovule overall shape alterations are likely due to the fact that the same amount of signal based growth has to be redistributed in the anticlinal and periclinal direction, limiting, though, the amount by which cells are prescribed to elongate. This, in turn, has a constraining effect on the inner layers. By comparing MS-Model 7 (Supplement Figure 3B), where the L1 performs exclusively strain-based growth and MS-Model 3b, where selectively the L1 grows isotropically, it is possible to infer that the anticlinal component of growth in the L1 actively constraints the inner layer expansion. This constriction, in turn, causes a lack of tension build up at the interface between the L1 and L2 which limits then further periclinal strain-based growth in the L1, producing an ovule with a shorter dome and narrower apical domain.

(G) *Model intervening on SMC division*. In the reference MS-Model 2, there is no explicit rule preventing division of the SMC candidate. To analyze the consequence of SMC growth on primordium growth in our models, we prevented the SMC candidate to divide at simulation time T = 110. Under the reference conditions where the turgor pressure is the same among all subepidermal cells, this generates a gigantic cell exerting a considerable tensile stretch on the surrounding cells, deforming the whole domain. This suggests that, in real ovules, SMC growth might be compensated by a reduction in turgor pressure, a hypothesis that needs to be tested. Note that the two direct adjacent cells have an elongated shape reminiscent of companion cells.

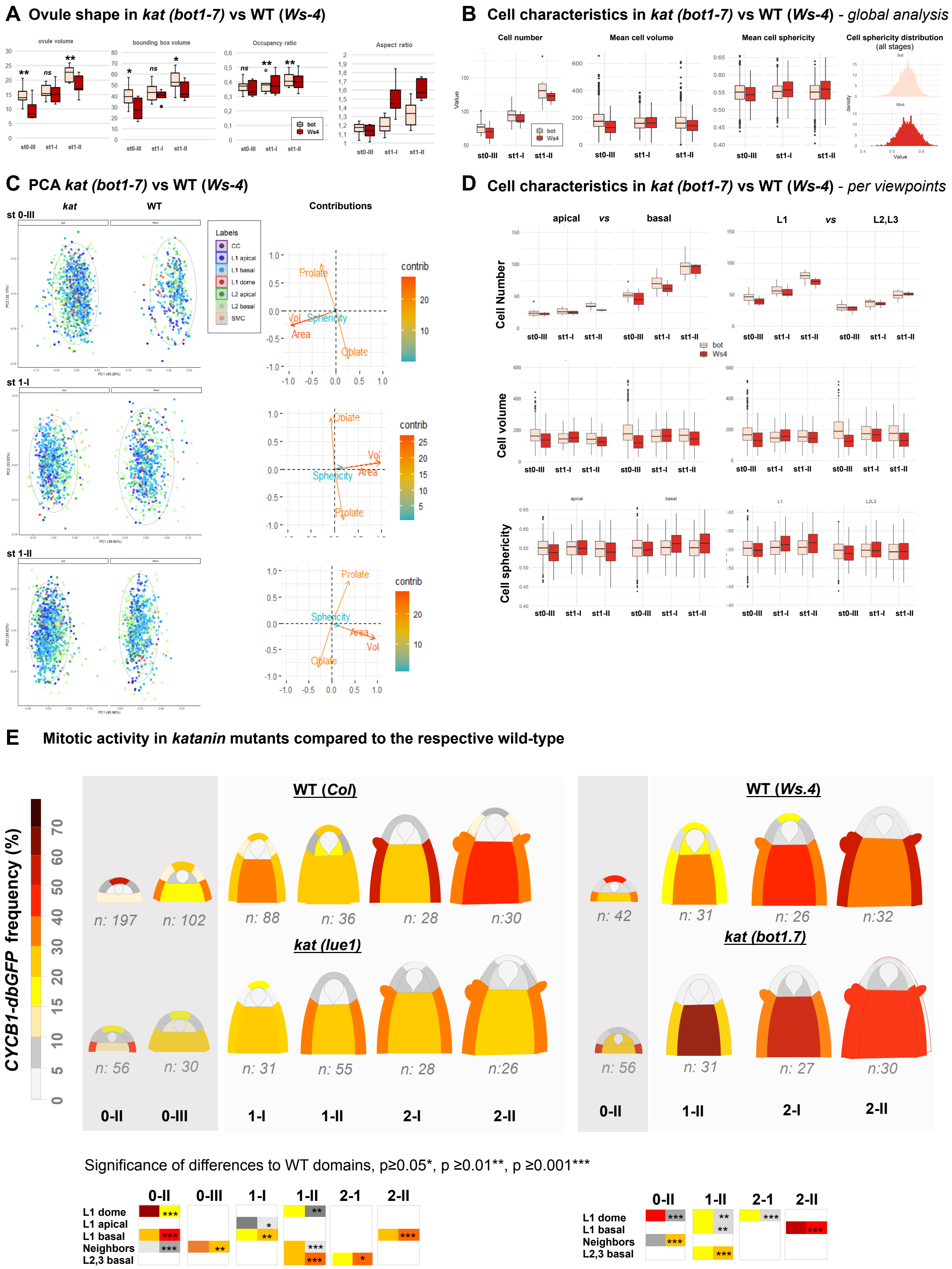


**Figure supplement 4_Related to Figure 4. *katanin* ovule growth defects.**

**Figure supplement 4_Related to Figure 4. *katanin* ovule growth defects.**

(A) *Ovule size, aspect ratio, bounding box, and bounding box occupancy.* Ovule volume was calculated as the sum of cell volumes for individual ovules (OvuleViz data). For each ovule, the volume of their fitting bounding box (represented Figure 4D) was calculated using box side lengths given in Imaris (Bitplane, AG) on ovule primordia treated as Surface objects (see Materials and Methods). Bounding box occupancy was calculated as the ratio of the ovule volume relative to its bounding box volume. The aspect ratio was calculated as the height (C length of the bounding box) divided by the mean of the width and depth lengths (A and B lengths of the bounding box). Man Whitney U test was used to assess differences between genotypes. Related to Figure 4B-D.

(B) Comparison of cell number, cell volume, and sphericity in wild-type (WT) (Ws-4) and *katanin* (*kat*) ovule primordia (*bot1-7 allele*). Man Whitney U test was used to assess the differences between genotypes.

(C) Global PCA analyses of cell characteristics in *kat* versus WT ovules (*Ws-4*). PCA was performed on a dataset containing Sphericity, Ellipticity oblate, Ellipticity prolate, Area, and Volume measurements of WT and *kat* cells from ovules at the indicated stages (0-III, 1-I, 1-II). Cell types are labelled according to colour code indicated. 95% confidence ellipses are indicated (dashed lines). Only the first and the second Principal Components (PCs) are shown. The loading plots indicate the contributions of the variables to each component. Related to Figure 4E.

(D) Comparison of mean cell number and volume in apical versus basal domains, and in L1 versus L2,L3 layers between WT (Ws-4) and *kat* (*bot1-7*) ovule primordia. Man Whitney U test was used to assess differences between genotypes. Related to Figure 4E.

(D) Comparison of cell sphericity as in (C). Related to Figure 4E.

(E) Mitotic activity in distinct domains of the ovule primordium of WT (*Col*, *Ws-4*) and *kat* (*lue1, bot1.7*) mutant plants. WT *Col* data are presented also in Figure 2 and shown here for better comparison only. The frequency of mitoses was scored using the CYCB1.1db-GFP reporter. Dark gray regions mark Phase I developmental stages. Two-tailed Fischer's exact test was used to assess differences between domains in the genotypes. Significant P values (*p≤0.05, **p≤0.01, ***p≤0.001) are indicated for the different domains in coloured boxes below the heatmaps (first column: WT; second column: *kat*). Non-significant differences are not represented. See also Source Data 2 and Source Data 4. Related to Figure 4F.

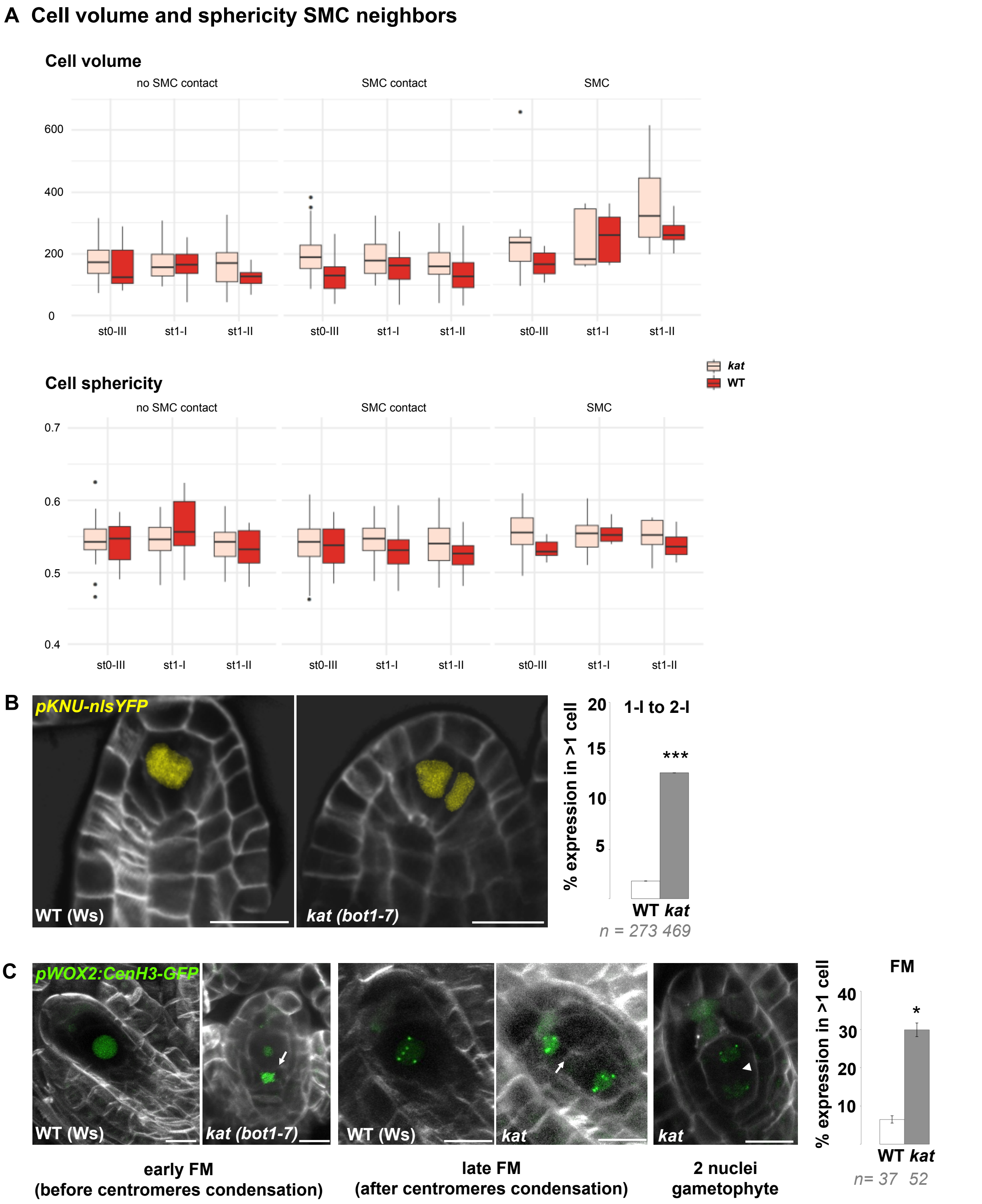


**Figure supplement 5_Related to Figure 5. Altered ovule primordium geometry in *katanin* mutants is associated with multiple SMCs.**

**Figure supplement 5_Related to Figure 5. Altered ovule primordium geometry in *katanin* mutants is associated with multiple SMCs.**

(A) Comparison of volume and sphericity of L2,L3 apical domain cells in wild-type (WT, *Ws-4*) and *katanin* (*kat*, *bot1.7* allele). Mann Whitney U test was used to assess the differences between genotypes for the mean cell volume and sphericity visualized by boxplots. Related to Figure 5B.

(B) Representative images and quantification of WT (*Ws-4*) and *kat* (*bot1-7*) ovule primordia expressing *pKNU::nls-YFP* (yellow signal), specifically expressed in the SMC. *kat* ovules show ectopic subepidermal cells with YFP signal, confirming SMC fate acquisition. Related to Figure 5E.

(C) Representative images (ca 10um maximum intensity projections of image stacks) and quantification of WT (*Ws-4*) and *kat* (*bot1-7*) ovule primordia expressing *pWOX2:CenH3-GFP* (green signal), labeling centromeres from the functional megaspore (FM) stage onward. Cells ectopically expressing the marker are detected in *kat* at both early (before centromeres condensation – left panel) and late (after centromeres condensation – middle panel) stages of FM formation. Ectopic spores were scored based on the presence of a cell wall (white arrows) separating the spores, by contrast to two-nuclei gametophyte stage (right image, shown as an example) where the two nuclei are separated by a vacuole (arrow head). Note that ectopic spores are aligned with the normal, most basal (chalazal) FM, suggesting that they belong to the same tetrad. They also display a similar number of centromeres (5) visible on the projections. Quantification was performed on pooled early and late FM stages. Related to Figure 5D-F.

White signal: Renaissance SR2200 cell wall label. Scale bar: 10µm. n: number of ovules scored. Error bar: standard error of the mean. Two-tailed Fischer's exact test was used to assess differences between domains in the distinct genotypes. P values: *p≤0.05, **p≤0.01, ***p≤0.001. See also Source Data 5.

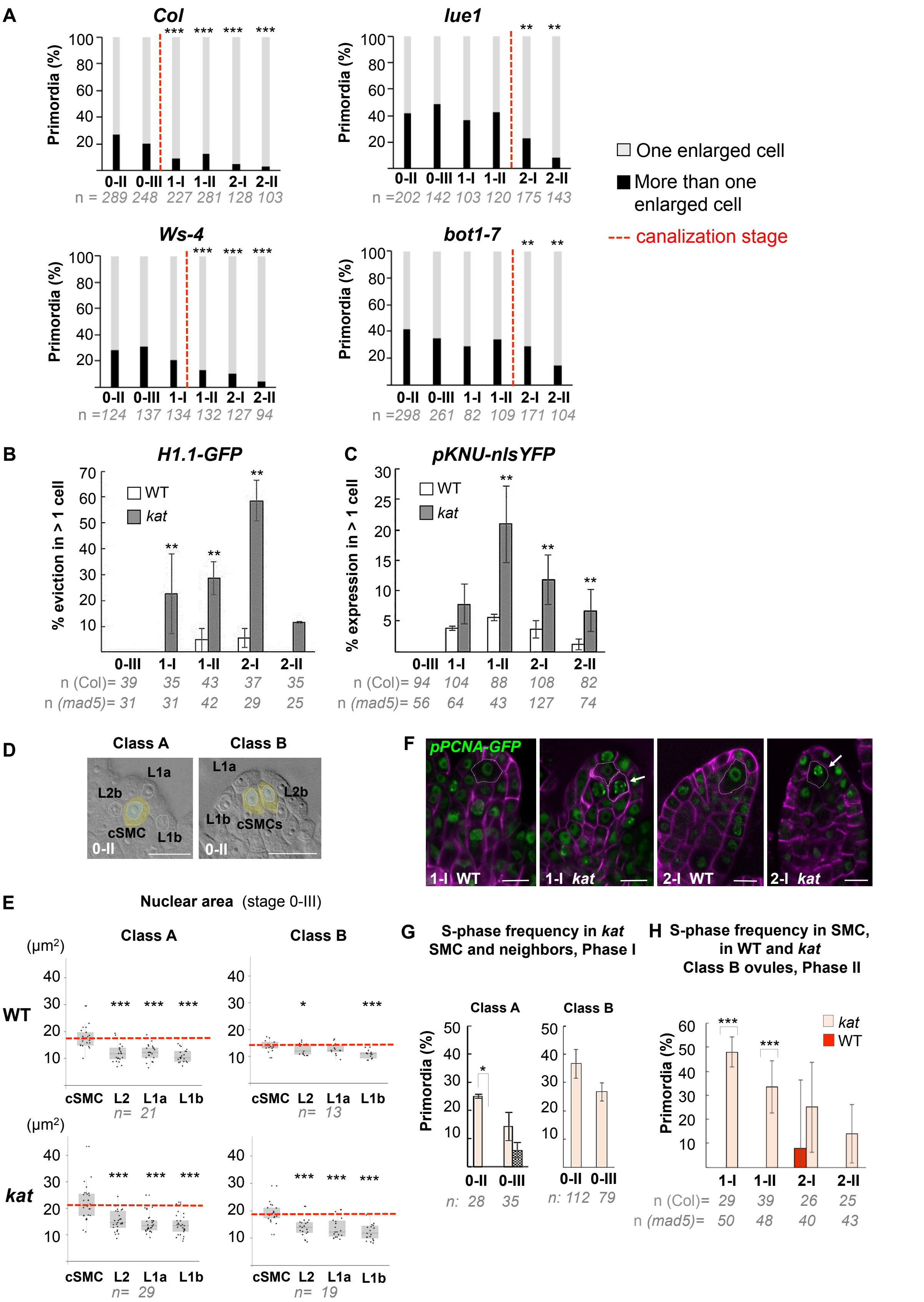


**Supplement Figure 6_Related to Figure 6. SMC singleness is progressively resolved during primordium growth.**

**Supplement Figure 6_Related to Figure 6. SMC singleness is progressively resolved during primordium growth.**

(A) Canalization of SMC fate in different *katanin* (*kat*) alleles (*lue1, bot1-7*) and respective wild-type (WT) backgrounds (*Col, Ws-4*). Percentages of class A and B primordia based on cleared tissue preparations as shown Figure 6A. The proportion of primordia with several SMC candidates at stage 0-II gradually decreases with developmental progression while the proportion of ovules with a single SMC increases, suggesting canalization. A significant delay in canalization (dashed red line) in *kat* mutants was confirmed by a two-tailed Fischer's exact test (see also Source Data 2). Related to Figure 6B-C.

(B-C) Ectopic expression of SMC fate in *kat* primordia (*mad5* allele) compared to WT (Col). Percentage of ovule primordia showing H1.1-GFP eviction (B) and KNU-nlsYFP expression (C) in more than one SMC candidate at all developmental stages. Two-tailed Fischer's exact test was used to assess differences between genotypes. Related to Figure 6B-C and Figure 6D-E.

(D-E) Nuclear area as a marker of SMC candidates. Illustration of the cells selected for nuclear area measurements in representative class A and B ovules (D). Nuclei areas were measured in two L1 cells types (L1a, apical position and L1b, closer to the placenta), in cSMC: candidate SMC; L2b: a neighbouring L2 cell. Dashed circles mark the nuclei of these cells used in the measurements. Related to Figure 6D. Quantification of nuclear area as shown in (D) for class A and B in WT (*Col*) and *kat* (*mad5* allele) ovule primordia at stage 0-III (E). Box plot representations include jittered points to visualize data variability. Red lines represent the median of cSMC nuclear area for comparison with the other cells. Wilcoxon signed rank test was used to assess difference between SMC and the other cell types. Related to Figure 6D.

(F-H) Frequency of SMCs showing an S-phase pattern in *kat* and Class B primordia. F: Representative images; green, PCNA-GFP, magenta: Renaissance SR2200 cell wall label. Scale bar 10µm. Related to Figure 6F, 6H. G: Quantification of S-phase occurrence in SMCs (salmon bars) and neighbors cells (dashed bars) of class A and class B *katanin* (*mad5*) ovule primordia in Phase I (stages 0-II and 0-III). H: Quantification of S-phase occurrence within the L2 apical domain of class B WT (*Col*) and *kat* (*mad5* allele) ovule primordia in Phase II (stages 1-I to 2-II). The plotted frequencies refer to the number of primordia showing a speckled PCNA-GFP pattern in SMC candidates. Two-tailed Fischer's exact test was used to assess differences between genotypes. Related to Figure 6H.

All graphs: n= number of ovules scored. Error bar: standard error of the mean. P values: *p≤0.05, **p≤0.01, ***p≤0.001. See also Source Data 6.
