## Supplemental File 1 for "Organ geometry channels cell fate in the Arabidopsis ovule primordium"

### Supplementary Methods

#### 1.1 Mass spring-based ovule growth simulations at cellular resolution

##### 1.1.1 Template preparation and initialization - Regular grid of cells

Following observation of microscopy images of longitudinal sections of carpels showing the placenta, a regular grid of cells has been created with MorphoGraphX(de Reuille et al., 2015) (www.MorphoGraphX.org) (Cell Maker plugin). The grid is composed of 28 columns by 5 rows of square cells of side 5.44 $\mu$m, the cell rows are staggered with respect to each other by 1.36 $\mu$m in the longitudinal direction.

The mass spring mesh is constituted by interconnected segments outlining the cells boundaries and representing the cell walls. The segments have a varying length of 1.36 $\mu$m and 2.72 $\mu$m, the top margin of the grid of cells has a segment length of 5.44 $\mu$m (this is to increase mass spring stability w.r.t. compressive forces generated during ovule growth). A dual mesh connecting the cell centers is associated to the mass-spring mesh.

The template initialization as well as the proper growth simulations are run with an in-house built plugin for MorphoMechanX (www.MorphoMechanX.org) available at a git repository:
github.com/GabriellaMosca/MassSpring_2DovuleGrowthModel.

Once the template has been generated, it needs to be initialized with the parameters illustrated in Figure S3B. These properties are assigned at the cell level (no gradient for these properties within a cell), so on the dual graph.

The growth signal is the result of a discrete, cell-based diffusive process. The user sets some cells with fixed high concentration and some other cells with fixed lower concentration of the diffusive substance, while the other cells get their concentration as a result of the diffusive process. More specifically, for the simulations of the growth signal in this work, some cells were assigned a fixed concentration of 0.2 [1] (the vertical column made by the three cells starting from L1 with the highest growth signal concentration in the template, see Figure S3B) and some cells were assigned a fixed concentration of 0 (a well-shaped arrangement of cells around the ones with a concentration different from zero, as can be seen in Figure S3B). In the model, this is done by running the process **Model/Cell Ovule Growth/10 Growth Signal/a Set Cell Type/Assign**, after setting the variables to the desired values.

Prior to pressurization, Dirichlet boundary conditions in the x-direction have been assigned to the lateral cell walls of the template (this means that those walls are not allowed to displace in the x-direction during pressurization). This condition has been assigned to simulate the presence of the surrounding lateral tissue. Dirichlet boundary conditions can be assigned by running the process **Model/Cell Ovule Growth/20 Mass Spring Mechanics/a Set Dirichlet** after having replaced the 3D null vector in the field “Dirichlet” with a value different from zero for each coordinate to be fixed. This will affect all the selected nodes of the mass spring mesh.

The process **Model/Cell Ovule Growth/20 Mass Spring Mechanics/b Set Cell Type Mechanics and Growth/Assign** has been used to assign (i) the different growth directionality (see Figure S3B); (ii) differ-

[1] The units have been a-dimensionalized.

ent strain threshold for strain-based growth for the L1 cells and the inner ones with a growth signal different from 0 (the remaining cells deform elastically, but are not allowed to perform any growth) (see Figure S3B); (iii) different target area for cell division (53 $\mu$m$^2$ for L1 cells, 100 $\mu$m$^2$ for the SMC precursor, 80 $\mu$m$^2$ for the L2 cells neighbors of the pSMC, while all the other cells got the global target area for division of 63 $\mu$m$^2$, see Figure S3B ). To correctly use the process, it is necessary to label L1 cells as "L1" and the other cells with special assigned properties as "Special Cell".

Furthermore, the L1 is prescribed to perform only periclinal growth, by running the process **Model/Cell Ovule Growth/20 Mass Spring Mechanics/d Only Periclinal Growth in L1** with the field "Value" set to "True". When this option is assigned, the L1 cell edges corresponding to anticlinal walls are prevented from growing (this process will act on the cells previously labeled as "L1").

It is now necessary to set the polarization field used to define growth directionality: this is obtained as the gradient of a diffusion process. The user needs to select cells with fixed high and low concentration (the specific value assigned here is arbitrary and functional only to be able to set a gradient field which will be normalized). In our simulation the cells in the left side of the template have been set with fixed high concentration, while the cells in the right side of the template have been set with fixed low concentration (whether high/low concentration is fixed on the left or right corner is irrelevant): this produced the gradient visible in Figure S3B. To set the cells with fix concentration for the polarization process, the user needs to run the process **Model/Cell Ovule Growth/30 Growth Process/a Set Fixed Concentr Diffusion Polarizer/Assign**, after having assigned in the dialog box the desired values.

The remaining field values are global and defined in the Table of Figure S3B.

After this preparation, the template can be inflated to the prescribed pressure (in our model 0.5 Kg/s$^2$). It is important to stress out that in these simulations, the pressure adopted does not have the units of a physical pressure, but those of a force acting on a 1D segment (instead of on a 2D surface). Connected to this point, we stress out that these 2D simulations have a purely qualitative purpose and no quantitative comparison between the parameters used and the bio-physical quantities should be attempted.

After pressurization of the template, new Dirichlet boundary conditions are assigned: the nodes outlining the boundary of the template (except for the L1 top nodes) are blocked in all degrees of freedom (XYZ). This mesh is saved as starting point for the growth and cell division simulation with mass springs.

##### 1.1.2 Confocal microscopy-based template preparation

A confocal stack from an almost flat placenta (dataset 2404_275a) has been used. The raw stack has been loaded in MorphoGraphX and by means of the clipping planes, a longitudinal central section of it has been selected. Then, the same procedure described in the MorphoGraphX manual (available here: www.mpipz.mpg.de/4085950/MGXUserManual.pdf) for generating $2\frac{1}{2}$D mesh has been followed, with the only difference that the surface mesh is generated with the CellMaker plugin as a rectangle with side length 0.5 $\mu$m. The signal projection depth has been calibrated so that only the walls of the midsection get projected (the longitudinal mid section has been as well selected accordingly to not display spurious walls from cells at different depth) and set to the value of 1 $\mu$. The final mesh gets converted into a 2D Cell Mesh by the process: **Process/Mesh/Cell Mesh/Convert to 2D Cell Tissue** with the field "Max Wall Length" set to 10. After cleaning spurious boundary cells, and some manual curation to remove edges which are not junctions on the top wall of L1 cells (this to reduce the artifact of edge collapsing due to the absence of bending resistance during compressive forces), one obtains the final template to be used in the simulation within MorphoMechanX. The template preparation is then identical to what explained in the previous section and the distribution of cells with fixed high concentration is shown in Figure S3F.

##### 1.1.3 Simulation cycle

The growth and division simulation starts from the pressurized template at mechanical equilibrium (see following). A simulation cycle consists of the following processes:

- growth signal diffusion calculation;
- polarization field calculation;
- mass spring mechanical equilibrium calculation;
- growth step;
- cell division.

The simulation cycle is repeated as many times as indicated by the simulation time T (which is not in physical units) mentioned in the main text, in Figure 3 and Figure S3). We explain now in detail how each process is computed.

**Growth signal diffusion calculation** The growth signal is the result of a diffusive process acting on the cell level (on the dual graph). In this formulation, no sources or sinks are present: only some cells with a fixed concentration are specified at the beginning of the simulation. The discretized diffusion equation, following what done in Bayer et al. (2009), models the concentration variation inside a cell according to this formula (expressed in adimensionalized units through rescaling of the diffusion constant D):

$$\frac{d\rho_i}{d\tau} = \frac{D}{A_i} \sum_{j \in N_i} l_{ij} \left(\rho_j - \rho_i\right) \tag{1.1}$$

where $\rho_i$ indicates the signal concentration in the cell $i$, $\tau$, $D$ the diffusion coefficient, $A_i$ the area of cell $i$, $N_i$ is the set or cells connected to the cell $i$, while $l_{ij}$ is the length of the interface between the two cells. The rationale for this heuristic formula, formally identical to the discrete Laplace-Beltrami operator if the weight factors are made to coincide with $l_{ij}$ (see Reuter et al. (2009)), is that the variation of concentration in the cell $i$ depends on the difference of concentration between this cell and the neighboring ones rescaled by the cell surface (the bigger the cell, the less the concentration variation) and the length of the interface with each neighboring cell. Since we are looking for the steady state solution in the absence of sources and sinks, the rescaled diffusion coefficient $D$ does not affect the final solution and so it is set to 1 in the following calculations. To find the steady-state concentration, an adaptive forward Euler scheme was adopted Teukolsky et al. (1992) (the problem is easily solvable by a direct method, but this diffusion process has been designed to treat more general, non linear problems as well). So an iteration of the forward Euler scheme, to calculate the variation in concentration for the cell $i$ at the discrete simulation time step $n+1$, is as follows:

$$\rho_i(\tau_{n+1}) - \rho_i(\tau_n) = \begin{cases} \frac{1}{A_i} \sum_{j \in N_i} l_{ij} \left(\rho_j(\tau_n) - \rho_i(\tau_n)\right) * \delta\tau, & \text{if } \rho_i \text{ is not fixed} \\ 0, & \text{otherwise} \end{cases} \tag{1.2}$$

The known fixed concentration are initialized with their assigned value. The time increment $\delta\tau$ is chosen based on an adaptive scheme which evaluates the magnitude of the concentration variation at the previous step to determine the new time increment (constrained between a preset max and min allowed time increment). The simulation is considered converged when the update on the concentration $\rho_i(\tau_{n+1}) - \rho_i(\tau_n)$ is smaller than a prescribed tolerance, more precisely when the mean between the averaging of the concentration update over all vertexes and the maximal of the concentration is below a threshold (concentrations taken as absolute values).

**Polarization field calculation** The polarization field is computed as the gradient of a diffusive process which is regulated by cells with a fixed concentration prescribed. The evolution process itself is computed as in the previous paragraph. Once the equilibrium concentration has been obtained, its 2D gradient (on the cell grid) is computed and then normalized. This gradient indicates the direction of the ”polarizer” $\mathbf{n}_{Pol}$, which is used to control the direction of anisotropic growth. The advantage with respect to using fixed coordinates is that the polarization field is updated while the mesh changes, creating a system of local coordinates that follows the organ natural curvature while it grows (Kennaway et al., 2011).

**Mass spring mechanical equilibrium calculation** The mass-spring system is constituted by the network of linear springs which describe the cell walls in the template. The mechanical equilibrium is obtained when the internal reaction forces generated by the mass springs counterbalance the externally applied loads at each point of the mass spring network (MSN):

$$\mathbf{F}_{I_v} + \mathbf{F}_{E_v} = \mathbf{F}_{TOT_v} = 0 \quad \forall v \in MSN \tag{1.3}$$

where $\mathbf{F}_{I_v}$ indicates the internal reaction force at the node $v$, $\mathbf{F}_{E_v}$ the sum of the external forces applied to the node $v$ and $v$ belongs to the mass-spring-network ($MSN$). More specifically, the internal forces are given by the Hooke's law for mass-springs:

$$\mathbf{F}_{I_v} = k \sum_{\mathbf{u} \in N_v} (l_{u \to v} - \|\mathbf{p}_u - \mathbf{p}_v\|) \frac{\mathbf{p}_u - \mathbf{p}_v}{\|\mathbf{p}_u - \mathbf{p}_v\|} \tag{1.4}$$

$k$ is the spring constant (Kg/s$^2$), $N_v$ is the set of points which are connected to $v$, $l_{u \to v}$ is the rest length of spring connecting the point $u$ to the point $v$, while $\mathbf{p}_u$ and $\mathbf{p}_v$ are the respective positions. Note that $\frac{\mathbf{p}_u - \mathbf{p}_v}{\|\mathbf{p}_u - \mathbf{p}_v\|}$ is the unit normal vector $\mathbf{n}_{u \to v}$ in the direction connecting the point $u$ to the point $v$. In our case forces are measured in $\mu$N and lengths in $\mu$m, but, being this a 2D representation of a 3D problem by means of 1D structures such as the mass springs, no quantitative comparison with bio-mechanical properties of the cell tissue should be attempted.

Regarding the externally applied forces, turgor pressure is considered. In our 2D mass-spring model this acts on the 1D line segment connecting two vertices in the mass spring system:

$$\mathbf{F}_{E_v} = \mathbf{F}_{P_v} = \frac{1}{2} \sum_{u \in N_v} \sum_{C \,|\, u \to v \in \mathcal{B}_C} P_C(l_{u \to v}) \mathbf{n}^{\perp}_{u \to v} \tag{1.5}$$

here $\mathbf{F}_{P_v}$ stands for the force due to turgor pressure on the node $v$, $\mathbf{n}^{\perp}_{u \to v}$ indicates the unit vector orthogonal to $\mathbf{n}_{u \to v}$ and pointing outwards with respect to the cell centroid. The summation is indented over all the vertexes $u$ connected to $v$ and over all the cells $C$ whose the edge $u \to v$ is part of the boundary ($\mathcal{B}_C$) and $P_C$ is the pressure afferent to cell $C$. To compute the forces equilibrium a semi-implicit Euler method has been adopted (Teukolsky et al., 1992):

$$\mathbf{p}(n+1) - \mathbf{p}(n) = h \left[\mathbf{1} - h\mathbf{J}(n)\right]^{-1} \mathbf{F}_{TOT}(n) \tag{1.6}$$

Eq. 1.6 is in a matrix form: $\mathbf{p}(n) = (\mathbf{p}_1(n), \cdots, \mathbf{p}_m(n))$, $\mathbf{F}_{TOT}(n) = (\mathbf{F}_{TOT1}(n), \cdots, \mathbf{F}_{TOTm}(n))$ and $\mathbf{J}(n)$ is the Jacobian (here computed numerically) of the force vector with respect to the nodal position vector $\mathbf{p}(n)$. $\mathbf{p}_i$ is the position of the $i$-th node in the $MSN$, while $n$ is the $n$-th iteration of the solver. The time stepping, $h$, is computed adaptively and based on the norm of the incremental displacement. The mechanical equilibrium is considered reached when the mean between the averaging of the norm of forces over all vertexes and the maximal force norm is below a threshold. The nodes with prescribed Dirichlet boundary condition will have the corresponding force equal to zero in the direction in which the condition is applied.

In case the option "Wall Stiffness based on morphogen" in the process **Model/Cell Ovule Growth/20 Mass Spring Mechanics/c Set Wall Stiffness/Assign** has been set to true, for the selected cells during the simulation runtime, stiffness will be assigned within the range specified by the user in a linear negative correlation with respect to the growth signal.

**Growth step** After mechanical equilibrium has been reached, an incremental growth step, acting on the rest lengths of the mass springs ($l_{u \to v}$) is computed. Different types of growth can be specified and added together. In the model there are signal based growth and strain based growth, both can be explicitly required to occur along or orthogonal w.r.t. the polarization field direction $\mathbf{n}_{Pol}$. The incremental growth operator acting on the rest length of a mass spring segment connecting $u$ to $v$ is given as follows:

$$l_{u \to u}(T+1) = \delta\tau (G^S_{u \to v} + G^{\epsilon}_{u \to v} + G^{\epsilon_{Pol}}_{u \to v}) l_{u \to v}(T). \tag{1.7}$$

Here $G^S_{u\to v}$ is the signal based growth contribution, $G^{\epsilon}_{u\to v}$ is the strain based growth contribution, $G^{\epsilon Pol}_{u\to v}$ is the strain based growth contribution in the polarizer basis ($\mathbf{n}_{Pol}, \mathbf{n}^{bot}_{Pol}$), $T$ is the discrete simulation time (number of simulation loop iterations) and $\delta\tau$ is a coefficient interpretable as a global growth scaling factor. More specifically:

$$G^S_{u\to v} = \begin{cases} K^S_{Par} & \text{if } abs(\mathbf{n}_{u\to v}\mathbf{n}_{Pol}) \geq 0.6 \\ K^S_{Per} & \text{if } abs(\mathbf{n}_{u\to v}\mathbf{n}_{Pol}) \leq 0.4 \\ 0.5(K^S_{Par} + K^S_{Per}) & \text{otherwise} \end{cases} \tag{1.8}$$

where $K^S_{Par}$ is the growth coefficient in the direction parallel to the polarization field, while $K^S_{Per}$ is the growth coefficient in the orthogonal direction. Since growth is performed by updating a rest length, the growth operator per-se can not affect directionality and introduce distortion. Therefore the notion of anisotropic growth is implemented by multiplying a rest length $l_{u\to v}(T)$ by $K^S_{Par}$ if its corresponding vector in the current configuration $\mathbf{n}_{u\to v}$ has a normalized projection along $\mathbf{n}_{Pol}$ bigger than 0.6. Vice-versa, if the projection is less than 0.4, the rest length will be multiplied by $K^S_{Per}$. For the remaining cases, an average of the two coefficients will be adopted. Regarding strain-based growth we have:

$$G^{\epsilon}_{u\to v} = \begin{cases} K^{\epsilon}\left(\epsilon(T) - \epsilon_{thres}\right) & \text{if } \left(\epsilon(T) - \epsilon_{thres}\right) > 0 \\ 0 & \text{otherwise} \end{cases} \tag{1.9}$$

where $\epsilon(T) = \left(\|\mathbf{p}_u(T) - \mathbf{p}_v(T)\| - l_{u\to v}(T)\right)/l_{u\to v}(T)$ is the linear strain computed at mechanical equilibrium at simulation loop time $T$ and $\epsilon_{thres}$ is the strain threshold. The rest length is not allowed to get bigger than the spring length in the current configuration.

The polarized strain based growth is prescribed as follows:

$$G^{\epsilon Pol}_{u\to v} = \begin{cases} K^{\epsilon}_{Par}\left(\epsilon(T) - \epsilon_{thres}\right) & \text{if } \left(\epsilon(T) - \epsilon_{thres}\right) > 0 \text{ and } (\mathbf{n}_{u\to v}\mathbf{n}_{Pol}) \geq 0.6 \\ K^{\epsilon}_{Per}\left(\epsilon(T) - \epsilon_{thres}\right) & \text{if } \left(\epsilon(T) - \epsilon_{thres}\right) > 0 \text{ and } (\mathbf{n}_{u\to v}\mathbf{n}_{Pol}) \leq 0.4 \\ 0.5(K^{\epsilon}_{Par} + K^{\epsilon}_{Per}) & \text{otherwise} \end{cases} \tag{1.10}$$

The growth coefficients used in these formulas are specified cell-based, while growth is performed edge-based: therefore the coefficients actually used are an average quantity over the cells related to the edge.

**Cell division** Cell division is performed according to the rule "shortest wall through the centroid" as described in Mosca et al. (2018). The target cell areas for division are assigned as described in Sec. 1.1.1. Cell wall pinching has been globally set to 0.1 (which corresponds to a proportionality factor connected to the new wall segment length or the distance between the newly inserted point and the cell centroid, depending on which is smaller), and is constrained to be overall smaller than 0.1 $\mu$m. The minimal distance from a pre-existing wall has been set to 0.8 $\mu$m. Cell wall sampling to search for the shortest wall has been set to 0.05 $\mu$m (the starting point for the sampling is aleatoric and in principle not guaranteed to be the same at different runs).

Upon cell division, cell properties (such as fixed growth signal concentration, fixed polarizer, cell type, target division area, special pressure, growth coefficients, etc.) are inherited by the daughter cells. For what concerns fixed high growth signal concentration, selective rules have been implemented to allow it to be propagated along a vertical line in the inner tissue. For what concerns cells in the inner layers, if a dividing wall is in the anticlinal direction, only one of the two daughter cells will inherit the fixed high signal concentration (the one with the smallest distance from the closest cell with a fixed high concentration). If the newly inserted wall is in the periclinal direction, if the mother cell (prior to division) is between two cells with a fixed high concentration, both daughter cells will inherit this property, otherwise only the one connected to the vertical stream of cells with a fixed high concentration. Regarding L1 cells, the daughter cells inherit both the fixed high concentration if the mother cell has two L1 neighbors with a fixed high concentration, otherwise only one cell will inherit this property with preference for the daughter cell with the shortest distance from the L2 cell with fixed high signal concentration. The propagation rules for fixed high concentration are not equivalent to a proper canalization model (which was not the aim of these simulations) and sometimes manual curation during the simulation time evolution has been required to ensure an uninterrupted vertical line of cells with fixed high concentration.

To assess the stability of the reference simulation (MS-Model 2), a robustness analysis has been performed by varying independently stiffness and growth signal intensity of 10% around the reference value. The results are qualitatively compatible with the reference model and available in the git repository: github.com/GabriellaMosca/MassSpring_2DovuleGrowthModel/tree/master/robustnessAnalysis.

#### 1.2 Cell anisotropy quantification

Similarly to what done in Louveaux et al. (2016), a shape matrix has been defined for each cell. This matrix is the covariance matrix of the coordinates of the points along the cell outline. Given a distribution of points in the 2D plane, their covariance matrix with respect to their coordinates in the Cartesian plane is defined as

$$\mathbf{Cov} = \frac{1}{N_c} \begin{pmatrix} \sum_{i=0}^{N_c}(x_i - x_0)^2 & \sum_{i=0}^{N_c}(x_i - x_0)(y_i - y_0) \\ \sum_{i=0}^{N_c}(x_i - x_0)(y_i - y_0) & \sum_{i=0}^{N_c}(y_i - y_0)^2 \end{pmatrix} \tag{1.11}$$

where $x_i$, $y_i$ are the i-th point coordinates, $N_c$ is the number of points and $x_0$, $y_0$ are the arithmetic average of all the points coordinates.

This matrix is symmetric and so diagonalizable, its eigenvectors indicate the directions of maximal and minimal points dispersion (and so the anisotropicity of the distribution). More precisely, the eigenvector connected to the biggest eigenvalue defines the direction around which the standard deviation of the projection of the points position along that axis is maximal, while the other eigenvector defines the direction around which this same quantity is minimal. At the same time, one can see the first eigenvector as the one minimizing the variance of the distance of the points distribution from the direction it indicates, while the second eigenvector as the one maximizing instead this quantity.

If the points are uniformly distributed around a shape outline, the eigenvectors can be used as a proxy for the maximal and minimal elongation directions of the shape, in the sense described above. If one considers a rectangle (where the concept of shape anisotropy can intuitively be assimilated to the aspect ratio of its sides), the eigevectors of this covariance matrix will be parallel to its sides and, if the sides ratio is greater than 1, also the corresponding eigenvalues will be greater than one. Furthermore the eigenvalues ratio depends on the sides lengths, but not on the rectangle area. If a rectangle has a aspect ratio greater than another (always computing longest size over shortest side), this relation will be valid also for the corresponding eigenvalues ratio.

If a rectangle is now a 2D cell which is growing and it is getting more elongated while not changing in the other direction, the eigenvalues ratio will increase. On the other hand if, while being anisotropic, it is still growing in an isotropic manner, its eigenvalues ratio will not change. So by looking at the eigenvalues ratio and their variation over time, it is possible to deduce the cell anisotropicity regardless of its surface and how it varies over time.

When a shape differs from that of a rectangle (or an ellipse), it is not possible to use the sides (main axes) aspect ratio as a measure of anisotropy, but the shape matrix provides an extension of this concept to other shapes. Clearly, the more the shape deviates form that of a rectangle, the more care should be used to interpret the results (especially with curved shapes, where a non-linear approach should be adopted).

In the measurements performed in this work, the cell perimeter has been enriched with intermediate points at an average distance prescribed by the user and the averaging has been weighted with the points distance to avoid artifacts due to non perfect homogenous points distribution. So the actual equation used to compute the covariace matrix is:

$$\mathbf{Cov} = \frac{1}{P_c} \begin{pmatrix} \sum_{i=0}^{N_c}(x_i - x_0)^2 l_i & \sum_{i=0}^{N_c}(x_i - x_0)(y_i - y_0) l_i \\ \sum_{i=0}^{N_c}(x_i - x_0)(y_i - y_0) l_i & \sum_{i=0}^{N_c}(y_i - y_0)^2 l_i \end{pmatrix} \tag{1.12}$$

Where $P_c$ stands for the cell perimeter and $l_i$ for the length of the perimeter segment containing the node $i$ in its middle. Furthermore, for the images shown in this work, it has been selected the option to measure cell anisotropy along the two orthogonal growth directions of the ovule. For this reason, by using $\mathbf{n}_{Pol}$ (the vector parallel to the polarization field direction, see Sec. 1.1.3) and $\mathbf{n}^{\perp}_{Pol}$, the orthogonal one, these two quantities are computed:

$$\alpha_{Pol} = \langle \mathbf{n}_{Pol}, \mathbf{Cov}\, \mathbf{n}_{Pol} \rangle \,, \qquad \alpha^{\perp}_{Pol} = \left\langle \mathbf{n}^{\perp}_{Pol}, \mathbf{Cov}\, \mathbf{n}^{\perp}_{Pol} \right\rangle \tag{1.13}$$

and their ratio is used as a proxy of cell anisotropy along the organ growth direction and the one orthogonal to it. More precisely, as in the figures the aim was to show the cell elongation along the ovule main growth direction, the anisotropy ratio displayed is given by this formula: $\alpha^{\perp}_{Pol}/\alpha_{Pol}$. This means that a displayed value of one indicates that the cell is fully isotropic, a displayed value bigger than one indicates that the cell is more elongated along the organ main elongation axis ($\alpha^{\perp}_{Pol}$), while a value between one and zero indicates that the cell is more elongated in the orthogonal direction to the main organ elongation axis. This calculation is performed in the model by running the process: **Model/Cell Ovule Growth/50 Shape Quantifier/01 Compute Skey Symmetric Tensor**, where the user can specify the average distance for the virtual points to be placed along the shape perimeter for the computation (one should refine this numer untill the result is not significantly affected). The eigenvectors displayed in Figure 3E are scaled by their pertinent eigenvalue divided over the sum of all three eigenvalues (normalization), all globally rescaled by a length factor.

#### 1.3 2D FEM based simulations for ovule growth - continuous limit

The finite element method (FEM) simulations are handled in MorphoMechanX by an in-house developed tool available at the git repository github.com/GabriellaMosca/FEM_2DOvuleGrowthModel. They run on a 2D mesh tessellated with triangles which constitute linear membrane elements with a mathematically prescribed thickness (not relevant in this type of simulation though).

##### 1.3.1 Template preparation

The initial template is build as rectangle tessellated by squares assembled in 28 rows and 56 columns. The squares have a side of length 13.0527 $\mu$m (the value assigned to the length is arbitrary as these simulations aim at a qualitative analysis in 2D of a 3D problem). The tessellation squares have been each triangulated with 4 triangles sharing a vertex in the square center (see Figure S3A).

This template represents a portion of the ovule placenta as seen through a longitudinal cut. The first three lines or squares from the top are identified with the L1 layer and are assigned specific properties different from the remaining tissue, as will be described in the following.

The description which follows is referred to the reference case template, FEM-model 1, as from Table 1 in the main text. The other models are obtained through minor modifications of the process described in the following.

As first thing, the growth signal intensity is assigned: this is obtained, similarly to what done with the mass springs, as the outcome of a diffusive process with some elements at a high and low fixed concentration. First of all, all the faces of the mesh need to be selected and the process **Model/CCF/Fem Diffusion Growth/04 Create Diffusion Element** has to be run, so that the element for growth signal diffusion is generated.

Then the nodes on the mesh with fixed signal concentration (high and low) need to be set. The nodes with high fixed growth signal concentration are forming a vertical line made of 16 rows by 7 columns starting from the top margin of the mesh, in a centered position (see Figure S3A for the signal distribution). By running the process **Model/CCF/Fem Diffusion Growth/03 Set Diffusion Dirchlet** with the field “Value” set to 0.2[2], the fixed high signal concentration is assigned to these vertices. Note that the “Dirichlet Attribute” and “Morphogen Attribute” (as well as the “Element Attribute”) names need to be assigned consistently for all the sub-processes contained in **Model/CCF/Fem Diffusion Growth**.

The vertices with a fixed low signal concentration are assigned with the same procedure. These vertices form the perimeter of a rectangle which encloses the vertical strip of cells with fixed high concentration. It is located at a distance from the mesh middle (in the horizontal coordinate) of 8 vertices, and from the mesh top (in the vertical coordinate) of 21 vertices (see SFigure A for the distribution of the signal). The fixed assigned concentration is 0.

After assigning the initial conditions, the diffusion process itself is run by **Model/CCF/Fem Diffusion Growth/00 Diffusion Solver**. The details on how the diffusive process is handled are provided in the paragraph 1.3.2.

Similarly to what done for the growth signal, the material and growth main directions (E2, KPar) are obtained through a diffusive process (with the exception of the L1 layer, which is set with a different method). More precisely the gradient of the diffusive process will be used, or, alternatively, the direction orthogonal to it. The process for assigning the directions is **Model/CCF/Fem Diffusion Anisotropy** and is formally identical to **Model/CCF/Fem Diffusion Growth**.

It is possible to handle growth and material anisotropy with a unique diffusive process, where the source with fixed high signal concentration is located at the central vertex (in the horizontal direction) at the mesh top, while the vertexes with fixed low concentration are located at the mesh boundary (excluding the top boundary): the outcome of this diffusive process produces a radial gradient pointing in the direction of the vertex with fixed high concentration (see Figure S3A, where the field is visible for the inner layers). For the

[2] Adimensional units adopted for diffusion as in the case of mass springs

mesh portion corresponding to the L1 layer, the material and growth anisotropy are set with a separate process, as it will be explained in the following.

At this point comes strictly mechanical part of the simulation. The mechanical elements need to be created by selecting all the faces of the mesh and by running the process **Model/CCF/04 Reference Configuration** which not only generates the elements for the FEM solver, but also sets their reference configuration as the current one. This process assigns also a thickness value, but this parameter is irrelevant in this 2D membrane simulation (so its has been left to 1 $\mu$m, as the membrane thickness won't affect the equilibrium configuration or the stresses, but will create a global rescaling factor for resultant forces).

Subsequently material properties get assigned. The template has a linear, hyperelastic, transverse isotropic, St. Venant material law, characterised by a special fiber direction (in this text named E2); for a detailed description of this material law and how it is used to compute the force vector for mechanical equilibrium, see Hofhuis et al. (2016).

As first thing, it is necessary to assign the intensity of the Young modulus in the fiber direction (E2) and in the isotropic plane (denominated E1E3 in the text: in the template this is represented by a vector, but conceptually it is a plane because of the virtual thickness). The process **Model/CCF/08a Set Aniso Dir** sets global values for the Young modulus and Poisson ratio. For the described template, it needs to be run with the field "Young E1E3" equal to "100", "E2" to "500" and "Poisson" equal to "0.3". An additional stiffness, negatively proportional to the growth signal previously computed through diffusion, is assigned to the E1E3 plane by running **Model/CCF/06b Material Properties based on Morphogens (for faces)** in all the models as it creates a sharper delimitation of the ovule protrusion w.r.t to the placenta (for a comparison see the outcomes at the git repository github.com/GabriellaMosca/FEM_2DOvuleGrowthModel/tree/master/constantStiffness)[3]

$$Y_{E1E3}+=\gamma\left(\frac{\text{MAX growth signal}-\text{growth signal}}{\text{MAX growth signal}-\text{MIN growth signal}}\right) \tag{1.14}$$

where $Y_{E1E3}$ is the Young modulus in the isotropic plane, $\gamma$ stands for the proportionality factor (set to 400), and growth signal is the signal obtained from the diffusive process **Model/CCF/Fem Diffusion Growth**(make sure to put the proper field signal name in the dialog box, its name was specified in Model/CCF/Fem Diffusion Anisotropy/05 Morphogen Visualize). The Young modulus distribution is the one show in Figure S3A.

For the all the template, except for the L1 layer, the special fiber direction (E2) is given by the vector orthogonal to the gradient of the diffusive field computed in **Model/CCF/Fem Diffusion Anisotropy**. To assign it this way, after selecting the face elements of the template, the user needs to run **Model/CCF/08b Set Aniso Dir Morphogens**, where, in the field "Direction Type" the key "E2" has been assigned, in the field "Direction from Diffusion Gradient" the key "Orthogonal to Gradient" and in the field "Morphogen Signal" the name chosen for the morphogenetic signal in **Model/CCF/Fem Diffusion Anisotropy**.

The L1 layer gets assigned a global E2 direction co-aligned with the y-axis (the vertical coordinate) by running the process **Model/CCF/08a Set Aniso Dir** with the accuracy of specifying direction type as "E2" and "Direction" as "0 1 0".

The same procedure is repeated to assign the growth anisotropy direction (Mosca et al., 2018), denominated KPar in this text, with the difference that the field "Direction Type" is "KPAR" and the "Direction from Diffusion Gradient" is "Parallel to Gradient". Regarding L1 layer, this is assigned a "Direction" of "1 0 0".

An edge-pressure acting on the template boundaries has been assigned (as the lateral and bottom boundaries will be constrained by Dirichlet condition, it is actually assigned only to its top boundary). This edge pressure acts exactly like the one described for the mass springs in Eq. 1.5 and has the same units. It has

[3]in the mass springs simulation the stiffness was instead assigned constant in the template, as a replication of what done with the FEM simulation produced a numerically unstable model (see github.com/GabriellaMosca/MassSpring_2DovuleGrowthModel/tree/master/variableStiffness. We argue that for mass spring models, the effect of softer cell wall is already implicitly encoded in the combination of signal based growth and passive strain based growth, so that adding such an effect explicitly exaggerates the dishomogeneity between faster and more slowly growing cells. In the FEM simulation, lacking cellular resolution, intead, the passive strain based growth is mostly acting as a relaxation to accumulation of stresses due to tissue conflicts.

been set to a value of 5 Kg/s$^2$ and its role is to simulate, to some extent, the action of turgor pressure into generating stresses in the tissue.

Dirichlet boundary conditions (as with mass springs) are assigned on the bottom and lateral boundary of the template by running the process **Model/CCF/15 Set Dirichlet** and blocking all the degrees of freedom (setting the field “Dirichlet Label” to “1 1 1”.

As last, it is required to assign the growth field. In the reference simulation (FEM-model 1) signal based growth is purely in the polariser direction (KPar) combined with strain-based growth (see following for explanation). This is assigned for the selected faces by running the process **Model/CCF/11b Set Growth Morphogens (on Faces)** after setting the field “KPar Scaling Factor” to 1, all the others to 1, and properly assigning the name in the field ”Morphogen Signal 1”.

##### 1.3.2 Diffusion simulation

Diffusion simulation is used in this model to generate a concentration field given some fixed concentration nodes. The problem is solved in 2D on a flat surface. It is slightly different from the one implemented for mass springs (see paragraph 1.1.3) as it is based on the FEM and starts from a continuous formulation originally expressed by this equation (heat equation):

$$D\nabla c(\mathbf{x}) = 0 \tag{1.15}$$

where $\nabla$ is the Laplace operator, $c$ the concentration of the diffusible substance and **x** are the coordinates used to described the space where diffusion is occurring. This problem requires boundary conditions (otherwise its solution would be trivial). For this diffusive process, unless prescribed Dirichlet condition (fixed concentration of the diffusing substance) have been assigned, zero flux condition is naturally assumed along the boundary. Internally to the domain, fixed concentration can be assigned as well, even if mathematically, this is equivalent to create an internal boundary (i.e. if inside a connected 2D surface a circle gets prescribed fixed concentration, which does not need to be constant in space, what will affect the solution outside this circle is only its boundary and not what has been set inside).

An explanation or introduction to the FEM is outside the purpose of these supplementary information, but there are several sources and manuals to consult, as for example Hughes (2012). In this case a direct solver method has been adopted, which means that the problem can be written, after the FEM discretisation, in the following way:

$$\mathbf{J}\mathbf{c} = \mathbf{B} \tag{1.16}$$

where **J** is a matrix, **c** is the vector of the nodal concentrations and **B** is a vector. The matrix **J**, is given, in components, is as follows:

$$J_{ij} = \begin{cases} \sum_{\mathfrak{T}\ni i} \sum_{j\in N_i} A_{\mathfrak{T}} \left(\mathbf{D}\phi_i \mathbf{D}\phi_j^T\right) & \text{if } c_i \text{ not fixed by Dirichlet} \\ 1 & \text{else if } i = j \\ 0 & \text{otherwise} \end{cases} \tag{1.17}$$

where the external sum is over all the triangles $\mathfrak{T}$ of the FEM mesh containing the node $i$, while the internal sum is over all the nodes $j$ which are connected to the node $i$. $A_{\mathfrak{T}}$ is the triangle area, $\mathbf{D}\phi_i$ is the vector of derivatives (in 2D) of the triangle linear basis functions $\phi_i$ (Hughes, 2012) referred to the triangle $\mathfrak{T}$(which has been omitted for simplicity of notation) and node $i$, explicitly:

$$\mathbf{D}\phi_i = \begin{pmatrix} \partial\phi_i/\partial X \\ \partial\phi_i/\partial Y \end{pmatrix} \tag{1.18}$$

with $X$ and $Y$ the local planar coordinates of the triangle. Regarding the **B** vector, it can be computed in components as follows:

$$B_i = \begin{cases} -\sum_{\mathfrak{T}\ni i} \sum_{j\in N_i} A_{\mathfrak{T}} c_j \left(\mathbf{D}\phi_i \mathbf{D}\phi_j^T\right) & \text{if } c_i \text{ not fixed by Dirichlet and } c_j \text{ fixed by Dirichlet} \\ 0 & \text{otherwise} \end{cases} \tag{1.19}$$

the final nodal concentration vector **c** $= (c_1, \cdots, c_m)$ is obtained as the solution to the matrix equation **Jc** $=$ **B** after adding to it the fixed nodal concentration values.

In this calculation the diffusion coefficient $D$ has been omitted as it does not affect the steady state concentration in this type of diffusion.

##### 1.3.3 One simulation cycle

A simulation cycle, which gets repeated in loop as many times (T) as specified by the user, consists of the following units:

- mechanical equilibrium simulation;
- growth step;
- mesh subdivision.

The template gets altered from a mechanical equilibrium condition because of the action of the edge pressure (assigned to the top margin) and of growth.

**Mechanical equilibrium simulation** After pressurization and each growth step, the system generally needs to find a new equilibrium configuration which accomodates the pressure induced stresses as well as the residual ones introduced by the growth process (see Rodriguez et al. (1994) and Goriely and Amar (2007) for an explanation of residual stresses).

As already introduced in Sec. 1.3.1, the material is transversely isotropic with a Sain-Venant Kirchhoff material law.

The equilibrium is computed by minimizing its strain energy by means of a semi-implicit Euler scheme (see Eq. 1.6). The elemental contribution to the nodal-force vector calculation is provided in detail in Hofhuis et al. (2016). The final nodal-force vector is obtained by summing for each node all the elemental contributions it is connected to. The Jacobian of the force vector is obtained through numerical derivation.

Details of material parameters and numerical convergence are provided in the Table in Figure S3A.

**Growth step** Growth is performed as a series of small deformations which are applied locally to the reference configuration and deform it without, directly, inducing stress or strain. As the deformation is local, no compatibility requirement is naturally satisfied and this has to be enforced by combining the growth deformation (which does not cause strain or stress) with an elastic deformation (which might induce strain and stress even in the absence of external loads), see Rodriguez et al. (1994).

The underlying hypothesis in this approach is that the timescale of growth is much bigger than that of elastic waves propagation, so that growth is supposed to occur in a quasi-static condition of mechanical equilibrium (Goriely and Amar, 2007).

In this formulation it is assumed (with no loss of generality) that all the rotational components of the growth tensor are absorbed in the elasticity tensor, so that growth is described purely by a stretch tensor, with no rotational component. In this description of growth, for each growth step, the reference configuration is the one obtained at the previous growth step and not the original one and it is called virtual as it might not ever occur physically (because of incompatibility). So, if $\mathbf{F}_E^n$ and $\mathbf{F}_G^n$ are the elastic deformation tensor and the growth deformation tensor at the $n$-th iteration, if $\mathbf{X}_{n-1}$ is the virtual reference configuration obtained at growth step $n-1$, the global deformation $\mathbf{F}_{EG}$ at the $n$-th growth step will be as follows:

$$F_{EG}^n = \mathbf{F}_E^n \mathbf{F}_G^n \mathbf{X}_{n-1} \tag{1.20}$$

In general the growth tensor $\mathbf{F}_G^n$ might depend on the current body coordinates $\mathbf{x}_{n-1}$ or on its state of elastic deformation/stress as computed from the previous virtual reference configuration. In the simulations in this paper, growth is constituted by an anisotropic part explicitly specified by the user through two diffusive process (one for the anisotropy direction $\mathbf{v}_{Kpar}$ and one for the growth intensity $KPar$, as described in Sec. 1.3.1) and a strain-based growth.

Growth is performed locally, by updating the rest length of the elemental triangles of the reference (possibly incompatible) configuration. It is possible to update this approach because triangles are simplexes, so their shape is fully determined by their side-lengths. The physical position of the reference triangle in space is not relevant.

Each triangle in the reference configuration stores in a ordered way the rest lengths of its edges and the angles made by the material fibers and growth anisotropy direction with its first side (the sides ordering is consistently preserved during the simulation). At the very first growth step, the growth direction and material anisotropy direction are naturally expressed in the reference configuration, so it is just a matter of computing the angle they make locally with the first triangle side. The growth anisotropy angle will be denominated $\alpha_{KPar}$, while the material anisotropy angle $\alpha_m$. The reference triangle is initially build from its ordered rest lengths in a local frame, where its first side is co-aligned and positively oriented with the x-axis. In this way

$$\mathbf{v}_{Kpar} = (cos(\alpha_{KPar}), sin(\alpha_{KPar})) \tag{1.21}$$

The growth tensor for this growth component, $\mathbf{G}_{KPar}$, is expressed in a diagonal basis:

$$\mathbf{G}_{KPar} = \begin{pmatrix} 0 & 0 \\ 0 & g_{KPar} \end{pmatrix} \tag{1.22}$$

where $g_{Kpar}$ is the growth coefficients for the local anisotropy growth in the direction parallel to $\mathbf{v}_{Kpar}$[4].

Consequently, it is necessary to rotate the reference triangle so that $\mathbf{v}_{Kpar}$ is representable in a canonical basis coordinates: given the explicit form of $\mathbf{G}_{KPar}$, as (0, 1).

The strain-based growth tensor $\mathbf{G}_E$, which will be summed up with the purely anisotropic one ($\mathbf{G}_{KPar}$) to constitute a global growth tensor $\mathbf{G}_{TOT}$, is expressed in the diagonal basis of $\mathbf{G}_{KPar}$. The strain here considered is the Green-Lagrange strain tensor, which is expressed in the current body configuration (more specifically it maps a vector in the reference configuration to a vector in the current one). The computation of $\mathbf{G}_E$ components proceeds as follows:

- the triangle represented in the current 3D configuration gets mapped by a rotation matrix in 2D;
- the Green-Lagrange strain tensor between the current triangle (mapped in 2D) and the reference one is computed (see Hofhuis et al. (2016) for how this is done) and eigenvalues and eigenvectors are derived;
- after computation of the engineering strain for each in-plane eigenvalue, the strain-based growth matrix in diagonal representation is built:

$$\mathbf{G}_E^D = g_\epsilon \begin{pmatrix} max(0, \epsilon_{11}^E - \epsilon_0^E) & 0 \\ 0 & max(0, \epsilon_{22}^E - \epsilon_0^E) \end{pmatrix} \tag{1.23}$$

  where $\epsilon_{11}^E$ and $\epsilon_{22}^E$ are the ordered engineering strain eigenvalues obtained from the Green-Lagrange strain eigenvalues $\epsilon_{11}$ and $\epsilon_{22}$ ($\epsilon_{ii}^E = -1 + \sqrt{(1 + 2\epsilon_{ii})}$), while $\epsilon_0^E$ is a threshold value below which no strain based growth occurs and $g_\epsilon$ is a growth proportionality factor[5];

[4]In general it is possible to specify growth also in the direction orthogonal to $v_{KPar}$, but, depending on the growth hypotheses made, this direction might be orthogonal to $v_{KPar}$ in the current configuration, but not in the reference one. Or it might loose the notion of orthogonality in both configurations if it is assumed that growth fields are specified at the beginning of the simulations and get deformed during growth and elastic deformation. This of course, only in the case anisotropic growth is coupled with another growth tensor which makes the final growth operator non-diagonal w.r.t to $v_{KPar}$ anymore. It will not be exposed here how the growth tensor should be computed in these cases.

[5]the engineering strain has been chosen due to its linear dependence on the stretch (for each principal direction)

- the strain-based growth tensor gets represented in the same basis which diagonalizes $\mathbf{G}_{KPar}$:

$$\mathbf{G}_E = \mathbf{R}_{eigen}^T \mathbf{G}_E^D \mathbf{R}_{eigen} \quad (1.24)$$

  where $\mathbf{R}_{eigen}^T$ is the matrix formed by the Green-Lagrange strain eigenvectors as columns.

The final growth tensor is then:

$$\mathbf{G}_{TOT} = \mathbf{1} + dt(\mathbf{G}_{KPar} + \mathbf{G}_E) \quad (1.25)$$

and it is expressed in the basis which diagonalises $\mathbf{v}_{Kpar}$. By applying it to the reference triangle coordinates, the triangles is grown (at the end only the rest lengths are stored, as the triangle arrangement in space is irrelevant).

It is now necessary to update the growth anisotropy and material angle with respect to the grown triangle first side, $\alpha_{KPar}$ and $\alpha_m$ respectively. The reference triangle configuration has been rotated so to be in the diagonal basis for $\mathbf{G}_{KPar}$, so the same rotation has to e applied to $\mathbf{v}_m$:

$$\mathbf{v}_m = \mathbf{R}_{KPar}\,(cos(\alpha_m), sin(\alpha_m)) \quad (1.26)$$

where by definition $\mathbf{R}_{KPar}$ is the rotation matrix which rotates $\mathbf{v}_{Kpar}$ into the vector represented by the coordinates (0, 1). At this point, by applying the final growth operator $\mathbf{G}_{TOT}$ to the two vectors, one obtains their orientation (plus an irrelevant stretch) in the new grown configuration, so it is possible to compute the updated angles between these two vectors and the first, consistently oriented, triangle side.

Underlying this approach there is the idea that the anisotropy direction got computed at the beginning of the simulation and gets passively deformed by the growing rest configuration. An alternative approach would have been to re-compute the diffusion process and its gradient at each growth step on the current configuration and then map it on the rest configuration. In this specific simulation scenario, it is possible to verify that the two gradients so obtained do not differ significantly.

**Mesh subdivision** When a triangle area in the current configuration exceeds a pre-set threshold (Max Triangle Area in Table in Figure S3A), it gets divided by a bisection algorithm which splits its longest side in two segments of equal length and generates so two new triangles (which replace the old one) by connecting the newly inserted vertex to the opposed one. To ensure a conformal mesh, the subdivision is propagated to neighbour triangles as described in Rivara and Inostroza (1995). After each subdivision the triangle, edge and vertex properties need to be propagated

- the **vertex** properties which need to be propagated to the newly inserted vertex are the Dirichlet condition (both for mechanics and diffusion): if the new vertex has been inserted between two vertexes with Dirichlet conditions assigned, it will inherit them, otherwise no;
- an **edge** in the current configuration gets divided in two equal parts, so the new edges inherit half the rest length of the undivided one each;
- by splitting an edge, the initial **triangle** is split in two smaller triangles (not necessarily identical): each of them inherit the same triangle properties of the undivided one (i.e. material properties, growth properties, anisotropy angles, etc.).

**Robustness analysis** As done with mass springs, a robustness analysis around the reference model (FEM-Model 1) has been performed by varying independently stiffness and intensity of growth signal of 10% around the reference values. As can be observed by examining the results in the git repository github.com/GabriellaMosca/FEM_2DOvuleGrowthModel/tree/master/robustnessAnalysis such variations do not affect significantly the outcome, confirming the stability of the reference model.

#### 1.4 3D analysis of ovule morphology and connectivity

##### 1.4.1 Ovule segmentation export from IMARIS

A in-house tool (ExportImarisCells, git repo github.com/barouxlab/ExportImarisCells) has been developed to export segmentations performed in the proprietary software IMARIS (imaris.oxinst.com/. This tool, to be launch within IMARIS, creates a cell-mask with a different integer value for each segmented cell and the value zero for everything else. All the cell-masks are summed algebraically together to provide a final mask where each cell (each 3D voxel occupied by a cell) is labeled with a different integer value and the outside space is labeled with the value 0. This algorithm is based on the hypothesis that cells occupy mutually exclusive space regions. It is possible to export with different labels $2^{16}$ different cells. The output of this operation is saved as a tiff images series.

##### 1.4.2 3D mesh cell-based and surface mesh creation in MorphoGraphX

After exporting the segmentation as a tiff-file series from IMARIS in the previous step, it is now possible to load it in MorphographX (de Reuille et al., 2015) (www.MorphoGraphX.org). The user needs to specify manually the z voxel size. At this point the stack is loaded and visible in MorphoGraphX and the option 16 bit as well as label need to be selected to proceed properly. Upon running the process **Process/Stack/Segmentation/Relabel** the image series will be relabeled and can be saved as a mgxs file, a native format in MorphoGraphX for segmented stacks. After this step, it is possible to properly generate the cell mesh by running the process **Process/Mesh/Creation/Marching Cubes 3D**, where the cube size, which is related to the mesh refinement, has to be specified by the user (a size of 0.5 $\mu^2$ has been specified for meshes connected to ovules till Stage 1-I, for later stages, to avoid underestimation of surface of contact between cells, a size of 0.1 $\mu^2$ has been adopted). This provides a cell mesh which can already be used for volume quantifications. As a further step, to be able to use it in MorphoMechanX, the cell mesh needs to be converted and saved as a 3D tissue by running the process: **Process/Mesh/Cell Mesh/Convert to 3D Cell Tissue** and then saving the result. Both mesh types can be saved in the native MorphographX mesh format mgxm. Starting back from the saved relabeled stack (the mgxs file), by following the instructions provided in the MorphoGraphX user manual (available here www.mpipz.mpg.de/4085950/MGXUserManual.pdf) it is possible to generate a surface mesh and by running **Process/Mesh/Signal/Project Mesh Curvature** the curvature signal will be projected on it. The curvature used in the analysis reported in Figure 1E-F is the minimal curvature, with manually set max and min range (for Stages 0-I and 0-II the range has been set between -0.15 and 0.15, while for later stages, as the ovule profile gets sharper, between -0.25 and 0.25).

##### 1.4.3 Connectivity analysis for the ovule layers

The 3D Cell Tissue mesh saved as described in the previous section can be loaded in MorphoMechanX to perform a semi-automated cell layer classification and connectivity analysis. An in-house tool to be run within MorphoMechanX has been developed to be able to detect semi-authomatically cell layers, after the user selects manually the L1 cells. If the user selects as well explicitly the pSMC/MMC, this will be marked as so in the analysis ad the companion cells will be identified (both cell types are considered sub-cases on L2 layer cell classification). The tool is available at the git repository: github.com/GabriellaMosca/3DsemiAutoLabelingOvule.

A cell is identified to be a L2 cells if it shares a portion of its wall with the L1 layer. To avoid spurious contacts due to segmentation imperfection, a minimal threshold for contact can be specified (in this case 0.01 $\mu^2$). Once the L2 cells have been identified (the pSMC/MMC and companion cells are considered L2 cells), the L3 cells will be detected as the cells sharing a portion of their wall with L2, but none with L1. Finally all other cells are gathered in a global group. This tool will save to a file, for each selected cell, its labeling (L1, L2, pSMC/MMC, L3, etc.), its volume and in case of L2 cell (including pSMC/MMC cell) its surface of contact with L3 layer. In the file are furthermore saved the three eigenvalues of the positional covariance matrix described in the next section. For a tutorial on how to use the tool, see the README.md file in the git repository.

##### 1.4.4 MMC shape characterization with its positional covariance matrix

Following the idea presented in Sec. 1.2 of this document, a 3D version of the covariance matrix of Eq. 1.11 has been used to characterize anisotropy of pSMC/MMCcell shape in 3D. The computation, analogously with what done for the 2D case, is performed along the cell surface and the already existing mesh is used for the point distribution. The points used for the computation are the centroids of each mesh triangle and the covariance matrix, with elements area as weights to compensate for dis-homogeneities in the mesh looks like:

$$\textbf{Cov} = \frac{1}{\sum_i^N A_i} \begin{pmatrix} \sum_{i=0}^N A_i(x_i - x_0)^2 & \sum_{i=0}^N A_i(x_i - x_0)(y_i - y_0) & \sum_{i=0}^N A_i(x_i - x_0)(z_i - z_0) \\ \sum_{i=0}^N A_i(x_i - x_0)(y_i - y_0) & \sum_{i=0}^N A_i(y_i - y_0)^2 & \sum_{i=0}^N A_i(y_i - y_0)(z_i - z_0) \\ \sum_{i=0}^N A_i(x_i - x_0)(z_i - z_0) & \sum_{i=0}^N A_i(y_i - y_0)(z_i - z_0) & \sum_{i=0}^N A_i(z_i - z_0)^2 \end{pmatrix} \tag{1.27}$$

in this formula $N$ is the total element number (not the number of nodes), $A_i$ is the element area, $x_i,\ y_i,\ z_i$ are the elements centroids and $x_0,\ y_0,\ z_0$ are the overall 3D shape centroid (computed as the weighted average of the element centroids positions).

In this case the three eigenvectors of the covariance matrix correspond to (i)the axis where the variance of the points projection on it is maximal, (ii) the axis where it is minimal and (iii) a third orthogonal direction to the other two where the variance is intermediate.

To get a proxy of cell anisotropy (so independent from the cell size) it is possible to rescale the eigenvalues of the covariance matrix by dividing each for the sum of all three (making them a partition of unity):

$$c_i^{rescaled} = \frac{c_i}{\sum_{j=1}^3 c_j}$$
