## Supplementary material for "Organ geometry channels cell fate in the Arabidopsis ovule primordium": Source Data 1

### Supplemental Dataset 1- Gallery of wild-type (Col_0) ovules used for the 3D digital atlas related to Figure 1

**Number of ovules segmented for the analysis: 92** *from 84 images*

| | |
|---|---|
| Stage 0-I | n=21 |
| Stage 0-II | n=17 |
| Stage 0-III | n=11 |
| Stage 1-I | n=15 |
| Stage 1-II | n=11 |
| Stage 2-I | n=10 |
| Stage 2-II | n=7 |

### Stage 0-I (1/2)

EM_C_25

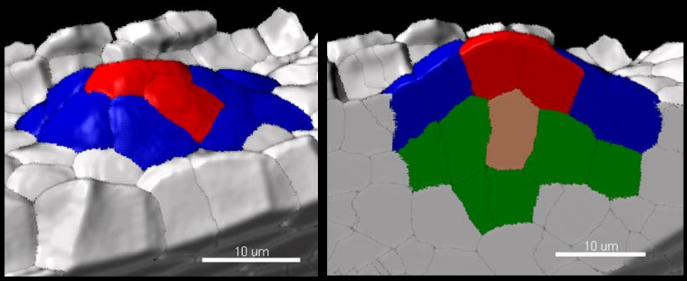


EM_C_149

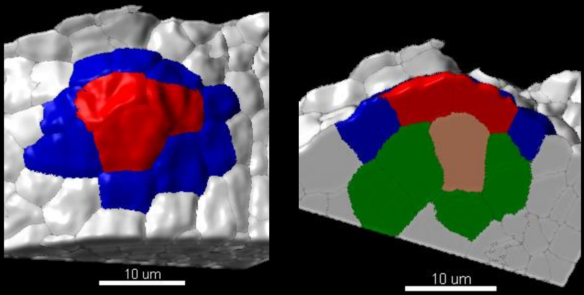


EM_C_58

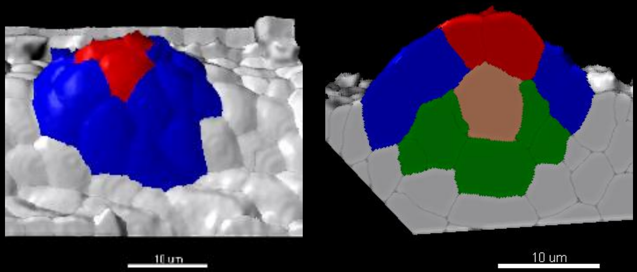


EM_C_237

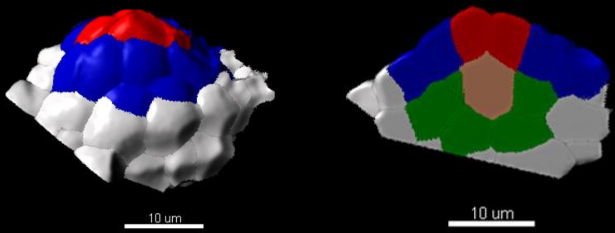


EM_C_59

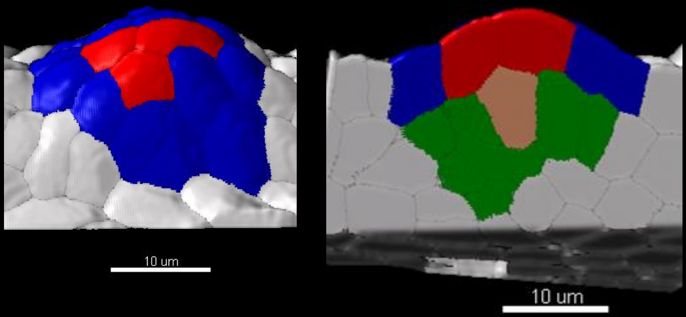


EM_C_240

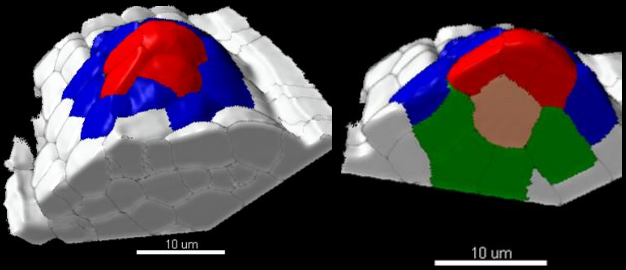


EM_C_60

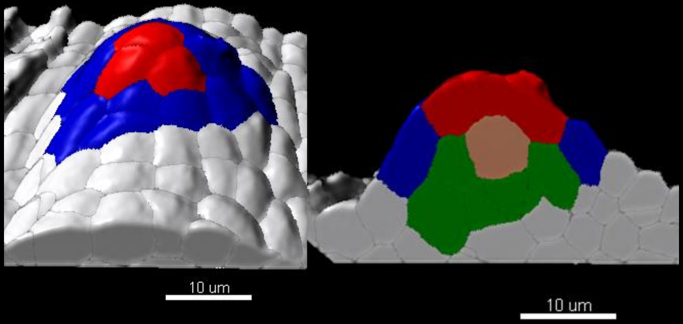


EM_C_241

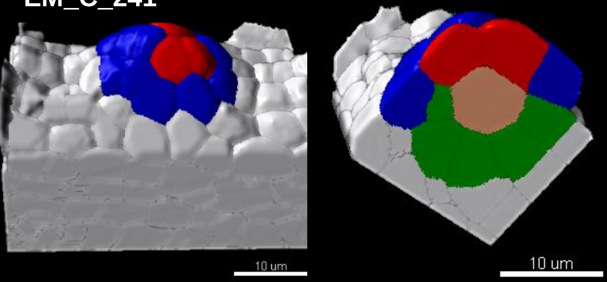


EM_C_76

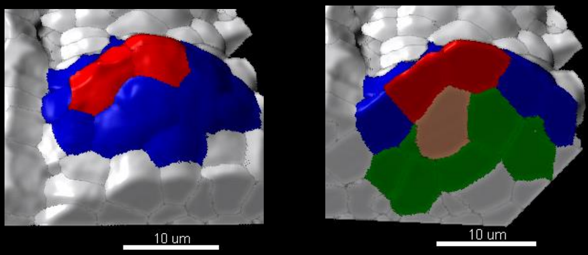


EM_C_242A

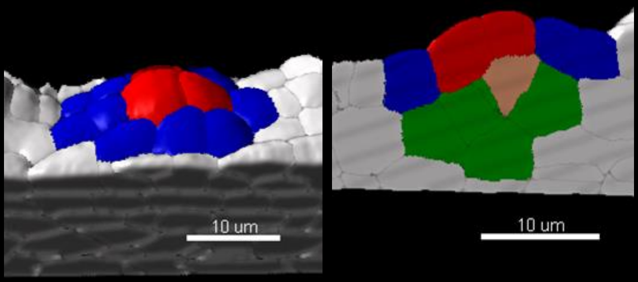

### Stage 0-I (2/2)

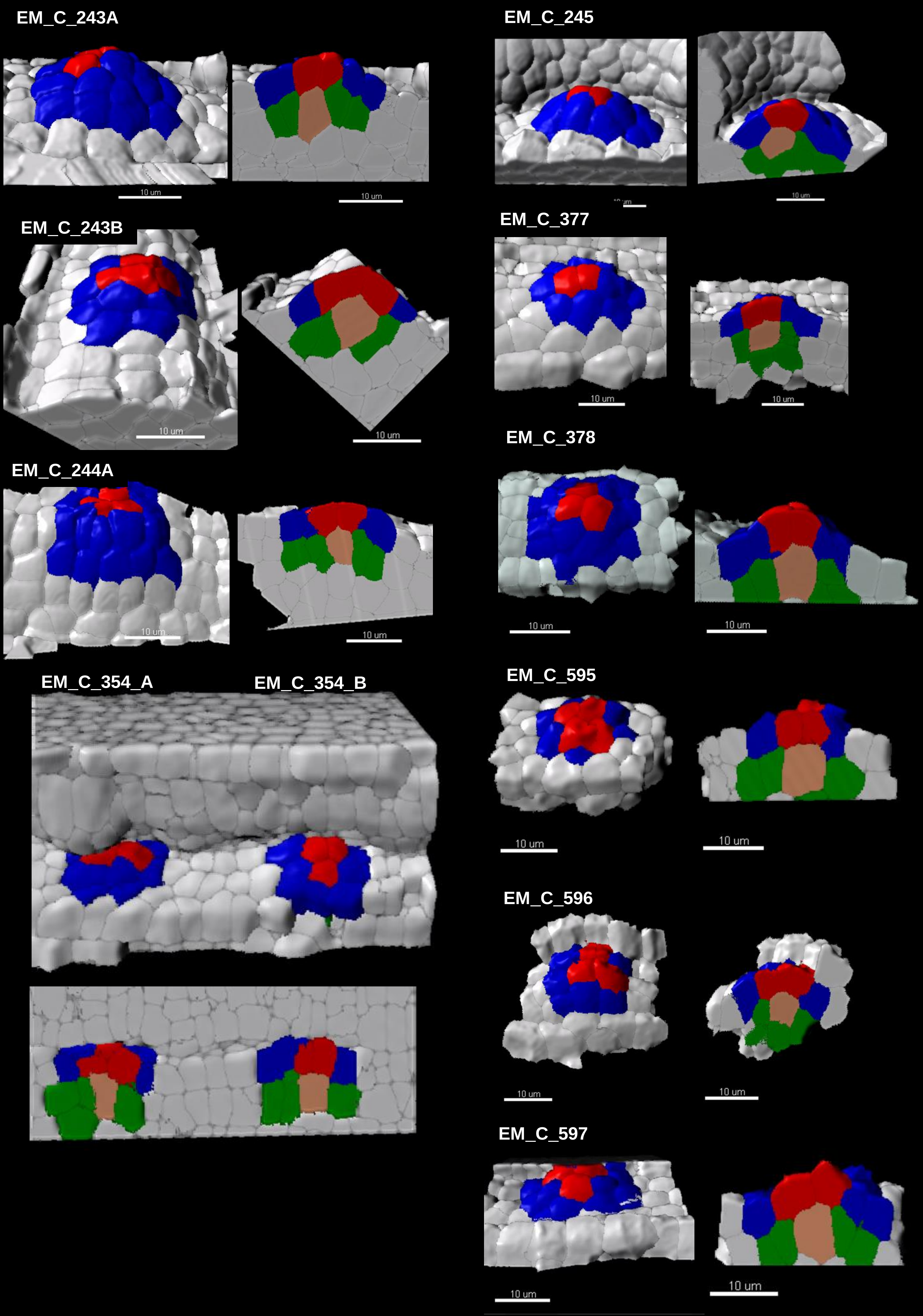
EM_C_243A
EM_C_245
EM_C_243B
EM_C_377
EM_C_244A
EM_C_378
EM_C_354_A
EM_C_354_B
EM_C_595
EM_C_596
EM_C_597
10 um

### Stage 0-II (1/2)

EM_C_22

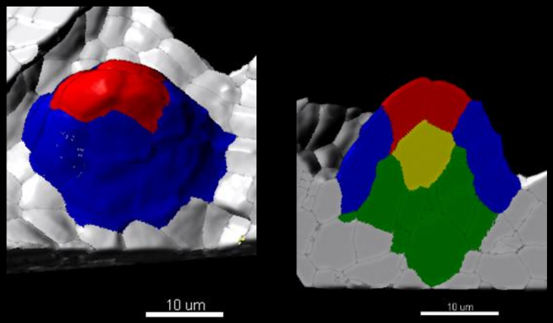


EM_C_57

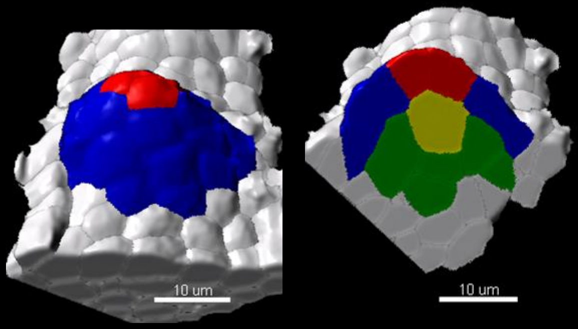


EM_C_61

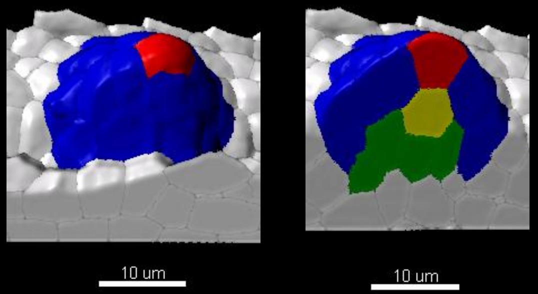


EM_C_75

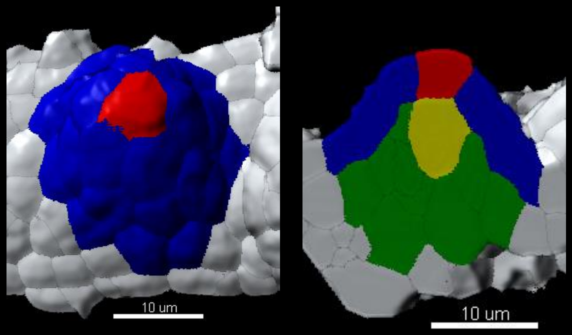


EM C 77

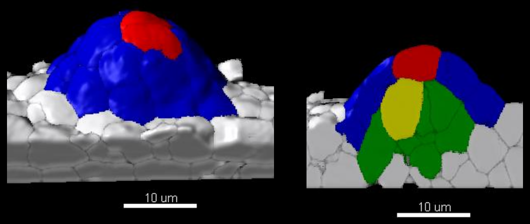


EM_C_78

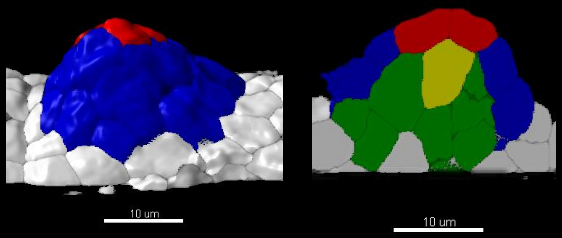


EM_C_146

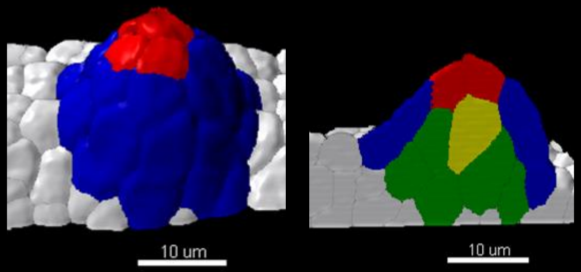


EM_C_148

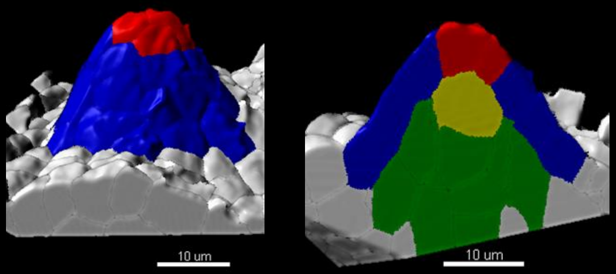


EM_C_150

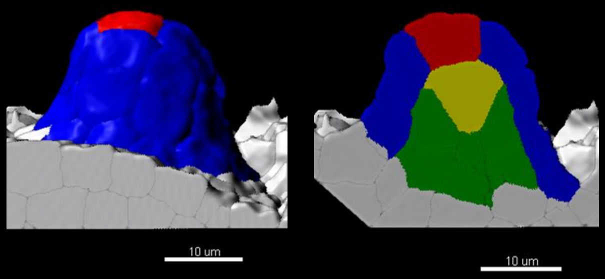

### Stage 0-II (2/2)

EM_C_211

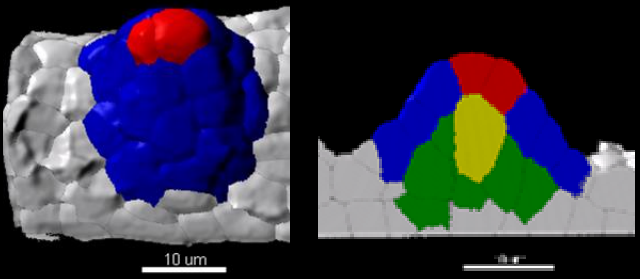


EM_C_214

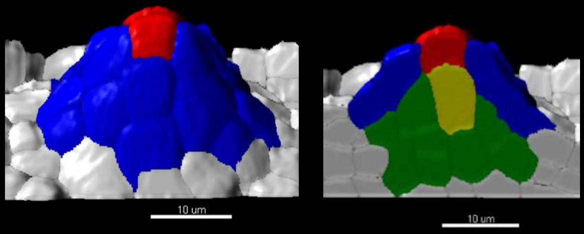


EM_C_212A

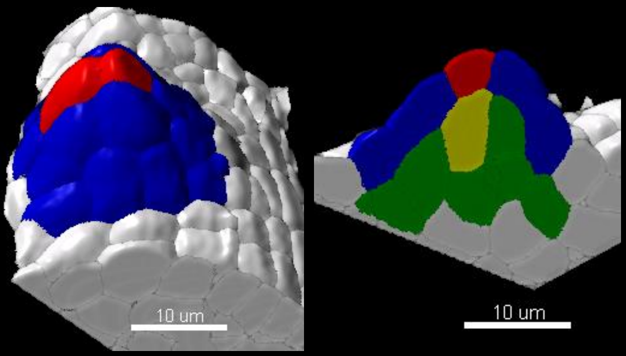


EM_C_238

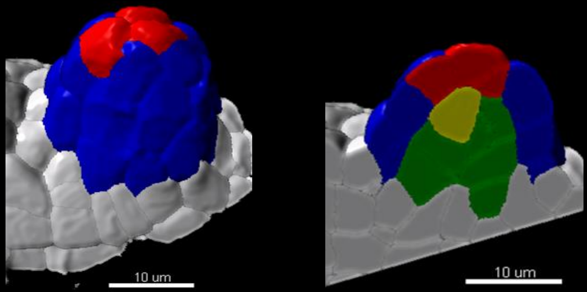


EM_C_212B

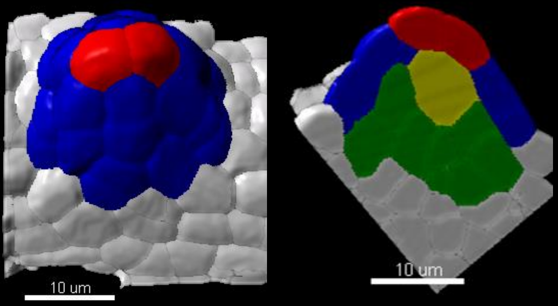


EM_C_239

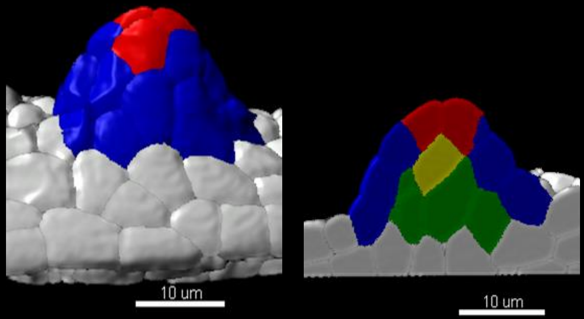


EM_C_213

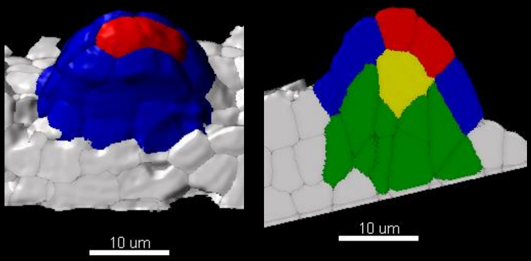


EM_C_246

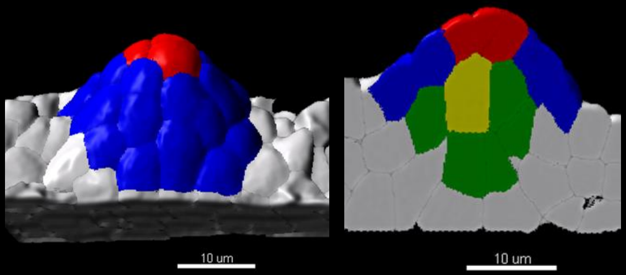

### Stage 0-III

EM C 136

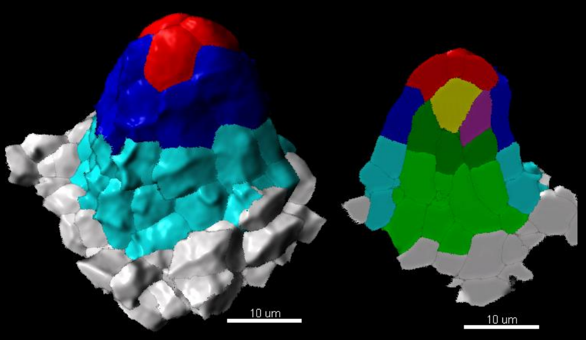


EM_C_138

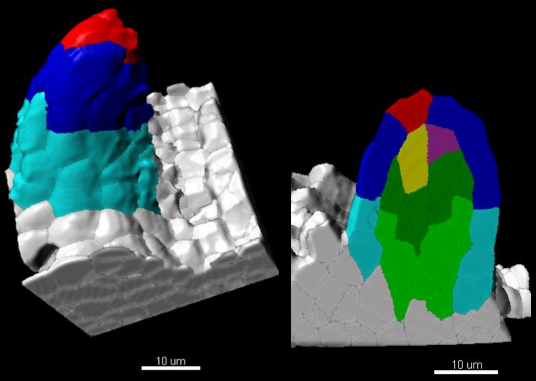


EM_C_139

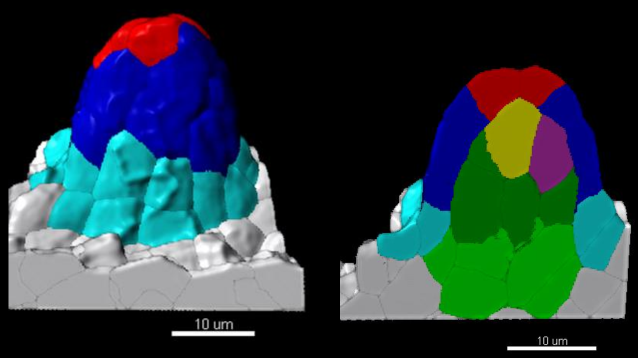


EM_C_140

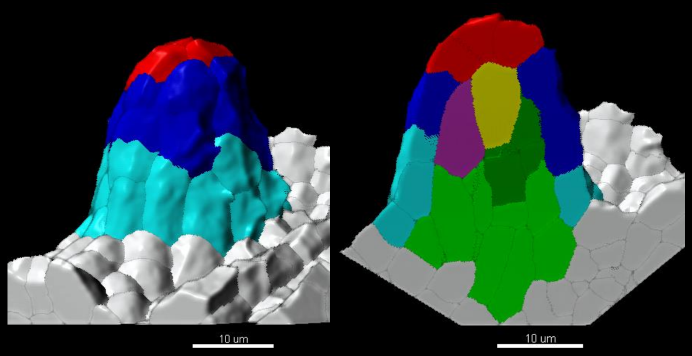


EM_C_141

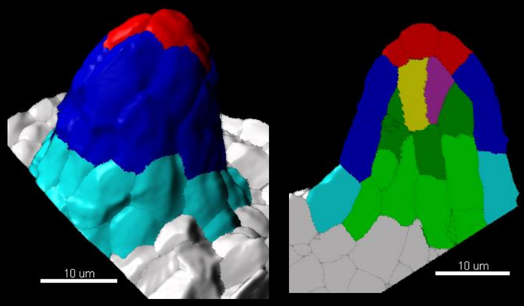


EM_C_142

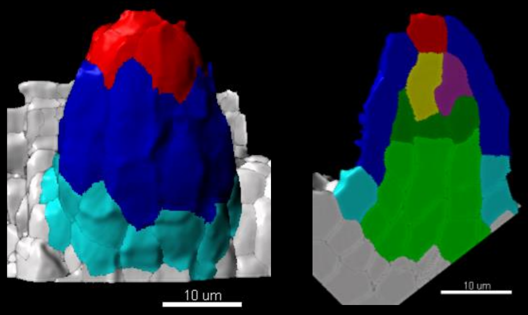


EM_C_187

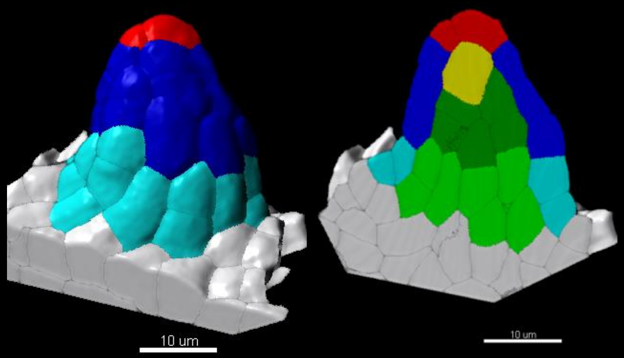


EM_C_403

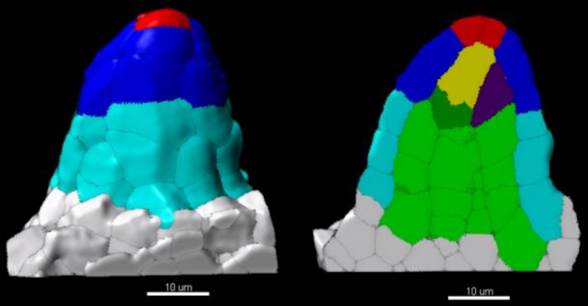

### Stage 1-I (1/2)

EM_C_6

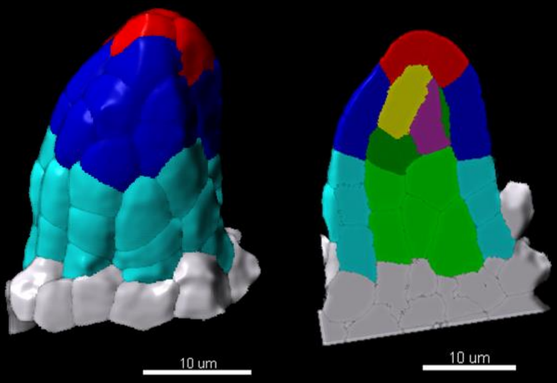


EM_C_189

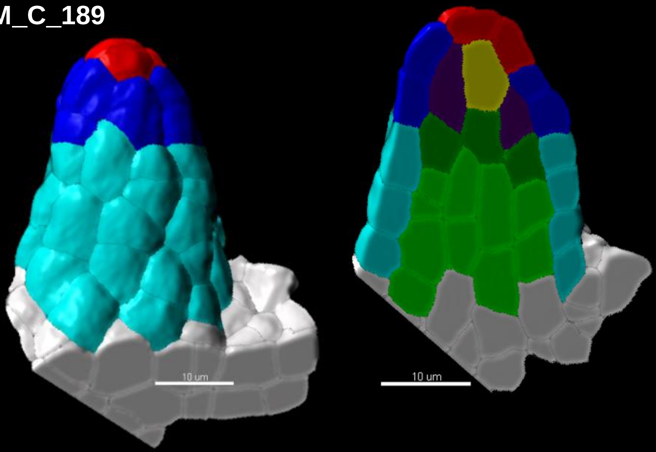


EM_C_185

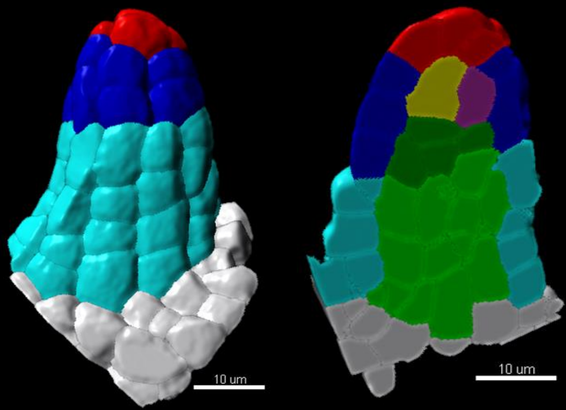


EM_C_196

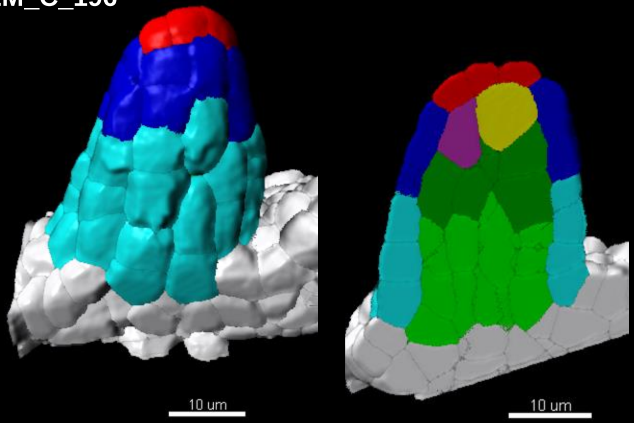


EM_C_186

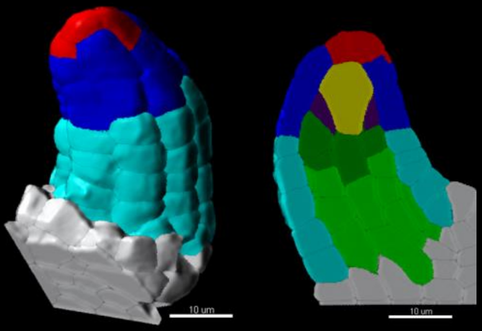

EM_C_198

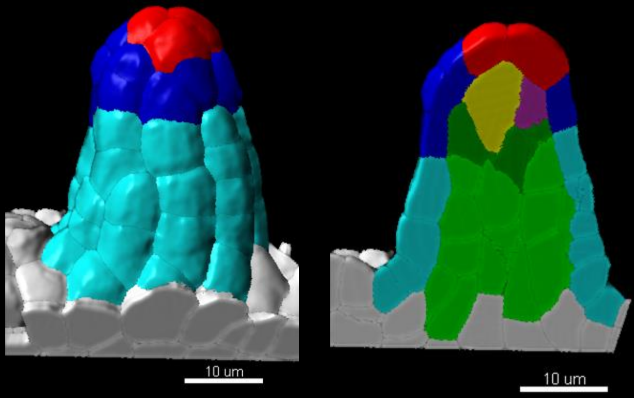


EM_C_188

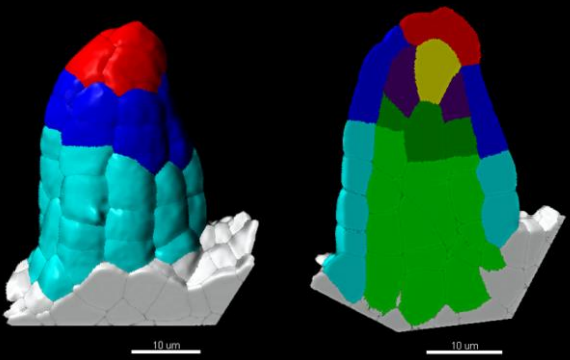


EM_C_200

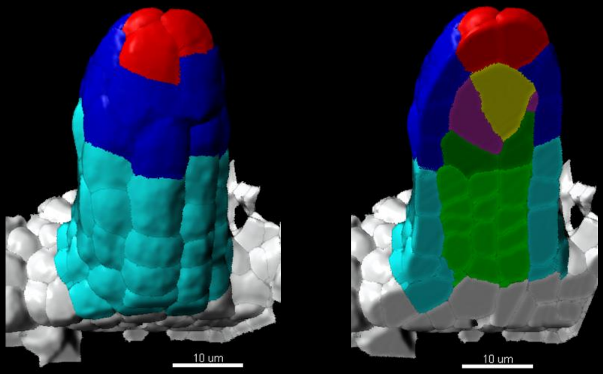

### Stage 1-I (2/2)

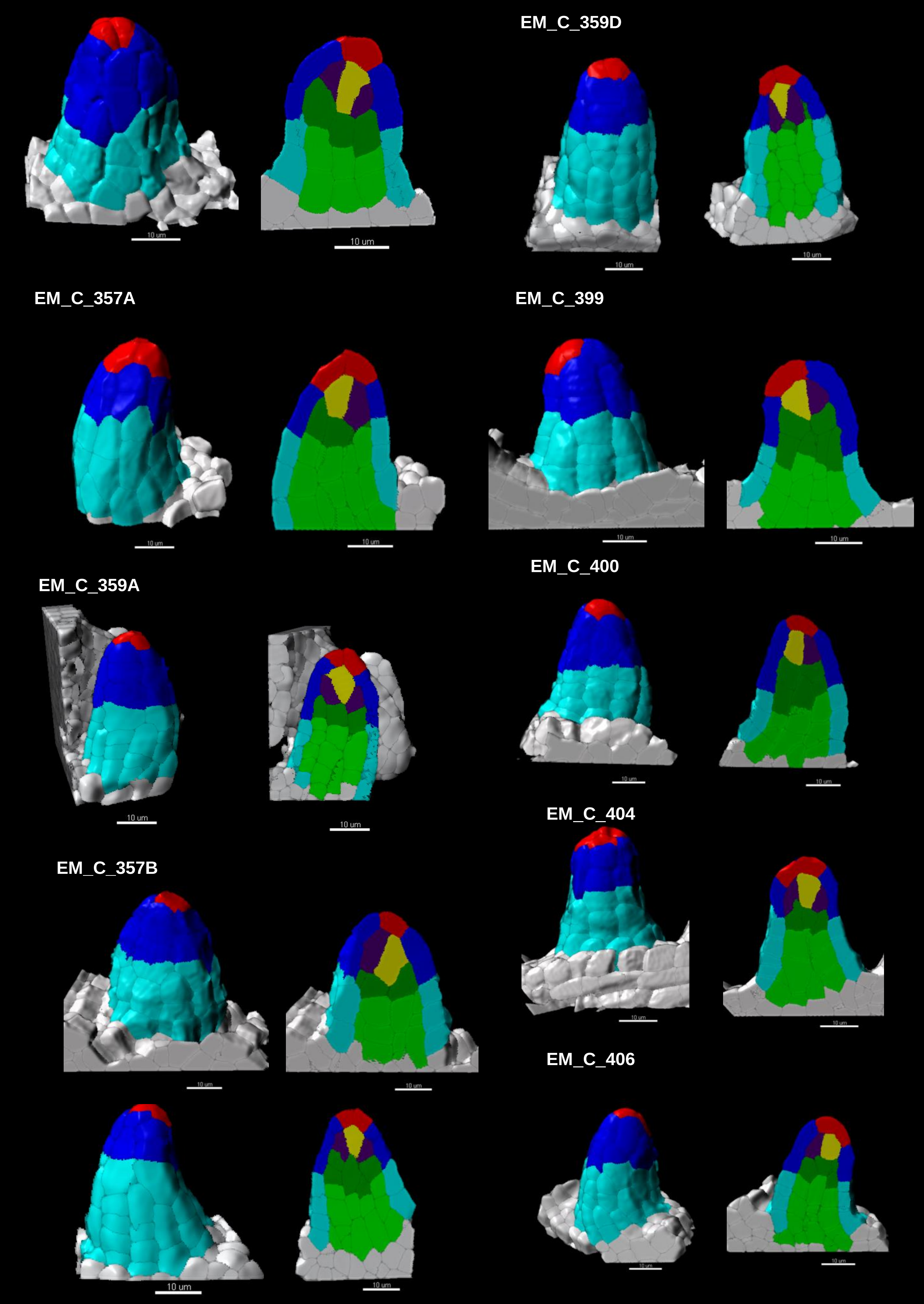
EM_C_202
10 um
10 um
EM_C_359D
10 um
10 um
EM_C_357A
10 um
10 um
EM_C_399
10 um
10 um
EM_C_359A
10 um
10 um
EM_C_400
10 um
10 um
EM_C_404
10 um
10 um
EM_C_357B
10 um
10 um
EM_C_406
10 um
10 um
10 um
10 um

### Stage 1-II (1/2)

EM_C_96

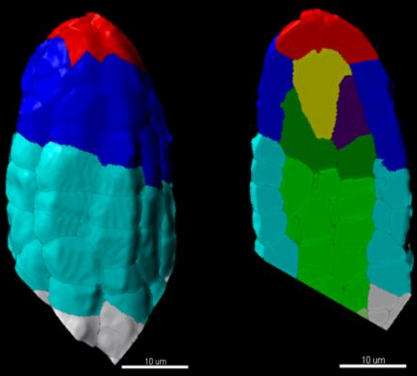


EM_C_102A

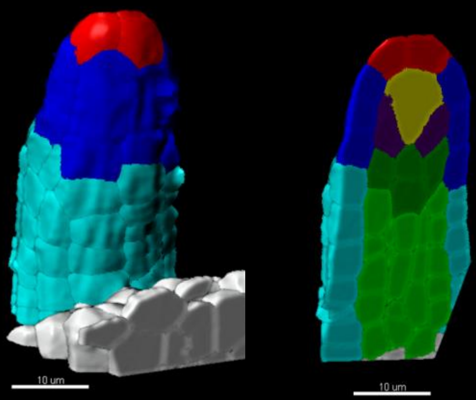


EM_C_112

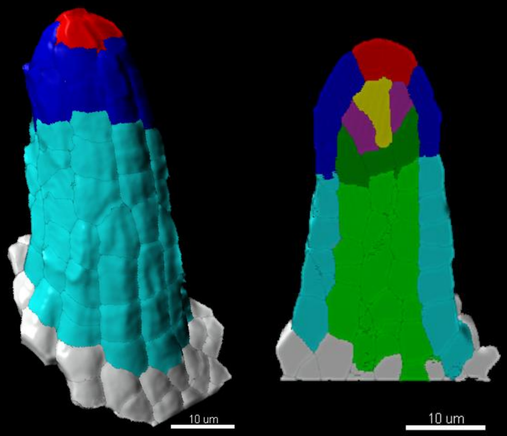


EM_C_162

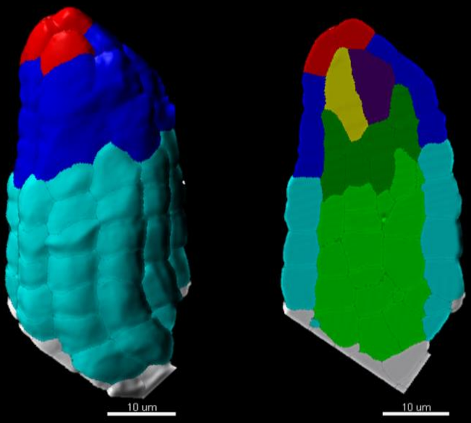


EM_C_184

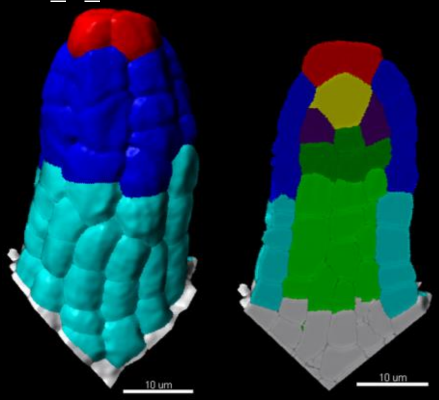


EM_C_199

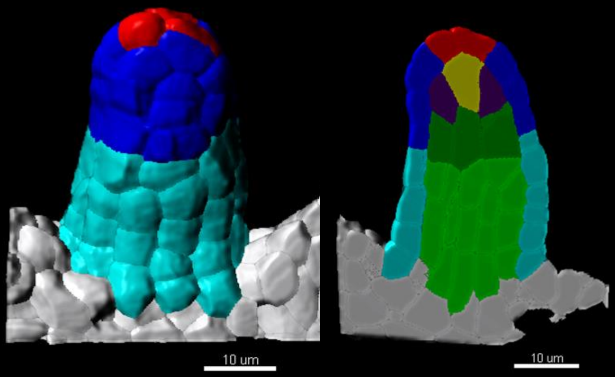


EM_C_201

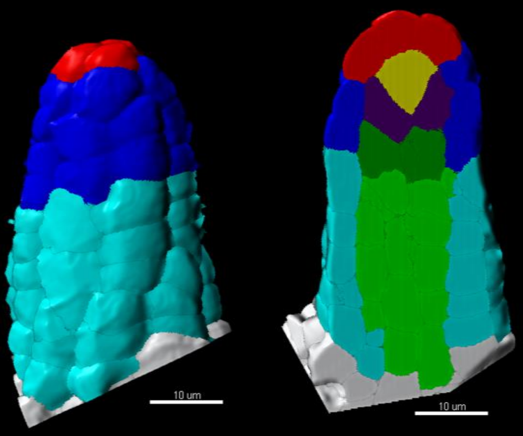


EM_C_204

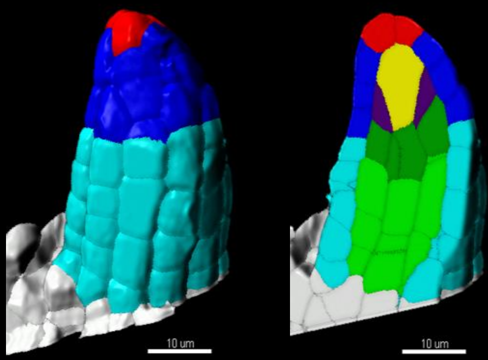

### Stage 1-II (2/2)

EM_C_359B

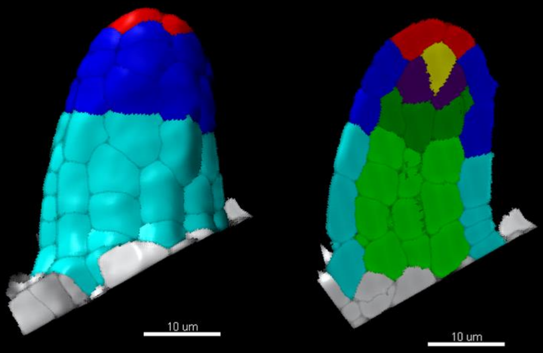


EM_C_607

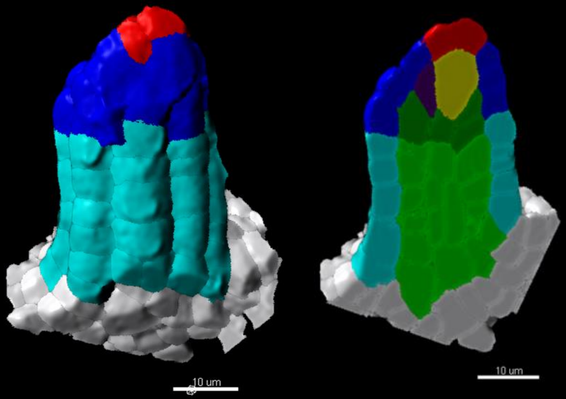


EM_C_160

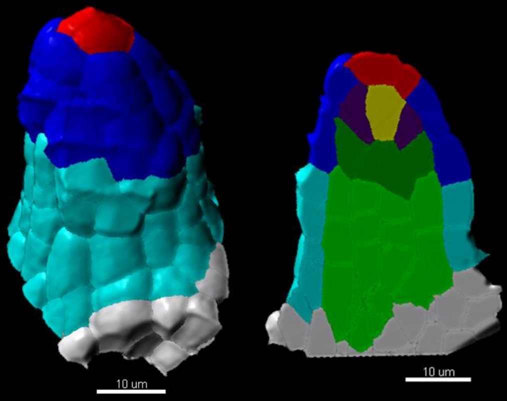

### Stage 2-I

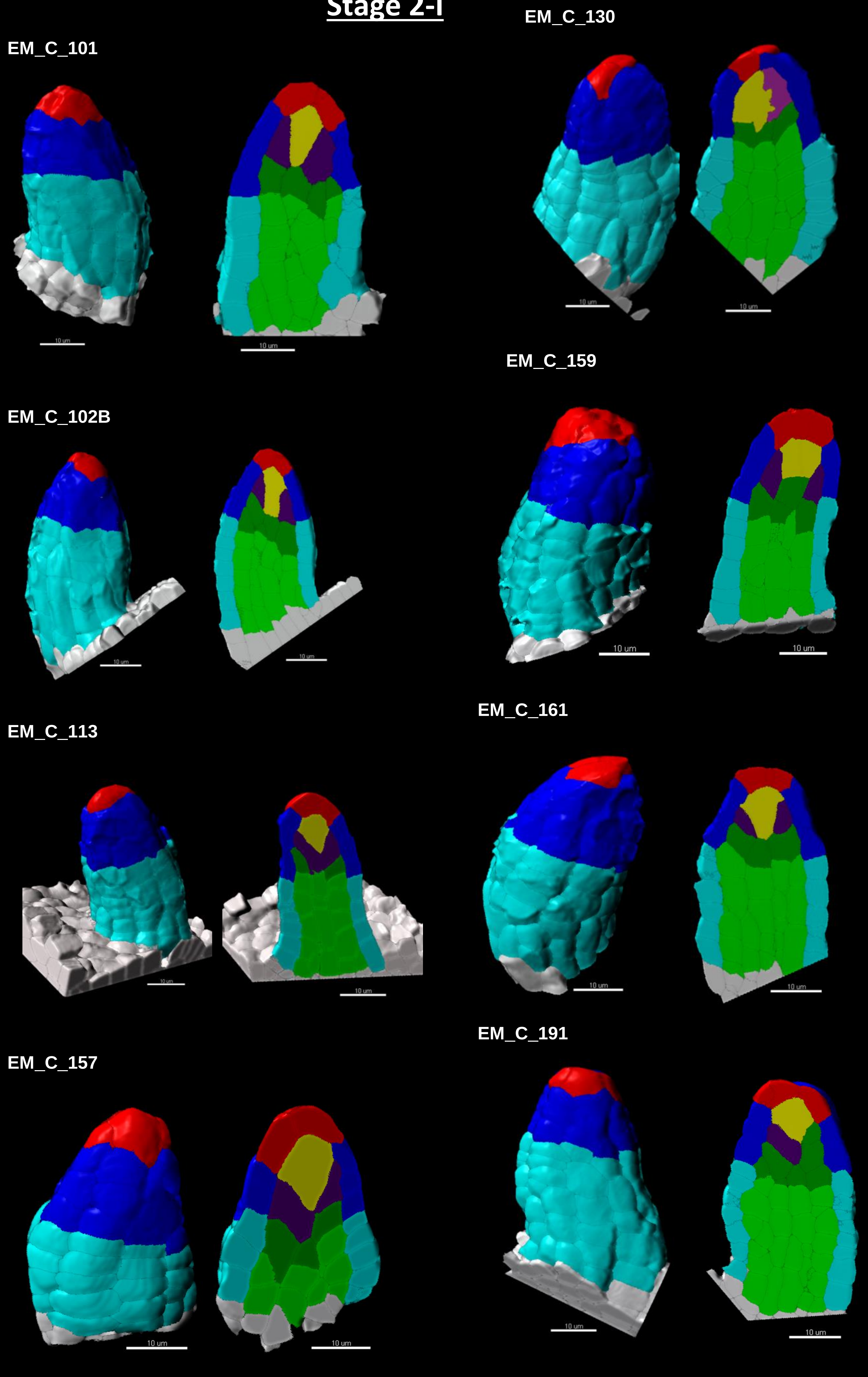
EM_C_101
EM_C_130
EM_C_159
EM_C_102B
EM_C_113
EM_C_161
EM_C_157
EM_C_191
10 um

### Stage 2-I

EM_C_229

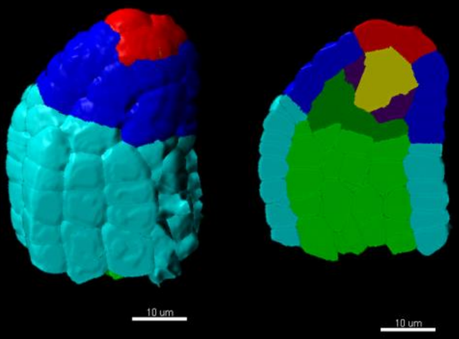


EM_C_232

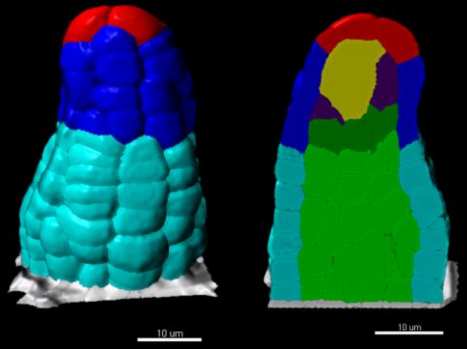

### Stage 2-II

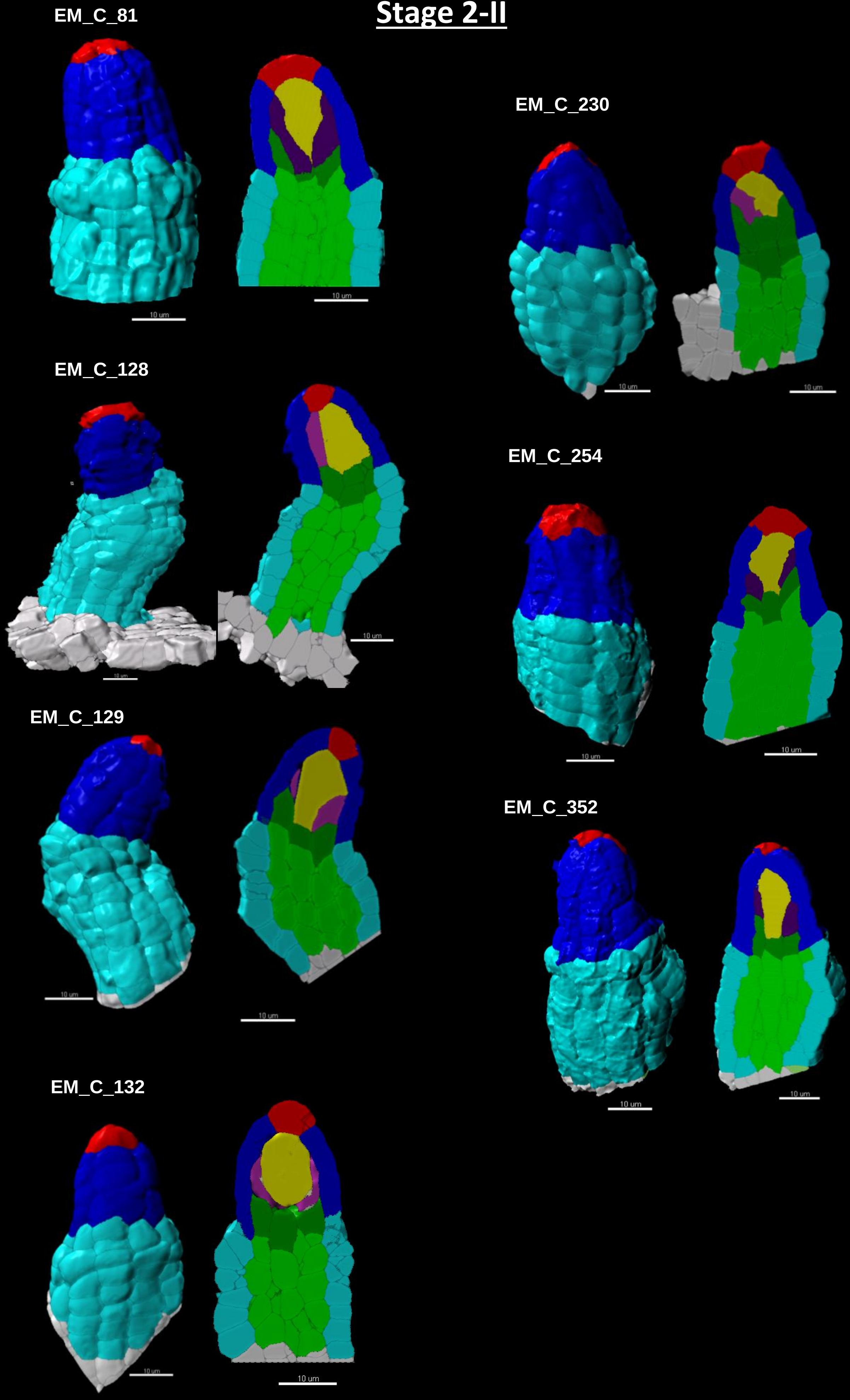
