## Supplementary material for "Organ geometry channels cell fate in the Arabidopsis ovule primordium": Source Data 4a

### Supplemental Dataset 4a- Gallery of *katanin* (*bot1-7*) and corresponding wild-type (*Ws-4*) ovules used for the 3D digital atlas related to Figure 4

**Number of ovules segmented for the analysis**

| | *bot1-7* | Ws-4 |
|---|---|---|
| Stage 0-III | n=16 | n=7 |
| Stage 1-I | n=9 | n=8 |
| Stage 1-II | n=12 | n=7 |

**bot1-7**

### Stage 0-III

## EM_C_746

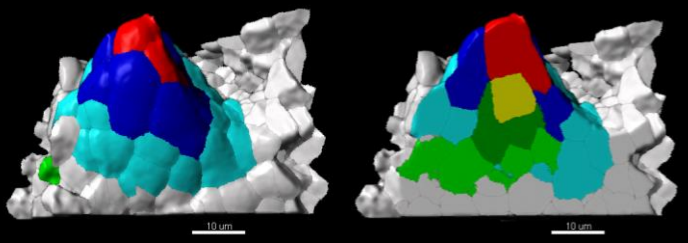


## EM_C_744A

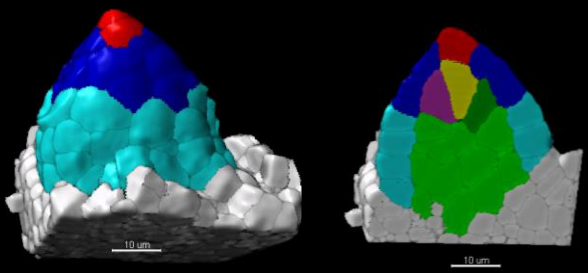


## EM_C_745B

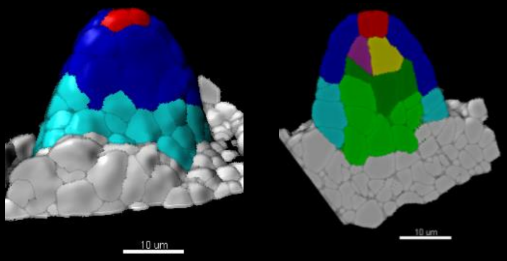


## EM_C_572

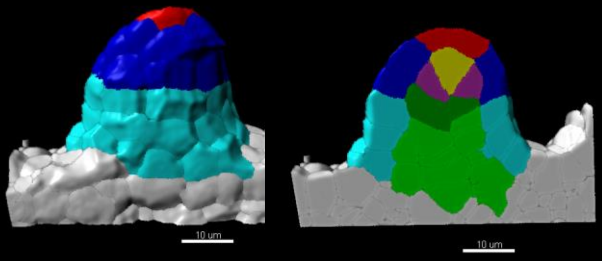


## EM_C_573

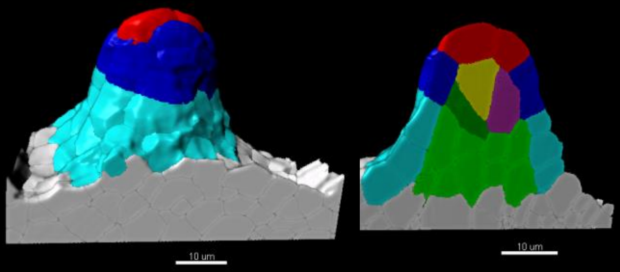


## EM_C_571

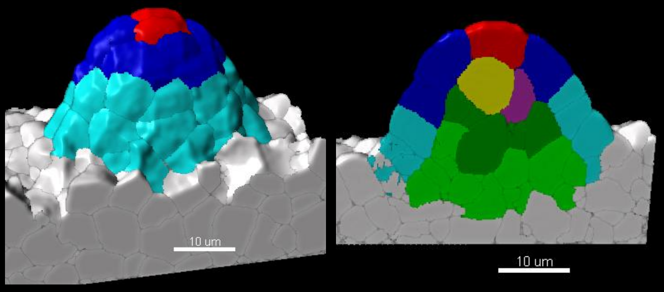


## EM_C_574

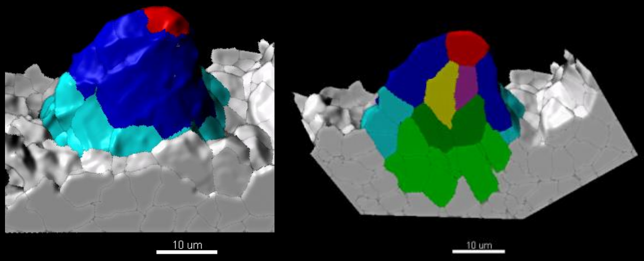


## EM_C_753

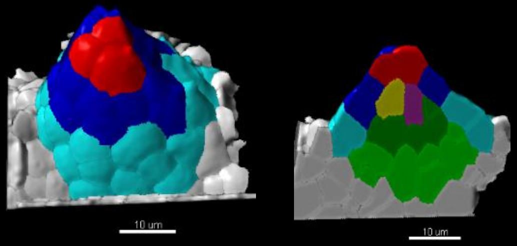


## EM_C_754A

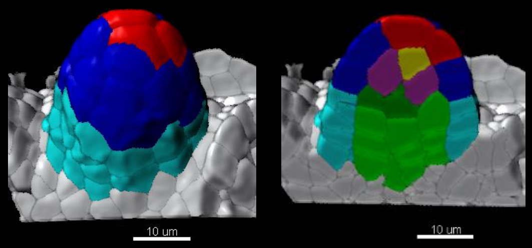


## EM_C_754B

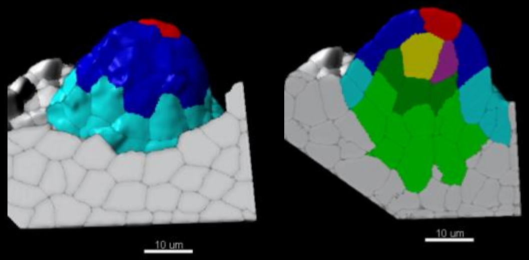

*bot1-7*

### Stage 0-III

## EM_C_280A

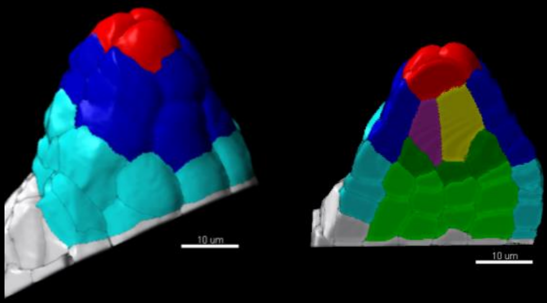


## EM_C_755B

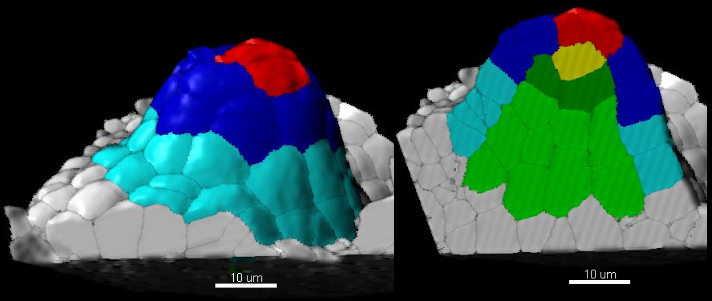


## EM_C_792B

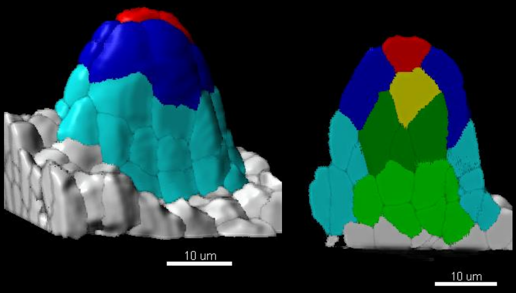


## EM_C_790

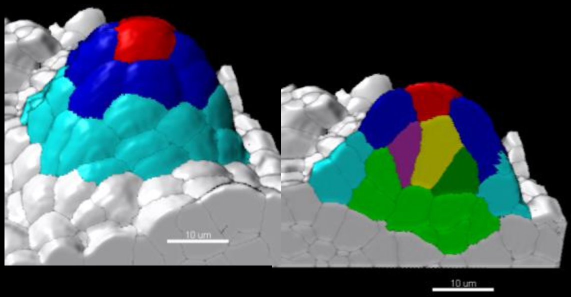


## EM_C_777

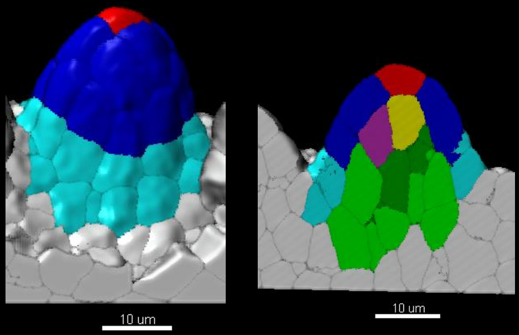


## EM_C_779

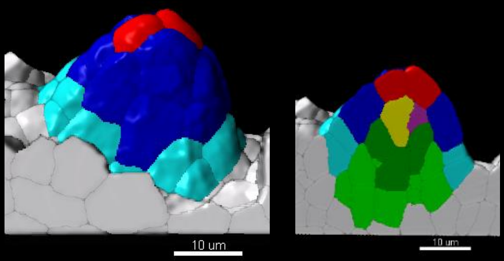

***bot1-7***

### Stage 1-I

EM_C_743A

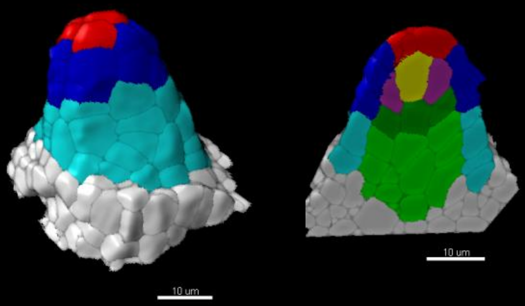

EM_C_757

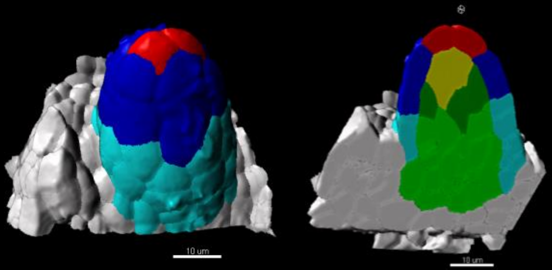

EM_C_752

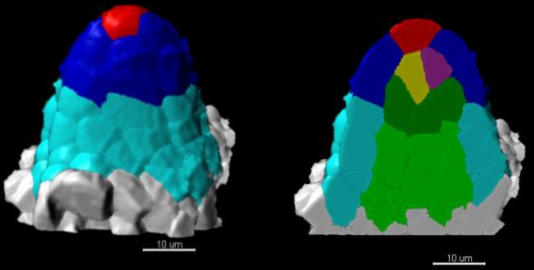

EM_C_590

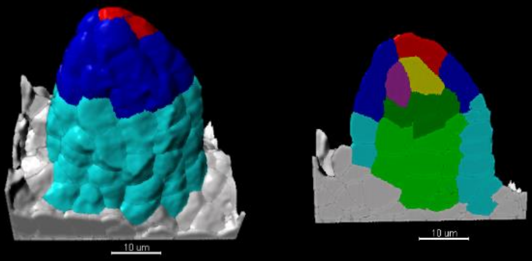

EM_C_921

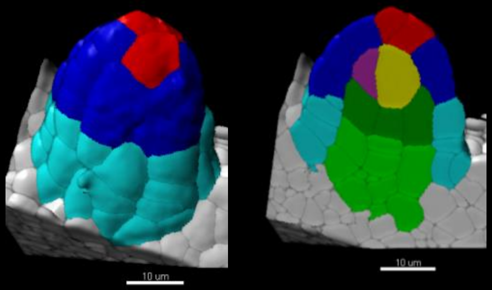

EM_C_913

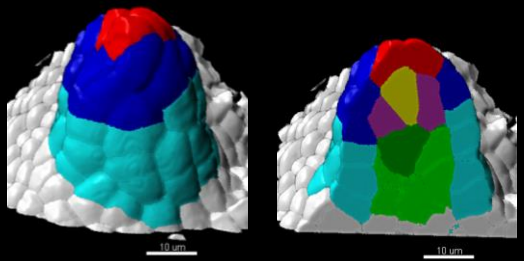

EM_C_783

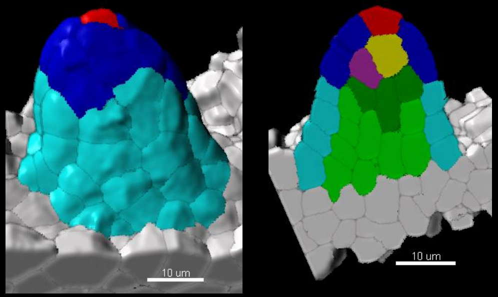

EM_C_782A

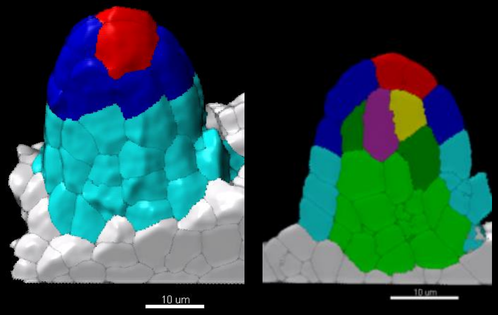

EM_C_778

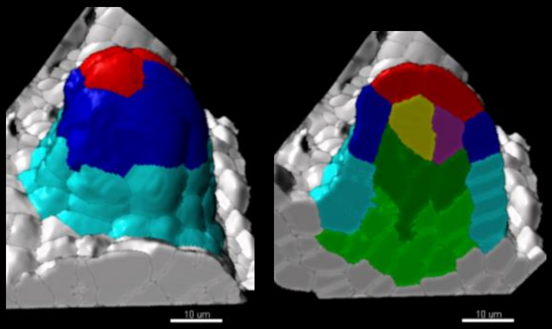

*bot1-7*

### Stage 1-II

EM_C_914

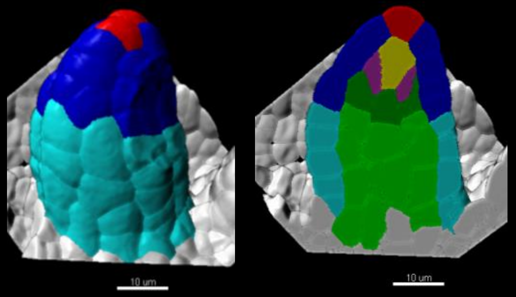


EM_C_747

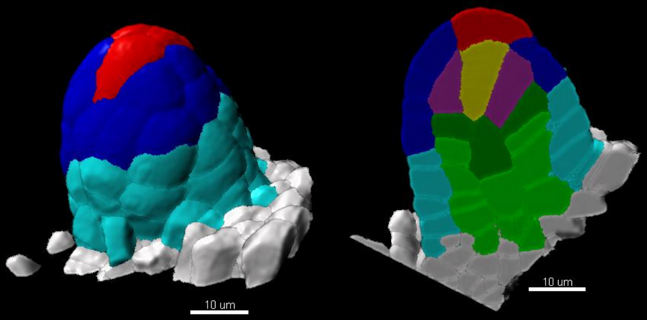


EM_C_743B

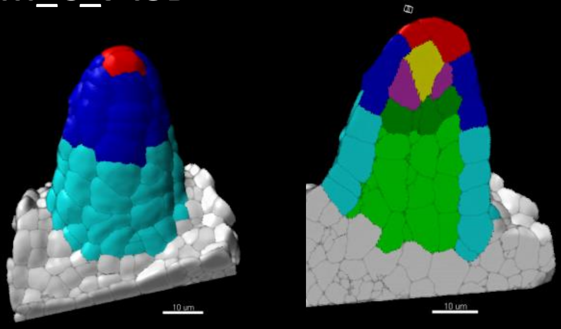


EM_C_750

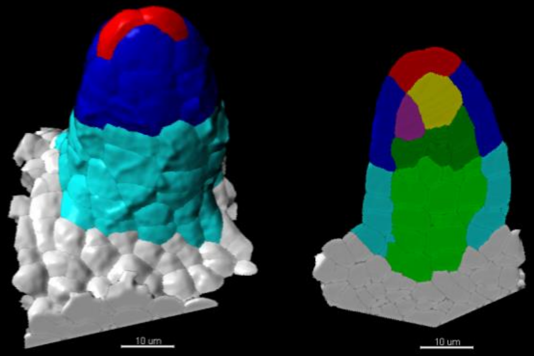


EM_C_744B

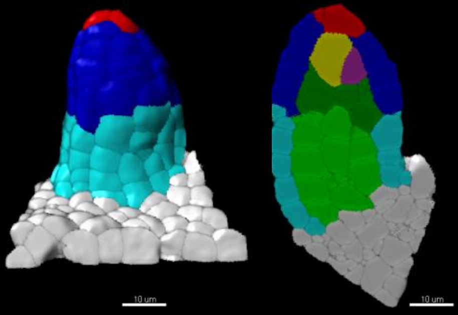


EM_C_751

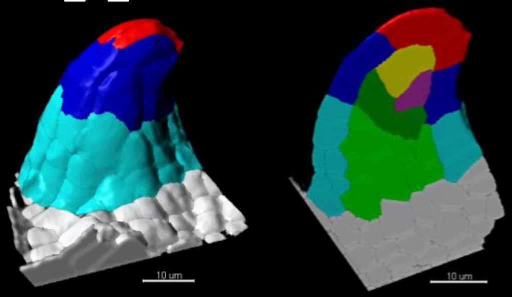


EM_C_745A

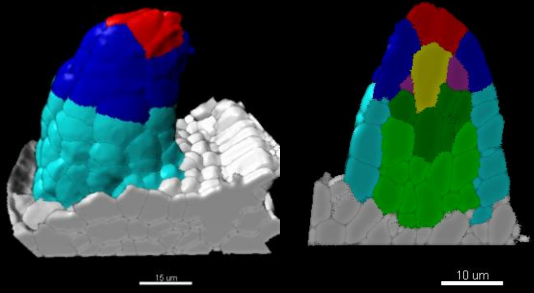


EM_C_591

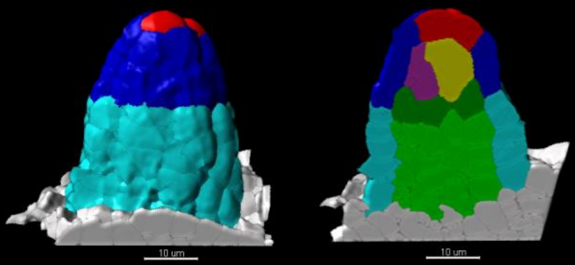

*bot1-7*

### Stage 1-II

## EM_C_755

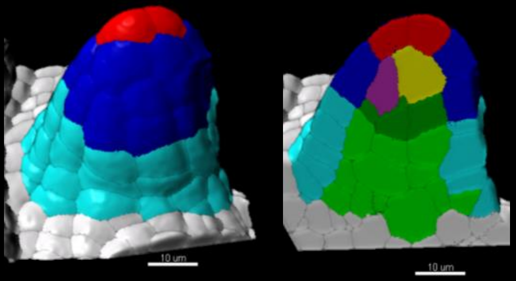


## EM_C_768

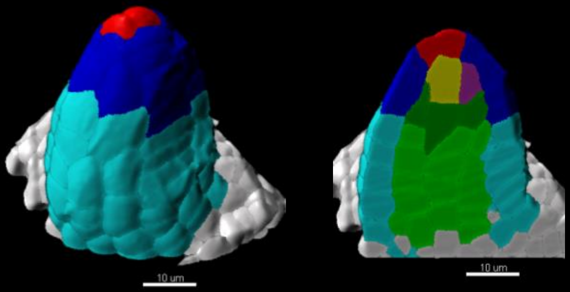


## EM_C_281B

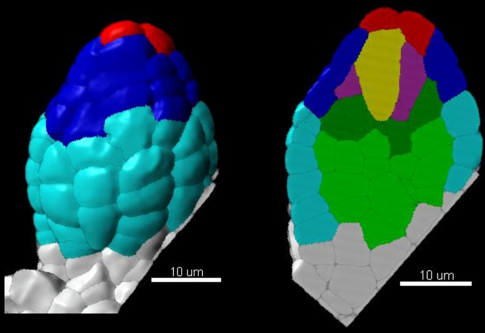


## EM_C_282B

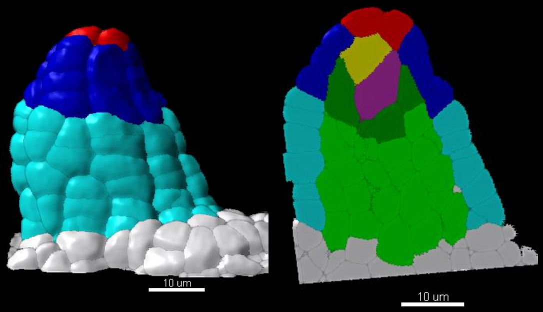

*Ws4*

### Stage 0-III

## EM_C_729B

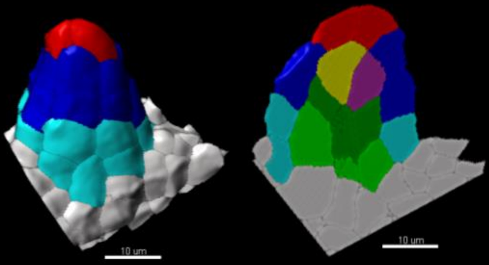

## EM_C_722

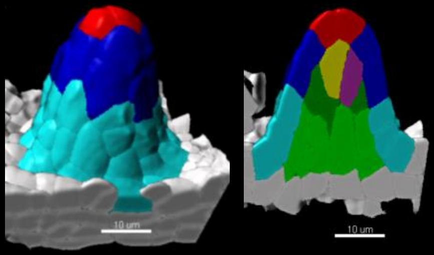

## EM_C_902B

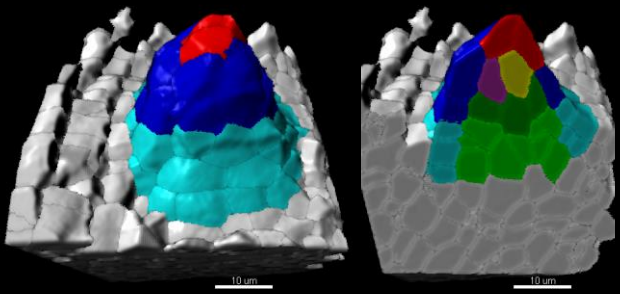

## EM_C_904

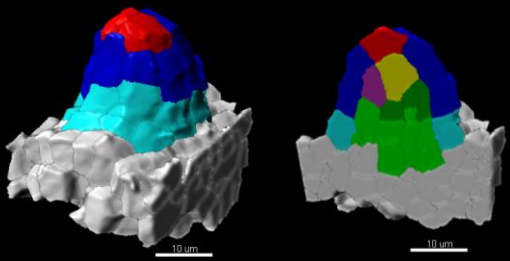

## EM_C_905A

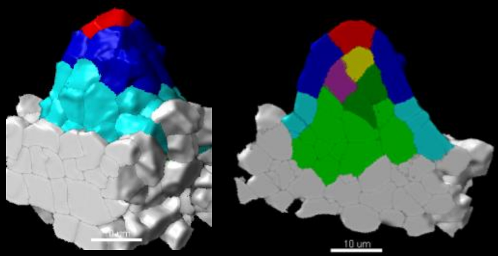

## EM_C_905B

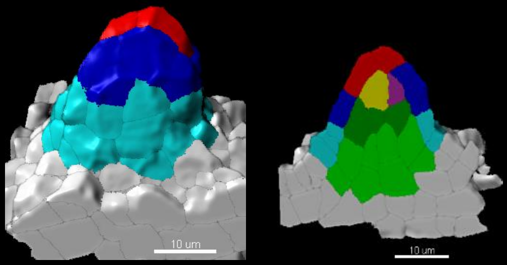

## EM_C_728B

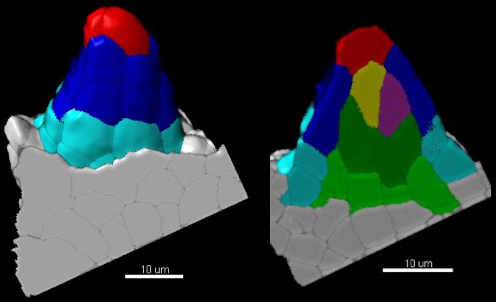

*Ws4*

### Stage 1-I

## EM_C_732

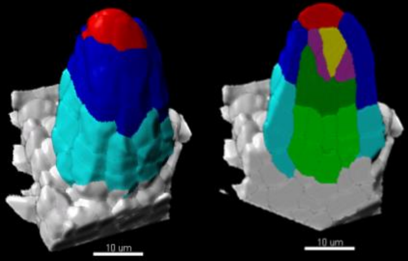


## EM_C_726

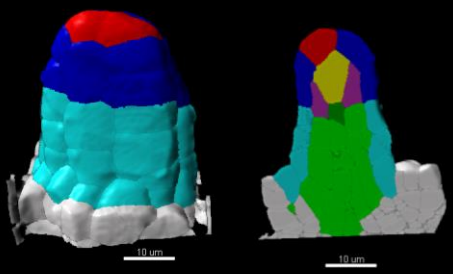


## EM_C_735A

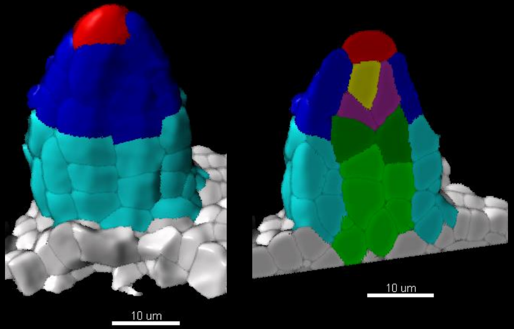


## EM_C_731A

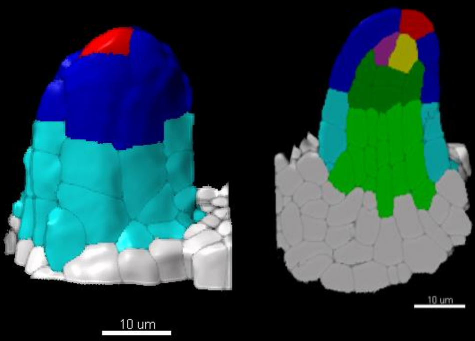


## EM_C_723

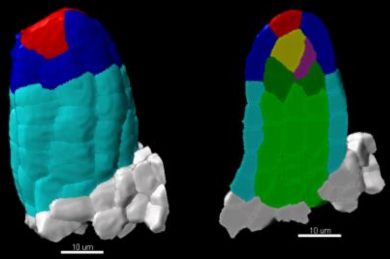


## EM_C_731B

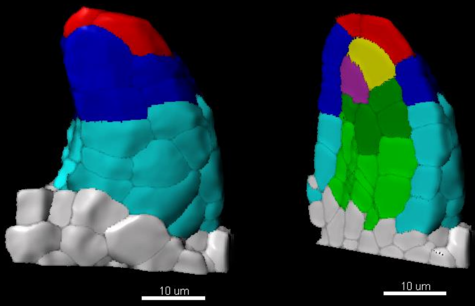


## EM_C_724

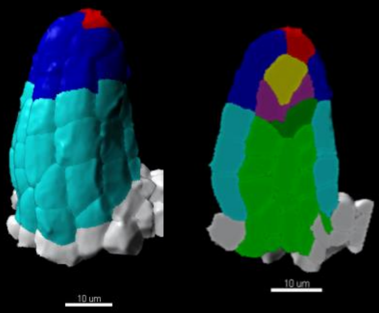


## EM_C_733

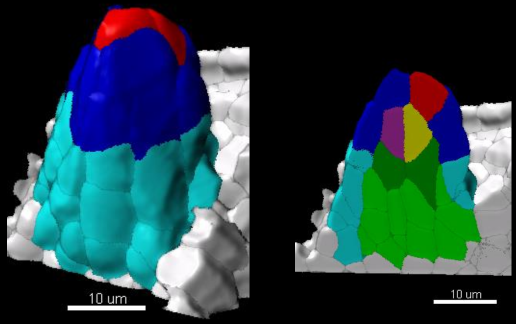

*Ws4*

### Stage 1-II

EM_C_729A

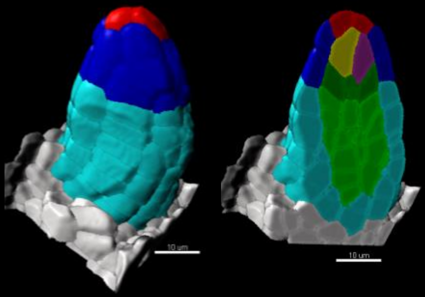

EM_C_719

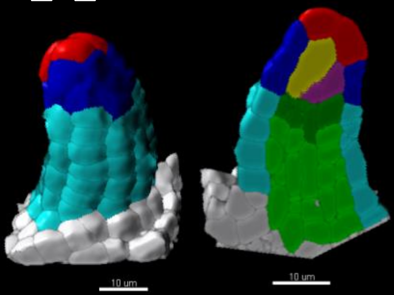

EM_C_734

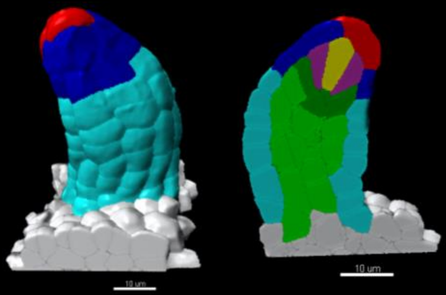

EM_C_735B

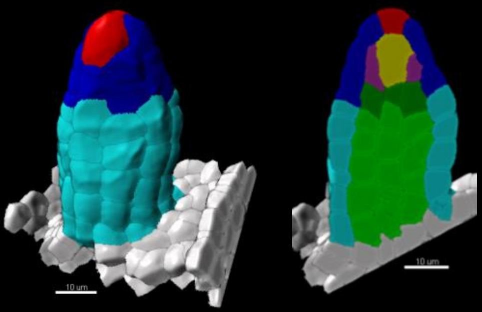

EM_C_736

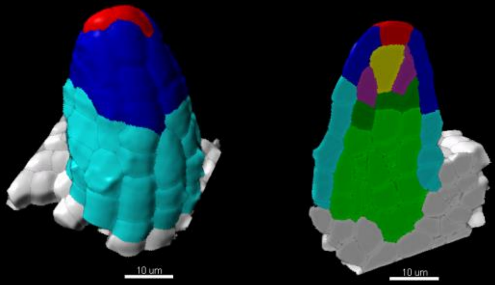

EM_C_727

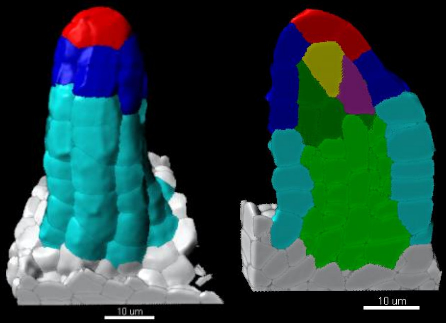

EM_C_677

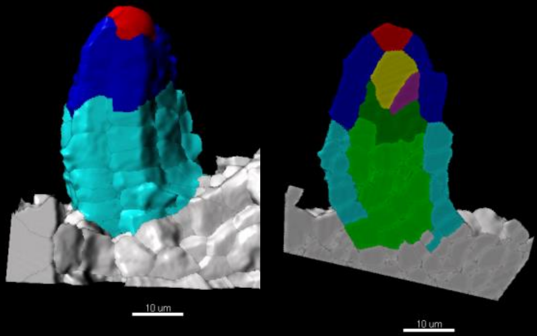
